# General cognitive function and the brain’s structural connectome

**DOI:** 10.64898/2026.03.30.715223

**Authors:** Colin R. Buchanan, Joanna E. Moodie, Venia Batziou, Eleanor L. S. Conole, Susana Muñoz Maniega, Mathew A. Harris, Hon Wah Yeung, Janie Corley, David C. Liewald, Paul Redmond, J. Douglas Steele, Gordon D. Waiter, Heather C. Whalley, Andrew M. McIntosh, Joanna M. Wardlaw, Michelle Luciano, Mark E. Bastin, Ian J. Deary, Elliot M. Tucker-Drob, Simon R. Cox

## Abstract

General cognitive function (‘*g*’) reflects a broad capacity for flexible information processing, yet how brain-wide structural connectivity supports it remains unclear. In 38,824 individuals (26–84 years) across three cohorts, we show that *g* is supported by a widely distributed white matter network. Across streamline count, fractional anisotropy (FA), and mean diffusivity (MD), meta-analytic associations with *g* were widespread at global, nodal and edge levels, spanning all cerebral lobes and key subcortical structures and dependent on both inter- and intra-hemispheric connectivity, particularly ipsilateral long-range inter-lobar connections. White matter node-*g* associations spatially mirrored independent cortical morphometry-*g* associations, indicating convergence of grey and white matter contributions. Effect sizes increased with age: MD associations became more negative and FA more positive, especially in frontal regions. Edge-level findings replicated across cohorts and predicted *g* in a hold-out sample. These findings suggest *g* reflects a distributed structural communication backbone which becomes increasingly important with age.

## Introduction

General cognitive functioning (‘*g*’) reflects the fact that, in humans, scores on diverse cognitive tests all correlate positively; *g* is an important predictor of lifelong outcomes, such as educational attainment, job performance, socioeconomic status, health, and mortality^1–3^. Individual differences in cognitive capability can be measured using a battery of tests (reasoning, memory, processing speed, visual-spatial ability, verbal ability, executive function). Latent *g* scores can be calculated by data-reduction methods such as principal component analysis^4^, reflecting the consistent positive correlation among diverse cognitive task scores, which typically accounts for approximately 40% of the variation in overall cognitive performance^5–7^. Furthermore, the relative ranking of individuals based on their *g* scores has found to be largely invariant to the test battery used^8,9^, making them particularly suited for multiple cohort studies^4^.

The prospect of cognitive decline is one of the most feared aspects of ageing, and can signal the onset of illness, dementia, and even mortality^10,11^. Cognitive decline can severely impact daily functioning and quality of life, leading to increased dependency and reduced autonomy^12,13^. Consequently, identifying the brain biomarkers associated with cognitive differences may be of value for early intervention and developing strategies to maintain cognitive health throughout the lifespan^14,15^. Exploring the biological basis of variation in *g* has practical implications for informing educational approaches, promoting healthy cognitive ageing, and advancing research into neurological conditions that impair cognitive functioning^16^.

Neuroimaging, particularly magnetic resonance imaging (MRI), has been used to explore the neural correlates of *g*, consistently finding that individual differences in cognitive abilities are associated with differences in brain structure, function, and connectivity^7^. In particular, there is a robust positive correlation (*r ≈ 0*.3) between *g* and total brain volume^17–19^. The Parieto-Frontal Integration Theory (P-FIT) has been prominent in understanding *g*’s association with brain imaging variables based on an extensive review of 37 structural, functional, and diffusion MRI (dMRI) studies^20^. P-FIT attributes intelligence differences to specific brain regions, particularly frontal, parietal, and temporal areas. However, the initial P-FIT study has limitations including small sample sizes, methodological variations, and very limited dMRI coverage^21^. Subsequent structural MRI studies have associated higher *g* scores to greater cortical volume and thickness, particularly in regions aligned with P-FIT^19,22^. However, a large-scale study discovered substantial associations (*r range: 0*.*13*–0.26) between *g* and volumes of the insula, posterior cingulate, precuneus, and thalamus^19^. These regions, absent from the original P-FIT, were also identified in a subsequent meta-analysis^23^, and reinforced by findings from cortical morphometry^24^, underscoring the evolving understanding of the neural correlates of *g*. These efforts have focussed on grey matter structures rather than the anatomical connections between them.

Previous dMRI studies have associated individual differences in *g* with the organisation of the brain’s white matter, which facilitates efficient communication between brain regions^23,25,26^. Higher *g* scores are associated with healthier white matter, specifically greater fractional anisotropy (FA), which reflects more coherent fibre organisation, and lower mean diffusivity (MD), indicating reduced overall water molecule diffusion consistent with more intact white matter microstructure^19,25–28^. These relationships are observed at both global and tract-specific levels. Certain pathways, including long-range cortico-cortical connections, the genu of the corpus callosum, and subcortico-cortical pathways have provided robust but modest associations (*r* < 0.11) with *g*^*19*^. Age-related white matter hyperintensities (WMH), often linked to cerebral small vessel disease, have been related to poorer cognitive function, especially in older age groups^29–31^. Additionally, the peak width of skeletonized mean diffusivity (PSMD) has shown to be a sensitive marker of diffuse white matter microstructural alterations related to cognitive performance^32^, but is derived from only a small proportion of total white matter. Tract-based studies have focused on a very limited number of long-range connections, which constitute only ~2% of cortico-cortical projections, overlooking the majority of local U-fibre connections^33^, which are thought to support local information integration.

Advances in neuroimaging have enabled the measurement of the connectome, which seeks to map whole-brain connectivity as a network^34^, either as a structural network of anatomical connections between brain regions^35^, or as a functional network of neural activity^36^. Functional MRI studies have found that individual differences in *g* are associated with functional connectivity throughout the brain, particularly in frontal and parietal regions^37–39^. A recent structural connectivity study predicting complex traits from all network connections reported modest correlations with *g* (r range: 0.13–0.20), with streamline count measures providing the highest prediction accuracy^40^. Structural connectivity has previously been related to visuospatial reasoning, information processing speed, and crystallised ability (i.e., learned knowledge) in 73-year-olds (β range: 0.10–0.22), with WMH volume partly mediating the connectome-processing speed relationship^41^. A related connectome study found that elements of the central executive network were particularly important for later-life cognitive function, especially for processing speed^42^.

Importantly, measures of structural connectivity differ in the biological properties they capture. Streamline count (SC) is strongly influenced by tractography sampling and macroscopic anatomical factors such as tract size and brain volume rather than reflecting direct axonal counts^43–45^. In contrast, diffusion tensor metrics such as FA and MD show only weak associations with volumetric measures and reflect microstructural features of white matter such as axonal organisation and myelination^46,47^. Considering these complementary network weightings together allows us to assess how cognitive differences are supported by brain size-linked wiring capacity and microstructural organisation, capturing both volume-related and volume-independent aspects of connectivity.

The precise relationship between *g* and brain connectivity remains an active area of research. Much work to-date has focused on selective pathways in small samples, making the robustness of these associations uncertain. Recent evidence suggests that small sample sizes in brain-behaviour studies can inflate effect sizes and limit replicability, with reliable brain-wide associations often requiring thousands of participants^48,49^. Consequently, robust characterisation of connectome-*g* relationships requires both large cohorts and independent replication, which are critical gaps in the current literature. There is a need for well-powered meta-analytic approaches at the network level, which would enable robust mapping of *g*’s correlations across thousands of white matter pathways with unprecedented detail.

Here, we conducted the most comprehensive analysis to-date of the relationship between cognitive functioning and structural brain connectivity. By applying connectome construction and meta-analysis across three cohorts, we examined how macroscopic wiring capacity, microstructural organisation, and age-related differences contribute to individual differences in *g*. We standardised macroscale network construction across scanners, estimated *g* from multiple cognitive tests within each cohort, and tested associations between *g* and network properties while accounting for age, sex, and scanner effects. We then meta-analysed and assessed cross-cohort consistency of brain-*g* associations across global networks, network nodes, and individual network connections. Finally, we examined correlations between network-*g* and cortical morphometry-*g* associations, tested age moderation, and evaluated replicability by predicting *g* in a large hold-out sample using meta-analytic estimates.

## Results

Using all available brain scans, we applied whole-brain tractography and standardised connectome processing to map the brain’s axonal wiring as a network of anatomical connections at the macroscale^50^. Whole-brain networks were constructed for 38,824 participants. The largest sample was from the UK Biobank (UKB, n = 37,284, 45–83 years)^51^, randomly split into a main (n = 18,642, 45–82 years), and hold-out replication sample (n = 18,642, 45–83 years). Additional cohorts were Generation Scotland: Stratifying Resilience and Depression Longitudinally (STRADL, *n = 937, 26*–84 years)^52^, and the Lothian Birth Cohort 1936 (LBC1936, *n = 603*, ~73 years)^53,54^. Structural networks were constructed using 85 neuroanatomical regions (network nodes) segmented from T1-weighted volumes^55–57^, with white matter connections (edges) determined by dMRI and probabilistic tractography^58–60^. Individual differences in node connectivity reflect how strongly regions are connected to the rest of the brain, while edges capture variation in the presence and strength of specific white matter connections. For each participant, three network types were computed to capture different aspects of connectivity: streamline count (SC), fractional anisotropy (FA), and mean diffusivity (MD), representing the total number of streamlines and mean diffusion values along streamlines between node pairs^50^. Conceptually, SC indexes the number of macroscale connections between a pair of regions, FA indicates how well-organised or coherent those connections are, and MD reflects how easily water diffuses within them, as a marker of microstructural integrity.

### A new reference brain network for cross-cohort comparison

Overall, network organisation was comparable across cohorts, although expected scanner-related differences were observed in network density and weight distributions (Supplementary Fig. 1)^61,62^. To minimise spurious connections and to ensure a uniform network density, we applied a two-step thresholding procedure informed by findings from a prior single-scanner study^50^. Firstly, consistency-thresholding^63^ was conducted independently in each cohort at 30% network density, resulting in 1071 network edges. Secondly, we intersected the resulting network masks from the three cohorts, each containing potentially distinct edges, and retained only the edges present in all three cohorts. Agreement between cohort pairs was high: UKB and STRADL shared 943 edges (88.0%), STRADL and LBC1936 shared 898 edges (83.8%), and UKB and LBC1936 shared 850 edges (79.4%). Overall, the shared edges were highly consistent across all three cohorts and resulted in a network of 818 edges (76.4% of 1071 thresholded connections).

The resultant macroscale connectome (Fig. 1) reflected a hierarchical and integrative topology, comprising 647 intra-hemispheric connections (79.1% of total connections) and 166 inter-hemispheric connections (20.3%), with a near-symmetric distribution between hemispheres (left: 328; right: 319). Total inter-lobar connections (558) were roughly double the number of intra-lobar connections (230), indicating a predominantly integrative and ipsilateral network organisation at this scale. The parietal, temporal, and frontal lobes contributed most to inter-lobar connectivity (241, 224, and 212 connections, respectively), while the frontal lobe exhibited the highest intra-lobar connectivity (83). Subcortical regions exhibited both substantial intra-lobar connections (51) and the highest inter-lobar connectivity (279), highlighting their central role in overall network integration. This 818-edge network was used for all subsequent analyses and is provided as a definitive multi-cohort macro-connectivity network for connectome research in the Supplementary Information.

**Fig. 1:**
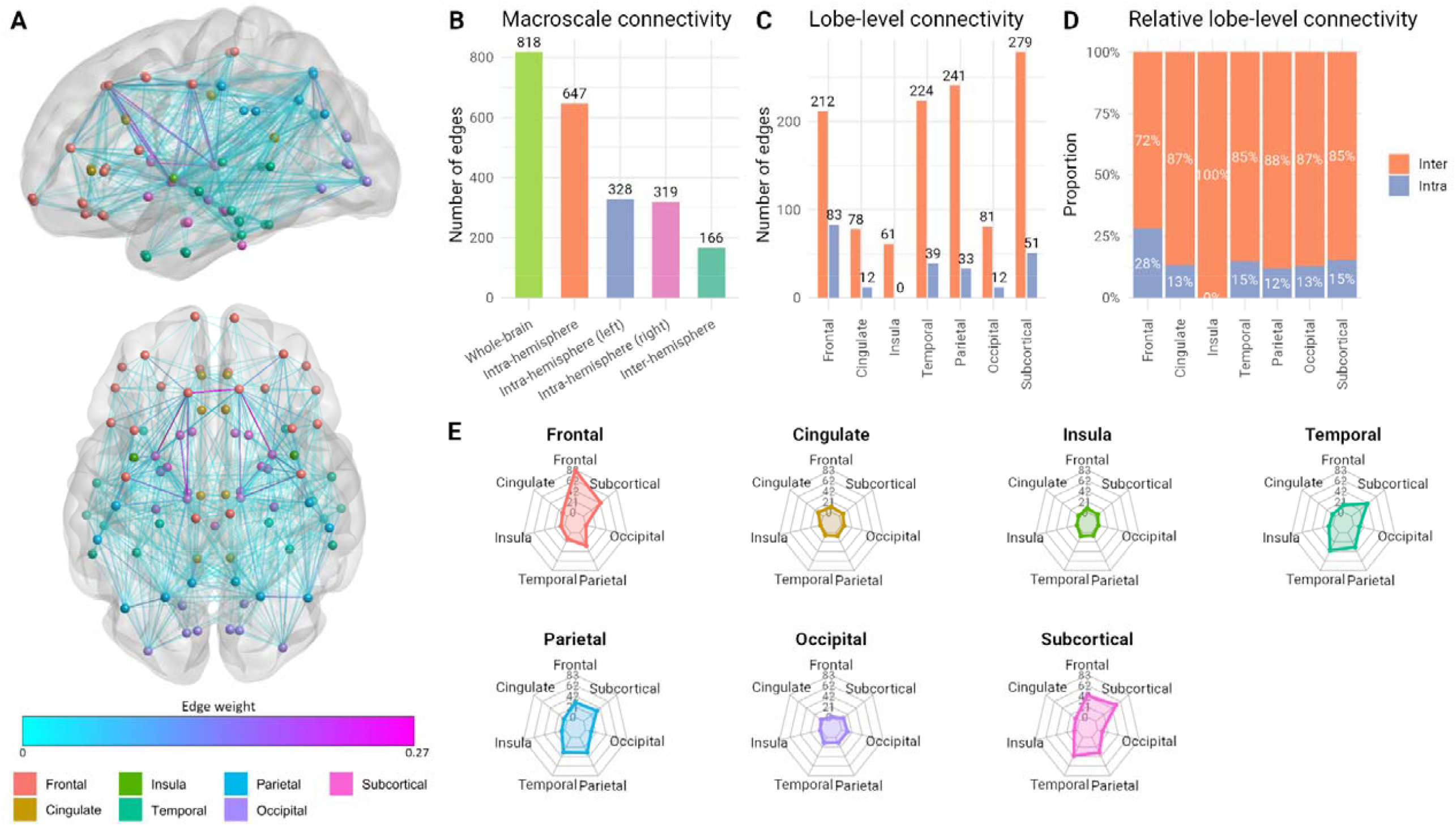
A robust, cross-cohort macroscale structural connectome. A) Anatomical plot (818 edges) where link colour indicates mean streamline count across participants (scaled), and nodes are coloured by lobe. B) Macroscale connectivity edge counts. C) Lobe-level connectivity edge counts, comparing inter- and intra-lobe connections. D) Relative lobe-level connectivity. E) Radar plots showing the proportion of edges involved in lobe-to-lobe connectivity.

### Global brain networks and general cognitive functioning

A latent factor of *g* was estimated for each cohort using diverse cognitive tests assessing multiple cognitive domains^52,64,65^, and confirmatory factor analysis within a structural equation modelling (SEM) framework. All models provided acceptable fit (CFI > 0.94; TLI > 0.92; RMSEA < 0.06; SRMR < 0.05), with factor loadings on *g* predominantly moderate-to-high across tests (0.28–0.73; Supplementary Tables 1-4). These *g* scores showed consistent correlations with age and education across cohorts (Supplementary Table 5). Brain network associations with *g* (standardised βs) were computed using structural regression in SEM, adjusting for age, sex, and scanner (scanner applies to UKB and STRADL). For each network, three global graph-theoretic metrics were calculated using weighted metrics^66^: mean edge weight, global network efficiency, and mean clustering coefficient. Mean edge weight quantifies the overall strength of connection across the network. Global efficiency reflects how effectively information can be integrated across the brain, with higher efficiency indicating a more optimally connected network. Clustering coefficient captures the tendency of regions to form local clusters that support segregated processing, with higher values suggesting more specialised and modular network organisation.

Across all three cohorts, global network analyses (Fig. 2) revealed positive associations with *g* for SC network metrics (median β = 0.18, range: 0.12 to 0.28), and for FA metrics (median β = 0.13, range: 0.06 to 0.30). As expected, all MD metrics were negatively associated with *g* (median β = −0.08, range: −0.22 to −0.01). CIs were narrower in UKB, reflecting its larger sample size compared with the other cohorts. Across cohorts, SC metrics showed more consistent associations (β ≈ 0.2) than for FA and MD metrics. For FA networks, UKB provided considerably lower estimates (β < 0.1) than STRADL, although LBC1936 confidence intervals overlapped with both other cohorts. For MD networks, although the confidence intervals overlapped, both UKB and LBC1936 provided lower estimates (|β| < ~0.1) than STRADL (β ≈ 0.2), and LBC1936 associations were non-significant. A three-way meta-analysis was then performed using a random effects model to calculate meta-analytic coefficients from the cohort-specific network-*g* associations. The meta-analytic βs were significant for all global metrics tested (*p* < 0.05, FDR; Supplementary Table 6).

**Fig. 2:**
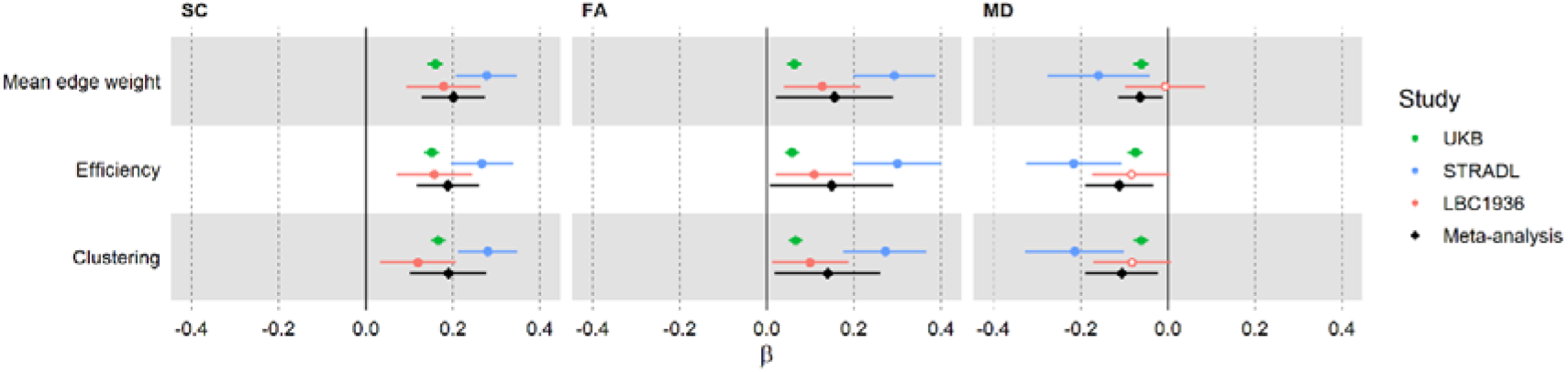
Associations between general cognitive function (‘*g*’) and global network organisation. Forest plots of standardised regression coefficients (βs) for associations between *g* and global network metrics. Associations are shown across three network weightings, three cohorts, and the meta-analysis. Solid markers indicate significance (*p* < 0.05, FDR), with error bars representing standard errors. Network weightings are streamline count (SC), fractional anisotropy (FA), and mean diffusivity (MD).

Our main analyses included all available participants. A supplementary sensitivity analysis excluding participants with neurological conditions (e.g., dementia, stroke; see Supplementary Information) showed an attenuation of *g* associations (0.8–7.3% in UKB; 2.1–28.2% in LBC1936). However, because neurological exclusion criteria were unavailable for STRADL and differed across cohorts, all subsequent analyses used the full samples, which also preserved statistical power in the smaller cohorts.

### Regional node-*g* associations

We localised our analyses by computing weighted local efficiency, which reflects each node’s capacity for information transfer within its immediate neighbourhood^66^. Across all three cohorts, associations between *g* and local efficiency were mainly positive for SC and FA, and negative for MD (Fig. 3; Supplementary Tables 7–10). STRADL showed the strongest associations (mean β: 0.15 for SC, 0.18 for FA, –0.18 for MD), whereas associations were weaker in UKB (mean β: 0.09 for SC, 0.05 for FA, –0.05 for MD) and LBC1936 (mean β: 0.06 for SC, 0.07 for FA, –0.08 for MD). A fuller summary, including medians and ranges, is provided in Supplementary Table 11.

**Fig. 3:**
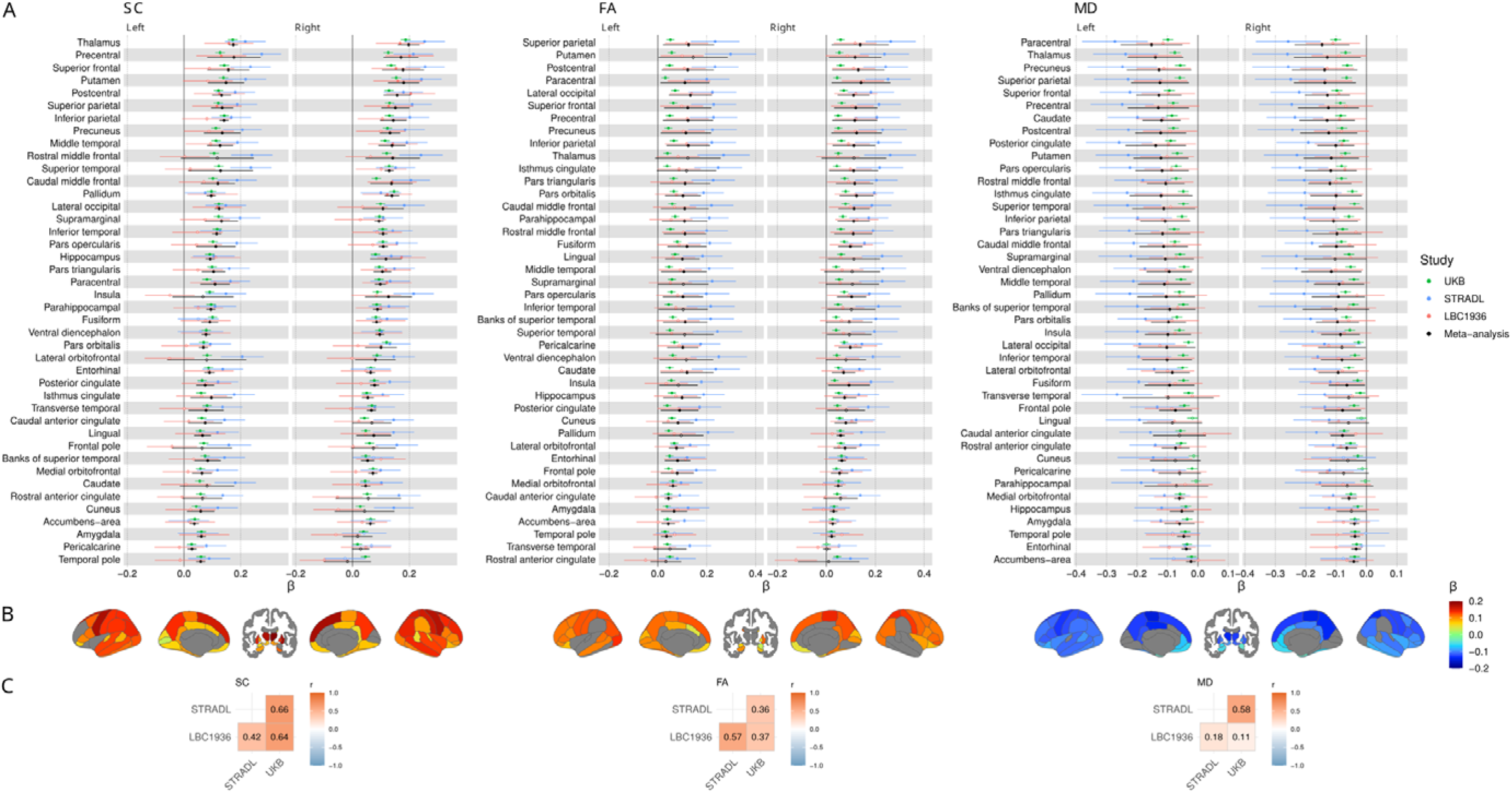
Regional associations between *g* and local efficiency. Standardised regression coefficients (β) for associations between *g* and local efficiency in streamline count (SC), fractional anisotropy (FA), and mean diffusivity (MD) networks, shown across three cohorts and the meta-analysis. A) Forest plots of nodal associations with nodes ordered by the average magnitude of the meta-analytic βs across hemispheres, and the brainstem is not plotted. Solid markers indicate significance (*p* < 0.05, FDR), with error bars representing standard errors. B) Anatomical plots of significant nodes, where colour represents the meta-analytic β. C) Correlations of β coefficients between cohorts.

The meta-analysis of local efficiency across 85 network nodes (Fig. 3) revealed widespread significant *g* associations involving both cerebral hemispheres, all lobes, and some subcortical areas: 72/85 nodes (84.7%) for SC, 62/85 (72.9%) for FA, and 70/85 (82.4%) for MD (*p* < 0.05, FDR). The strongest meta-analytic node-*g* associations (|β| > 0.1) were observed across all brain lobes, forming a distributed frontotemporoparietal-subcortical network, with some strong associations in occipital, cingulate, and insular regions. Notably, almost all parietal nodes showed strong associations across SC, FA, and MD networks. By calculating the correlation between all left and right hemisphere node associations, we found that these meta-analytic associations were highly symmetrical (*r = 0*.*84 for* SC, *r = 0*.*86 for* FA, *r = 0*.*88 for* MD; *p* < 0.001; Supplementary Table 12).

To evaluate cross-cohort agreement, we calculated correlations between the set of node βs for each of the three possible cohort pairs (Fig. 3C). Despite differences in the absolute values of node βs across cohorts (mean |β|: ~0.17 in STRADL, ~0.07 in LBC1936, ~0.06 in UKB), these correlations were weak-to-strong (*r range: 0*.*11* to 0.66, Supplementary Fig. 2), although strongest for SC (*r ≥ 0*.42) and FA (*r* ≥ 0.37), and lowest for MD (r ≥ 0.11). However, some FA estimates varied to the extent that substantial meta-analytic βs were rendered non-significant, such as for the thalamus (β < 0.13, *p* > 0.07). Cross-cohort differences were more pronounced for MD networks (*r range: 0*.*11* to 0.58), with STRADL and UKB showing moderate agreement (*r = 0*.*58), bu*t weaker agreement for LBC1936 (*r ≤ 0*.18). While UKB and LBC1936 provided comparable node-*g* magnitudes for both FA and MD networks, in LBC1936 all nodal estimates were non-significant after FDR correction, likely reflecting reduced statistical power due to the smaller sample size. The direction of node-*g* associations was consistent across cohorts for most nodes: 72/85 (84.7%) for SC, 76/85 (89.4%) for FA, and 82/85 (96.5%) for MD.

Our main analyses used uncorrected SC, which was strongly correlated with total brain volume (TBV; *r = 0*.*80, p* < 0.001). In contrast, FA and MD networks showed only weak correlations with TBV (|*r*| < 0.10, *p* < 0.001). Given the strong SC-TBV correlation, we conducted a supplementary analysis using streamline density (SD), which applies regional volumetric correction to SC values, to assess SC-specific contributions. Overall, SD provided weaker node-*g* associations and fewer significant nodes than SC, and the meta-analytic associations were lower for SD (mean β = 0.03; range = –0.06 to 0.11) than for SC (mean β = 0.10; range = –0.02 to 0.20; Supplementary Table 8). This attenuation may indicate that a substantial component of SC-*g* associations reflects volumetrically-dependent macroscopic wiring capacity, rather than measurement artefact. However, in line with prior work noting limitations of SD as a correction approach^44,50^, we report uncorrected SC in subsequent analyses, with results interpreted in light of the strong SC-TBV correlation.

### Network edge-*g* associations

We further localised our analyses to the 818 individual network connections (edges) identified from the cross-cohort reference connectome. Rather than being confined to a small set of specialised pathways, *g*-related associations were widely distributed across the structural connectome (Fig. 4), consistent with the view that general cognitive performance reflects the efficiency and robustness of distributed communication across brain systems.

**Fig. 4:**
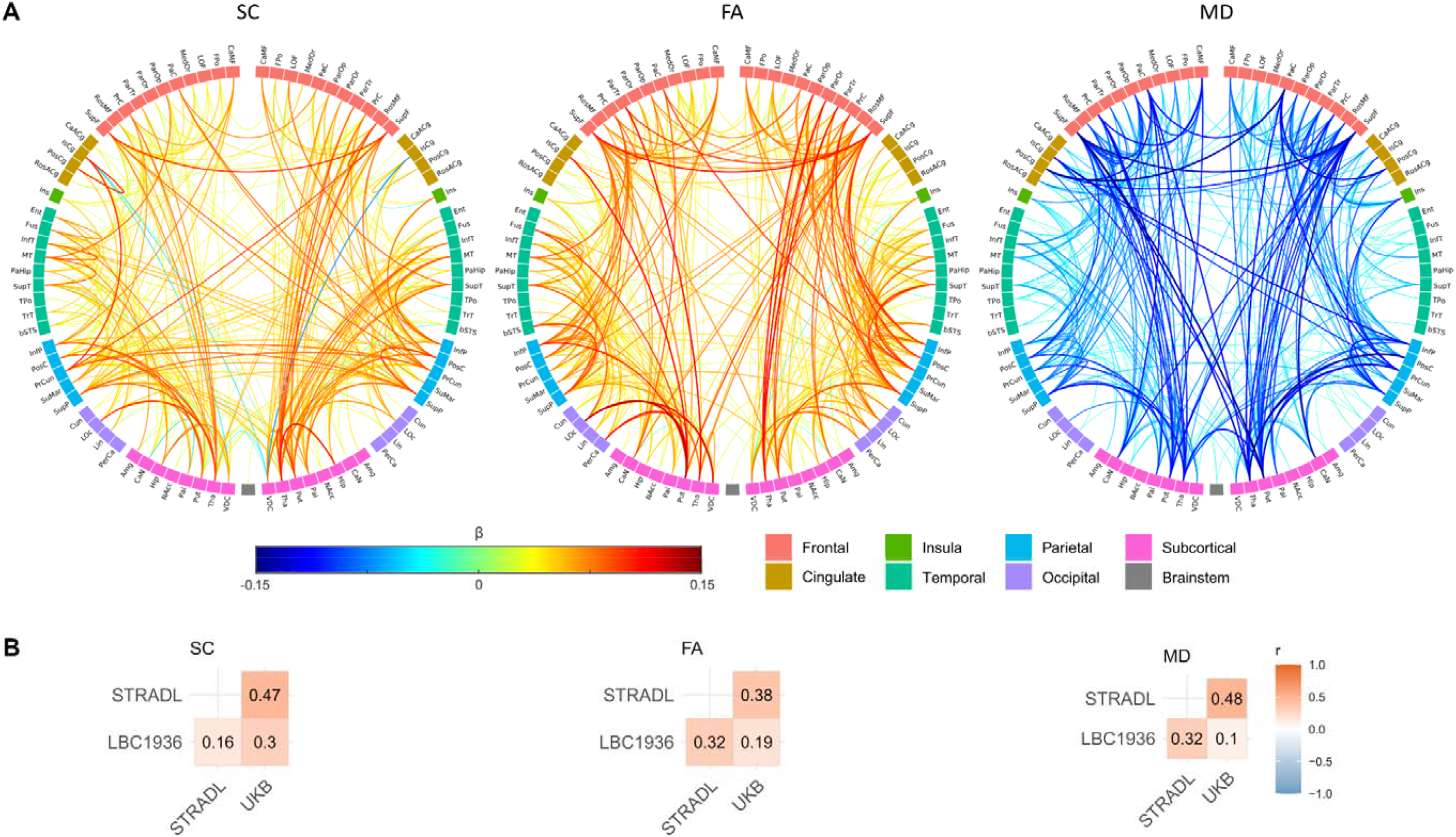
Edge-level associations between *g* and white matter connections. Significant edge-*g* associations (*p* < 0.05, FDR) from separate meta-analyses of streamline count (SC), fractional anisotropy (FA), and mean diffusivity (MD) networks. A) Significant associations, where link colour represents standardised β coefficients and link thickness reflects the magnitude of the association. The number of significant edges was 391 for SC, 433 for FA, and 325 for MD. Node abbreviations are listed in Supplementary Table 13. B) Correlation of β coefficients between cohorts (cf. Fig. 3 for node-level correlations).

Consistent with global and nodal analyses, associations between *g* and edge weights were mainly positive for SC (mean β = 0.030, median = 0.030, range: −0.152 to 0.229) and FA (mean β = 0.052, median = 0.046, range: −0.146 to 0.240), and negative for MD (mean β = −0.046, median = −0.039, range: −0.278 to 0.162). Effect sizes varied across cohorts and were generally smaller in LBC1936 and UKB than in STRADL (Supplementary Fig. 3; Supplementary Table 14). In particular, STRADL provided stronger and broader MD associations (mean β = −0.078, median = −0.076, range: −0.278 to 0.124) than UKB (mean β = −0.038, median = −0.036, range: −0.104 to 0.024) or LBC1936 (mean β = −0.021, median = −0.023, range: −0.219 to 0.162), which may partly reflect the broader age range in the STRADL cohort. After FDR correction, few edges were significant in LBC1936 (16 for SC, 4 for FA, 7 for MD; *p* < 0.05, FDR), likely reflecting limited statistical power, although spatial patterns were visually similar across cohorts (Supplementary Fig. 3). We quantified cross-cohort agreement by the correlations between edge-level β estimates across cohort pairs (Fig. 4B; Supplementary Fig. 4). Agreement was moderate (*r range: 0*.*10*–0.48) and lower than at the node level (*r range: 0*.*11*–0.66; Fig. 3C).

Meta-analysis revealed significant edge-*g* associations for 39.7–52.9% of 818 edges (*p* < 0.05, FDR), distributed widely across all lobes and broadly symmetrical across hemispheres (Fig. 4). Significant edges comprised 47.8% (391/818) of edges in SC networks, 52.9% (433/818) in FA, and 39.7% (325/818) in MD. A moderate overlap was observed among the significant edges across the three network weightings (SC, FA, and MD), with 369 edges shared by at least two weightings but only 94 common to all three weightings. This highlights the distinct aspects of connectivity captured by these dMRI-based measures, particularly for MD.

Significant *g*-related associations were more prevalent for intra-hemispheric than inter-hemispheric connections, with a ratio exceeding 4:1 across network weightings (Supplementary Table 15). Inter-hemispheric associations were most pronounced between frontal and parietal regions, but inter-hemispheric connectivity was more widespread in MD than in SC or FA networks (Fig. 4; Supplementary Fig. 5). Additionally, significant inter-lobe connections were more numerous than intra-lobe connections and were predominantly ipsilateral (Fig 4; Supplementary Table 16). The strongest associations (|β| > 0.1) were found for the precuneus and insula, which are considered integral to network communication. Notably, strong subcortico-cortical associations included the thalamus, caudate, putamen and hippocampus (Fig. 4).

**Fig. 5:**
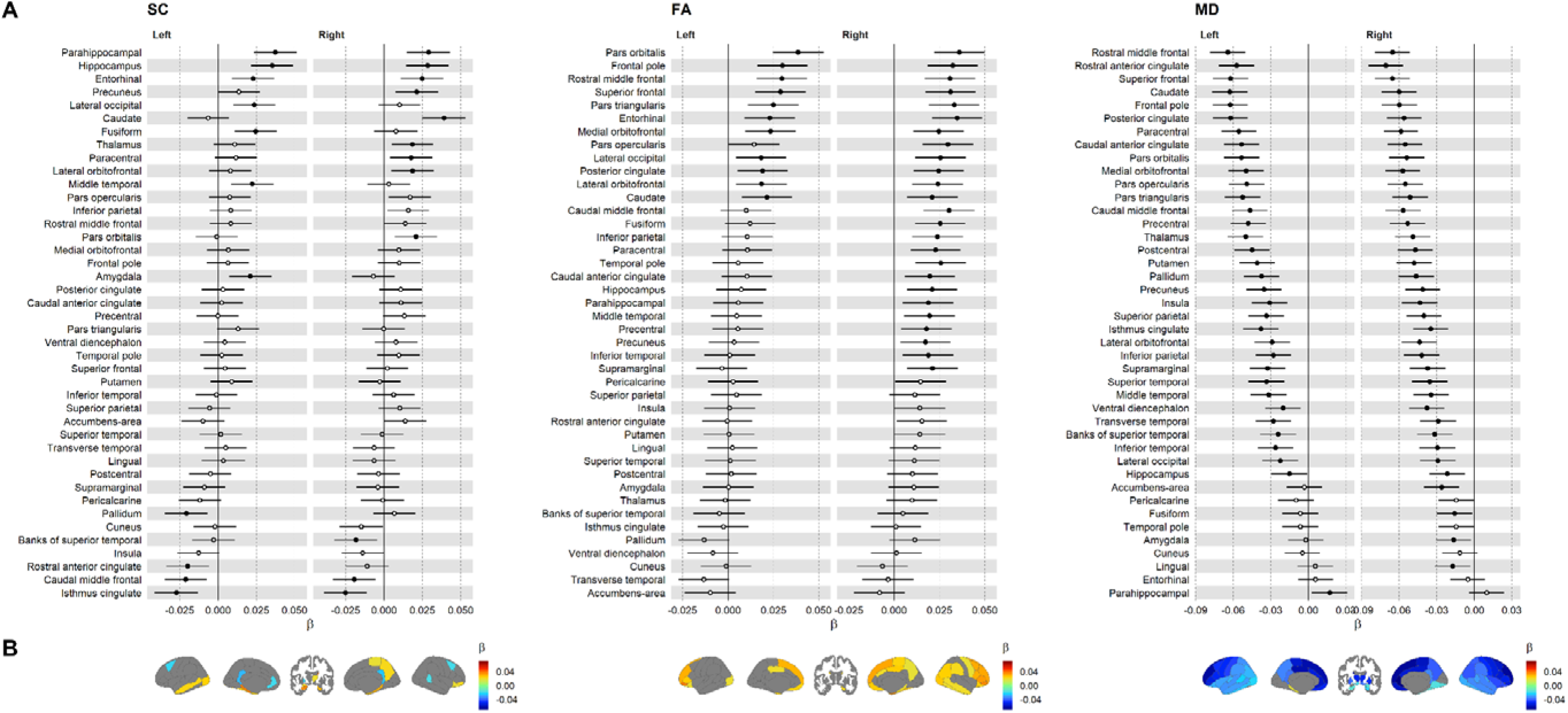
Age moderation of node-*g* associations. A) Forest plots of age × node interactions (standardised β coefficients showing per SD increase in age) for local efficiency in streamline count (SC), fractional anisotropy (FA), and mean diffusivity (MD) networks (UK Biobank data only). Nodes are ordered by average β magnitude across hemispheres, and the brainstem is not shown. Solid markers indicate significance (*p* < 0.05, FDR), with error bars representing standard errors. B) Anatomical plots of significant nodes, with colour indicating the magnitude and direction of age moderation.

### Comparison of node-*g* and cortical morphometry-*g* associations

Additionally, we assessed the relationship between network node-*g* associations from this study and corresponding *g* associations of cortical morphometry reported in a prior analysis of the same cohorts^24^. This approach allows direct quantitative comparison between regional white matter connectivity and cortical grey matter properties in the same atlas. Specifically, correlations were calculated between the meta-analytic β coefficients from our node-*g* associations and the β coefficients from the structural MRI-based cortex-*g* associations. Correlations were calculated across 68 cortical regions from the Desikan-Killiany atlas^55^, for all nine pairings of network weightings (SC, FA, and MD) and cortical measures (cortical volume, surface area, and thickness). Despite the conceptual distinction between grey and white matter measurements, correlations between node-*g* and cortex-g coefficients ranged from weak to strong (|*r*| *ra*nge: 0.12–0.65; Supplementary Table 17). As expected, SC and FA measures showed positive correlations, and MD showed negative correlations. Across all three network weightings, cortical volume showed the strongest correlations with node-*g* (|*r*| = 0.41–0.65), followed by surface area (|*r*| *=* 0.35–0.56) and cortical thickness (|*r*| *=* 0.12–0.33). All correlations were statistically significant (*p* < 0.05), except for the thickness-MD correlation. Scatter plots coloured by lobe did not reveal any particular regional drivers of these relationships (Supplementary Fig. 6).

### Age moderation of network-*g*

Age moderation analyses were conducted for UKB (45–81 years) and STRADL (26–84 years) for global and nodal properties (Fig. 5 and Supplementary Tables 18–21). No significant age moderation effects for network-*g* associations were observed in STRADL after FDR correction (all |β| < 0.12), likely reflecting limited statistical power. In the larger UKB cohort, significant age moderation effects were observed at both global and node levels.

Five of nine global network metrics showed significant age moderation (*p* < 0.05, FDR). All three MD metrics showed negative age moderation effects (age × network interaction β = –0.047 to –0.041; *p* < 0.001), indicating that negative associations between MD network properties and *g* became stronger with increasing age. Conditional standardised slopes estimated from the interaction models confirmed this age-related strengthening. For example, for mean edge weight the estimated MD-*g* association increased in magnitude from β = −0.000 at 56.1 years (−1 SD age), to β = −0.042 at the mean age (63.7 years), and β = −0.083 at 71.4 years (+1 SD age). Two FA metrics (mean edge weight and clustering coefficient) showed positive age moderation effects (interaction β ≈ 0.017; *p* < 0.05), whereas global efficiency did not reach significance (interaction β = 0.013). These effects indicate that positive FA network-*g* associations became stronger with increasing age, although the magnitude of FA moderation effects was smaller than observed for MD. For example, for mean edge weight, conditional standardised slopes from the interaction model indicated an increase in the FA-*g* association from β = 0.036 at −1 SD age, to β = 0.052 at the mean age, and β = 0.069 at +1 SD age.

At the nodal level, similar patterns were observed for local efficiency (Fig. 5). Significant age moderation effects were identified in 72/85 nodes in MD networks (all β ≥ −0.071), 37/85 nodes in FA networks (all β ≤ 0.039), and 23/85 nodes in SC networks (all β ≤ 0.039). Among these, 71/72 nodes in MD networks showed negative moderation, and all 37 significant nodes in FA networks showed positive moderation, consistent with stronger node-*g* associations at older ages. Notably, the magnitude of age moderation in MD networks appeared to follow an anterior-posterior gradient, with the strongest effects in frontal regions, but this gradient was less pronounced in FA networks and absent in SC networks.

In UKB, we also computed spatial correlations of age-moderation effects between dMRI weightings. Correlations were moderate for FA-MD (*r = −0*.401) and SC-FA (*r = 0*.*346), b*ut weak for SC-MD (*r = 0*.*149), i*ndicating partial spatial correspondence while suggesting that these network weightings reflect distinct, age-sensitive aspects of connectivity and microstructure.

### Replication of network-*g* associations in a hold-out sample

To assess how well network connectivity predicts g, we calculated composite scores by weighting each of the 818 network edges by its meta-analytic β coefficient and summing across edges. Correlations between these predicted scores and the observed g values were calculated both within the training samples and in the unseen UKB hold-out sample (Supplementary Table 22). Within the training samples (UKB, STRADL, and LBC1936) these correlations ranged from r = 0.08 to 0.26. Notably, predictions remained stable between the UKB main (r range: 0.14 to 0.26) and hold-out samples (r range: 0.13 to 0.26). SC networks showed the strongest correlations in UKB and STRADL (r ≈ 0.26) but were lower in LBC1936 (r = 0.19). FA correlations were consistent across samples (r range: 0.13 to 0.17). MD correlations were comparable between UKB samples (r ≈ 0.18) but lower in STRADL (r = 0.13) and LBC1936 (r = 0.08). Notably, for UKB and LBC1936, predictions based on edge-wise composite scores (r range: 0.08 to 0.26) exceeded the associations between g and global network metrics across network weightings (|β|range: 0.01 to 0.18; Fig. 2), demonstrating that edge-level connectivity was more informative than summarising whole-brain networks with a single global metric.

## Discussion

This study, involving nearly 39,000 participants across three cohorts, provides the most extensive and well-powered analysis to date of how brain connectivity relates to general cognitive function, a cardinal human behavioural trait. In addition, we established a robust reference brain network of the most consistently identified white matter connections across three different cohorts. This approach reduces false-positive connections while ensuring an anatomically plausible network density at the macroscale^50,63^. We share this resource with the research community to support future connectome analyses.

Our mapping of network-g associations advances understanding of the relationship between general cognitive function (g) and brain structure. Consistent with prior dMRI studies, we observed predominantly positive g associations with fractional anisotropy (FA), while associations with mean diffusivity (MD) were mainly negative^19,25,26,28^. Network-based streamline counts (SC), which index the number of fibre connections between regions, showed primarily positive associations with g and provided the strongest and most replicable effects. The strongest network-g associations were observed for SC and FA networks, where some β values approached ~0.3, and are of comparable magnitude with some of the most robust brain-g associations reported in the literature, such as those involving brain size^17–19^. Our findings suggest that associations between brain size and cognitive function may be partly instantiated through the macroscopic organisation of white matter connectivity. Streamline count networks (which are strongly correlated with total brain volume) showed the largest and most replicable associations with g, consistent with the idea that individuals with larger brains possess a greater number of long-range anatomical connections that support distributed information integration^67^. In this view, SC represents a structural pathway through which volumetric differences translate into functional cognitive capacity. At the same time, the presence of widespread g associations in FA and MD networks (which were largely independent of brain size) identifies aspects of white matter connectivity that contribute meaningfully to cognitive variation beyond gross anatomical scaling^46^. Together, these findings quantify robustly how general cognitive function is supported by both the quantity and quality of large-scale brain wiring.

We found that meta-analytic network-g associations for nodes (mean |β| = 0.03–0.10) and edges (mean |β| = 0.03–0.05) were modest but comparable in magnitude to prior tract-g associations reported in UKB (|β| < 0.11)^19^, and similar to meta-analytic cortical morphometry-g associations (mean β = 0.10–0.30) observed in the same cohorts^24^. Our regional meta-analysis found widespread significant nodal associations of g across all cerebral lobes and several subcortical regions: 73-85% of all network nodes had significant associations. The strongest associations (|β| > 0.1) formed a frontotemporoparietal-subcortical network, with multiple brain regions contributing to cognitive function, highlighting the distributed nature of g. This widespread distribution of significant edge-g associations suggests that general cognitive function is supported less by isolated ‘critical’ connections than by the collective integrity of a large-scale communication backbone^44,67^. Such a distributed architecture may support cognitive performance by enabling flexible routing of information, redundancy against local inefficiencies, and efficient integration across functionally diverse regions. From a behavioural perspective, this implies that individual differences in g reflect differences in the overall capacity for coordinated information processing rather than reliance on specific cognitive modules^7^. The symmetrical distribution of node-g associations across cerebral hemispheres supports the idea that cognitive functions often depend on the coordinated activity of both hemispheres through callosal fibres^68^.

Our meta-analysis of network connections (edges), found that 40-53% of all edges had significant edge-g associations, which again were widespread across all lobes, and were broadly symmetrical across hemispheres. While the magnitude of the edge associations varied across cohorts, the overall pattern of associations was relatively consistent. Our analyses at the edge level revealed that significant intra-hemisphere connections considerably outnumbered significant inter-hemisphere connections. However, significant intra-lobe connections were fewer than significant inter-lobe connections, which were predominantly ipsilateral rather than contralateral. This suggests that, at the macroscopic network level, *g* is supported more by ipsilateral, long-range, inter-lobe connectivity than by local, within-lobe connectivity. However, strong associations for inter-hemispheric connections were observed, particularly between frontal regions, but also between bilateral parietal regions (mainly SC networks). Within-hemisphere connections with strong *g* associations were observed in subcortical, parietal, temporal and frontal regions across all three networks weightings. Strong associations were found for subcortico-cortical connectivity, particularly for the thalamus, caudate, putamen and hippocampus, which mainly involved ipsilateral connectivity for SC networks, but some contralateral cortical connectivity for FA and MD networks. The widespread distribution of significant network edges across cerebral lobes, demonstrates a complex network that relies on both intra- and inter-hemispheric connectivity^19,38,40^. These findings resonate with previous studies indicating that long-range cortico-cortical connections, the genu of the corpus callosum, and subcortico-cortical pathways play a significant role in cognitive performance^19,28,38,69^.

The findings from our study broadly align with the Parieto-Frontal Integration Theory (P-FIT), which attributes intelligence to predominately frontal, parietal and temporal brain regions^20^. Previous research has provided supporting evidence for specific patterns of functional connectivity related to g, particularly in frontal and parietal regions^37–39^. Nevertheless, our study, conducted on a larger scale, demonstrates an evolving understanding by revealing additional brain areas including the insula, cingulate, subcortical and some occipital regions as potential contributors to g, some of which align with prior reports^19,23,38^. Subcortical regions, including the thalamus, hippocampus, caudate and putamen, have traditionally been associated with sensory processing, memory, and motor control^70^. However, emerging research suggests that these regions and their connections play a more prominent role in shaping g than previously thought^19,38,71^. These results reinforce the view that g is a multifaceted trait supported by the coordinated function of multiple brain regions^7,72,73^.

Our research contributes to a broader understanding of the neural bases of g by comparing the findings of our white matter network analysis with a previous study of cortical grey matter^24^. We observed strong spatial correlations (|*r*| range: 0.12 to 0.65) between node-g and cortical morphometry-g associations. Our findings provide quantitative evidence that the brain regions most strongly associated with cognitive performance differences are those that possess both larger volumes and greater local network efficiency, with both factors contributing to a distributed g-related network. The strongest correlations were observed for cortical volume and streamline count networks, both of which are volume-driven, suggesting that overall brain size influences both cortical structure and efficient white matter organisation to facilitate cognitive functioning^7^.

Age moderation effects were subtle and only significant in the large UKB sample, especially in MD networks where 78% of all nodes showed a significant age moderation. These effects appeared to follow an anterior-posterior gradient, with stronger age-related alterations observed in frontal regions, consistent with the established pattern of brain ageing where age-related changes tend to begin earlier in the late-maturing anterior areas^74,75^. These effects indicate that white matter microstructural differences become increasingly pertinent to cognitive performance with age, perhaps because the diffusion signal indexes a greater proportion of ageing-related pathological processes in older age, compared to a greater proportion of developmental variation among younger adults^69,76^.

A key strength of our study is the large sample, with over 20,000 participants included in the meta-analysis and an additional 18,000 for replication, far exceeding previous white-matter meta-analyses (*n* < 2,000)^25^, and enabling robust estimation of network-g associations across cohorts and age ranges. We also conducted network-*g* mapping at the level of individual network connections (edges), which have not been reported previously. Our network approach highlights the benefit of mapping short-range connections (e.g., U-fibres), which appear to support g but would likely be overlooked in tract-based studies focused on long-range pathways. We used standardised connectome processing, as well as a cross-cohort thresholding step to ensure consistent network density and therefore comparable network measures across cohorts.

The use of large and relatively different cohorts allowed us to assess cross-cohort consistency. While network-g associations showed considerable overlap across cohorts, we also identified notable differences. Relatively weak correlations, especially for FA and MD networks at both the node and edge levels, demonstrate the challenges of replicating connectomic findings^35,77^. Key sources of variability likely include differences in cohort composition, such as the focus on depression-related traits in STRADL, as well as differences in MRI field strength, and acquisition protocols^62,78^. In particular, differences between 1.5T and 3T scanners are known to affect dMRI measures and network density, potentially impacting the reproducibility of our findings^62^. However, in spite of these differences, cross-cohort agreement was often good and further cohort discrepancies may have been partly mitigated by use of standardised connectome processing.

Our network analysis also showed that associations between g and uncorrected SC networks were inflated by the strong correlation between SC and total brain volume; more inter-regional streamlines are identified in larger brains^44,50^. These associations were attenuated after correcting for regional node volumes. Nonetheless, the strong SC-*g* associations raise the possibility that total brain volume is partly shaped by the number of macroscopic fibre bundles, implying that individuals with more extensive brain wiring may also have larger brains, which could contribute to higher cognitive performance^73^.

Although we used *g* models known to be comparable across cohorts^5,7^, our analyses were restricted by the brief, unsupervised touchscreen assessments in UK Biobank^64^, which provided limited coverage of cognitive domains and only moderate test-retest reliability for several measures. These limitations likely attenuated the observed network-*g* associations. Nevertheless, the shared pattern of association across cohorts and the fact that network-*g* associations were well replicated in a further hold-out sample of UKB data, demonstrate the robustness of these regional relationships with *g*.

We summarise our findings as follows. First, *g* is supported by a highly distributed white matter network, indicating that cognitive performance reflects the integrity of a large-scale communication backbone that includes both volumetrically-dependent wiring capacity and size-independent microstructural organisation. These extend more widely than previous white matter tract studies suggest, involving substantial cortico-subcortical connectivity, including the thalamus, hippocampus, caudate, and putamen. Second, edge-level composite scores appear to outperform global network metrics in out-of-sample prediction, suggesting that behaviourally-relevant information is embedded in fine-grained patterns of connectivity. Third, age reshapes which aspects of neurobiology matter for cognitive functioning: the moderation patterns observed for FA and MD networks indicate that the biological features of white matter most relevant to cognitive differences change across adulthood, consistent with shifting constraints on information processing in the ageing brain.

## Materials and methods

### Participants

We mapped the associations between structural brain networks and general cognitive function (g) using data from three cohorts. The largest was the UK Biobank (UKB, n = 37,284; 19,857 female, 17,427 male, 45–83 years)^51^, randomly split into a main sample (n = 18,642, 9,911 female, 8,731 male, 45–82 years), and a hold-out replication sample (n = 18,642, 9,946 female, 8,696 male, 45–83 years). The other cohorts were Generation Scotland: Stratifying Resilience and Depression Longitudinally (STRADL, n = 937, 553 female, 384 male, 26–84 years)^52^, and the Lothian Birth Cohort 1936 (LBC1936, n = 603, 282 female, 321 male, ~73 years)^53,54^, resulting in a total meta-analytic sample of 20,182 participants. All participants provided informed consent.

These cohorts are outlined below with further details in the Supplementary Information. The UK Biobank is a large-scale epidemiology study which recruited ~500,000 adults aged 40–69 years across Great Britain (2006–2010), collecting demographic, medical, and cognitive data^51^. A subset (~40,000) underwent brain MRI 9.0 ± 1.8 years after baseline assessment^79^. Participants were generally healthy^51^. Ethical approval was granted by the North West Multi-centre Research Ethics Committee (11/NW/0382), and this study was conducted under UK Biobank application 10279. STRADL is a population-based sub-cohort (n=1188) of Generation Scotland: Scottish Family Health Study^80^, designed to study major depressive disorder, although recruitment was not restricted to individuals with mood disorders^52^. Data included brain MRI, cognitive assessment, and health measures. Participants were generally healthy, although ~30% had a lifetime history of a mood disorder. Ethical approval was obtained from the NHS Tayside Research Ethics Committee (14/SS/0039). The LBC1936 is a longitudinal study of community-dwelling older adults, most of whom took part in the Scottish Mental Survey of 1947 at ~11 years old, and who volunteered to participate in this cohort study at ~70 years old^53,54,65^. Data were drawn from the first brain imaging wave (~73 years) and included cognitive testing and medical data. Participants were generally healthy, with no self-reported symptoms of dementia, although 76 (12.6%) met criteria for neuroradiologically identified stroke. Ethical approval was given by the Multi-Centre Research Ethics Committee for Scotland, (MREC/01/0/56), the Lothian Research Ethics Committee (LREC/2003/2/29), and the Scotland A Research Ethics Committee (07/MRE00/58).

Our primary analyses included all participants. A supplementary sensitivity analysis excluded participants with neurological conditions (see Supplementary Information), but the full sample was otherwise retained because exclusion criteria were unavailable for STRADL and differed across cohorts.

### MRI acquisition and image processing

Full details of MRI acquisitions are provided in the Supplementary Information. Briefly, UKB imaging was conducted with Siemens Skyra 3T scanners at four UK sites, acquiring 3D T1-weighted (1 mm isotropic voxels) and dMRI volumes (100 directions; 2 mm isotropic voxels)^79,81^. STRADL imaging was conducted at Aberdeen (Philips Achieva 3T) and Dundee (Siemens Prisma-FIT 3T) with closely matched protocols, acquiring T1-weighted (1 mm isotropic) and dMRI (64 directions; 2.3 mm isotropic voxels)^52^. LBC1936 imaging was conducted with a GE Signa Horizon 1.5T scanner, acquiring T1-weighted (1 × 1 × 1.3 mm resolution) and dMRI (64 directions; 2 mm isotropic voxels)^82^.

All T1-weighted volumes were segmented into 85 neuroanatomical regions using FreeSurfer, incorporating the Desikan-Killiany atlas (68 cortical regions)^55,56^, and 17 subcortical structures^57,83^. Grey and white matter masks were also extracted. Processing was performed with FreeSurfer v5.1.0 for LBC1936, v5.3.0 for STRADL, and v6.0 for UKB. All dMRI volumes were processed using the FSL toolkit, with cohort-specific pipelines reflecting differences in acquisition. LBC1936 (FSL v4.1.9) and STRADL (FSL v5.0.9) data were processed locally, applying brain extraction^84^, and eddy current correction using affine registration^85^. UKB data were processed by the UKB imaging team (FSL v6.0), using a multi-shell pipeline with reverse phase encoding and distortion correction^81^. Diffusion tensors were fitted to derive mean diffusivity (MD) and fractional anisotropy (FA)^86^. Whole-brain probabilistic tractography was performed using BEDPOSTX/ProbtrackX^58–60^, with tracking initiated from white matter voxels^50^. Full processing details and quality control procedures are provided in the Supplementary Information.

### Network construction

Whole-brain structural networks were constructed using an automated pipeline harmonised across cohorts. Using T1-weighted and dMRI data, 85×85 connectivity matrices were constructed based on the 85 neuroanatomical regions described above and streamline connections obtained from probabilistic tractography^50^. Quality control and processing details are available in the Supplementary Information. After exclusions, the STRADL dataset included 937 participants (age range: 26–84 years), LBC1936 included 603 (71–74 years), and UKB included 18,642 (44.6–81.7 years). An additional 18,642 UKB participants formed the hold-out replication sample.

For each participant, three network types were computed to capture different aspects of connectivity. Streamline count (SC) represented the total uncorrected number of streamlines connecting each pair of grey matter nodes. FA and MD networks were computed as the mean values of these water diffusion measures across all voxels along the streamlines connecting each node pair. Additionally, streamline density (SD), a variant of SC, was computed by normalising streamline counts by the volumes of the corresponding grey matter regions^87^. To address variability in network density across cohorts – resulting from differences in MRI field strength, acquisition parameters, and dMRI processing – a cross-cohort thresholding procedure was applied. First, consistency-thresholding^63^ was independently performed for each cohort, retaining the top 30% most consistent connections based on findings from a large single-scanner study^50^. Second, the three cohort-specific masks were intersected to identify edges consistently present across all cohorts. This unified cross-cohort mask was then applied to all networks, ensuring comparability across cohorts.

For each network, three global graph-theoretic metrics were calculated using weighted metrics^66^: mean edge weight (mean of all edge weights per participant), global network efficiency (a measure of integration) and mean clustering coefficient (a measure of segregation). Although these metrics are highly correlated (Supplementary Fig. 7)^88^, we selected them for their widespread use in the literature. Additionally, at the node level, weighted local efficiency^66^ was computed to assess the efficiency of information exchange within the immediate neighbourhood of each node. Anatomical network plots were generated with BrainNet Viewer^89^, and circular chord diagrams using the Circos visualisation tool^90^.

### Cognitive testing

Cognitive test data were collected from cohort-specific test batteries, which covered both crystallised and fluid abilities. UKB included ten tests^51,64^, STRADL five^52^, and LBC1936 thirteen tests^65^ (details in Supplementary Information). This approach allowed us to estimate a latent factor of *g* that is known to be highly consistent across samples despite having been extracted from different tests^8,9^. For each cohort, a latent factor of *g* was estimated from these multi-domain cognitive tests using confirmatory factor analysis in a structural equation modelling (SEM) framework. For STRADL, a residual covariance was set between Mill Hill vocabulary and digit symbol substitution tests. For LBC1936, within-domain residual covariances were added for four cognitive domains (visuospatial, crystallised ability, verbal memory, and processing speed) in keeping with previous models^24^. We used a full information maximum likelihood (FIML) estimator to handle participants with missing test data. Model fits were assessed using the following fit indices: Comparative Fit Index (CFI), Tucker Lewis Index (TLI), Root Mean Square Error of Approximation (RMSEA), and the Root Mean Square Residual (SRMR).

### Statistical analysis

To ensure *g* estimates were comparable across cohorts, correlations were calculated between *g* scores and *g*-related variables, e.g., age, years of education, and highest qualification. SEM was used to estimate *g* and network associations with *g* via structural regression, adjusting for age, sex, and site as required. Analyses were conducted across three network weightings (SC, FA, MD) and for three scales: global network metrics, nodal metrics (85 regions), and edge weights (818 connections). SEM provided standardised βs, standard errors, confidence intervals, and *p*-values.

The separate network-g associations from each of the three cohorts were then used to compute a three-way meta-analysis using the ‘metafor’ package in R^91^. The meta-analysis was calculated using the βs and SEs obtained from structural regression using a random effects model with a restricted maximum likelihood estimator. This provided a meta-analytic coefficient (i.e., average β), confidence intervals and *p*-value. To evaluate cross-cohort agreement at the nodal and edge level, we computed correlations (Pearson’s r) between each set of βs for each of the three possible cohort pairs. Node plots were ordered by the β value averaged across cerebral hemispheres for each of the 42 bilateral grey matter regions; the brainstem, as the only unpaired midline node, was included in all analyses but was not shown in node plots. Lateralisation of network-g associations was evaluated by correlating meta-analytic βs across hemispheres at the node level. For all significant meta-analysed edge-g associations, we quantified intra- and inter-hemispheric and intra- and inter-lobar edges to summarise the distribution of *g*-related connectivity.

Additionally, we assessed the relationship between the meta-analytic node-g associations and previously reported *g* associations of cortical surface measures in the same cohorts^24^. Using 68 Desikan-Killiany regions^55^, correlations were computed between βs from node-g associations and those from cortical surface measures (volume, area, thickness). Given LBC1936’s narrow age range (71–74 years), age moderation analyses were conducted only in UKB (45–81 years) and STRADL (26– 84 years) for global and nodal network properties, with significant interactions interpreted using conditional standardised slopes estimated at −1 SD, the mean, and +1 SD of age.

To assess the capacity to predict *g* from network edges, we calculated a composite score using bivariate prediction. For all samples including the UKB hold-out sample, variables were standardised, and a weighted sum was computed (∑ β_k_ x edge_k_, where k=818) using the β coefficients from the meta-analysis.

Statistical analyses were performed in R v4.2.2 using lavaan v0.6-12 and metafor v3.8-1. Significance was determined at p < 0.05 and all p-values were corrected for multiple comparisons using false discovery rate (FDR)^92^.

## Acknowledgements

We thank all study participants, the respective study imaging teams, the local research teams and radiographers who contributed to data collection. UK Biobank research was conducted using the UK Biobank Resource under approved project 10279 at The University of Edinburgh. Generation Scotland/STRADL is supported by the Wellcome Trust [216767/Z/19/Z, 104036/Z/14/Z], CSO of the Scottish Government Health Directorates [CZD/16/6], and the Scottish Funding Council [HR03006]. The LBC1936 is supported by the BBSRC and the ESRC [BB/W008793/1], Age UK [Disconnected Mind project], the MRC [G0701120, G1001245, MR/M013111/1, MR/R024065/1], and the University of Edinburgh. CRB, MEB, IJD, EMT-D and SRC were supported by a NIH research grant [R01AG054628], and CRB also by a Biotechnology and Biological Sciences Research Council project grant (BB/T000813/1). ELSC is Junior Research Fellow in Applied AI supported by Lady Margaret Hall, University of Oxford [EPT-AI]. EMT-D is a member of the Population Research Center at the University of Texas at Austin, which is supported by NIH Center Grant [P2CHD042849]. SRC and JEM are supported by SRC’s Sir Henry Dale Fellowship, jointly funded by the Wellcome Trust and the Royal Society [221890/Z/20/Z]. JMW is part funded by the UK Dementia Research Institute (UKDRI – 4002; UKDRI – 4205) which receives its funding from the UK Medical Research Council, Alzheimer’s Society and Alzheimer’s Research UK, and by the UK National Institutes of Health and Care Research Health Technology Assessment Panel (NIHR159009).

