## Supplementary Information for "General cognitive function and the brain’s structural connectome"

­­­­

­­­­Colin R. Buchanan^1,2,3^ *, Joanna E. Moodie^1,2^, Venia Batziou^4^, Eleanor L. S. Conole^1,2,5^, Susana Muñoz Maniega^1,3,6^, Mathew A. Harris^1,2^, Hon Wah Yeung^1,2^, Janie Corley^1,2^, David C. Liewald^1,2^, Paul Redmond^1,2^, J. Douglas Steele^7^, Gordon D. Waiter^3,8^, Heather C. Whalley^9,10^, Andrew M. McIntosh^9,10^, Joanna M. Wardlaw^1,3,6,11^, Michelle Luciano^1,2^, Mark E. Bastin^1,3,6^, Ian J. Deary^1,2^, Elliot M. Tucker-Drob^12^, Simon R. Cox^1,2,3^

1. Department of Psychology, The University of Edinburgh, Edinburgh, UK.

2. Lothian Birth Cohorts, Edinburgh Futures Institute, The University of Edinburgh, Edinburgh, UK.

3. Scottish Imaging Network, A Platform for Scientific Excellence (SINAPSE) Collaboration, UK.

4. College of Medicine & Veterinary Medicine, The University of Edinburgh, Edinburgh, UK.

5. Department of Biochemistry, University of Oxford, Oxford, UK.

6. Institute for Neuroscience and Cardiovascular Research, The University of Edinburgh, Edinburgh, UK.

7. Division of Neuroscience, University of Dundee, Dundee, UK

8. Aberdeen Biomedical Imaging Centre, University of Aberdeen, Aberdeen UK

9. Department of Psychiatry, The University of Edinburgh, Edinburgh, UK.

10. Generation Scotland, Centre for Medical Informatics, Usher Institute, The University of Edinburgh, Edinburgh, UK

11. UK Dementia Research Institute Centre, The University of Edinburgh, Edinburgh, UK

12. Department of Psychology, The University of Texas at Austin, Austin, USA

### Detailed materials and methods

#### Participants

We evaluated the associations between white matter networks and general cognitive function (*g*) using data from three large and well-characterised cohorts. The largest cohort was UK Biobank (UKB; *n* = 37,284; aged 45–82 years)^1^, which was randomly split into two halves, comprising a main sample (*n* = 18,642), and a held-out sample for replication purposes. The other cohorts were Generation Scotland: Stratifying Resilience and Depression Longitudinally (STRADL; *n* = 937; 26–84 years)^2^, and the Lothian Birth Cohort 1936 (LBC1936; *n* = 603; ~73 years)^3,4^. This resulted in a total sample size of 20,182 participants for meta-analysis. All participants provided informed consent to participate as stated below.

The UK Biobank is a large-scale epidemiology study which recruited approximately 500,000 community-dwelling adults aged 40–69 years from across Great Britain between 2006 and 2010^1^. Participants provided comprehensive demographic, psychosocial, medical information, blood samples for DNA extraction, physical fitness testing, and cognitive assessment during an initial visit to a UK Biobank assessment centre. A subset (~40,000) underwent brain MRI 9.0 ± 1.8 years after baseline assessment^5^. Participants were generally healthy, and the full connectome sample included 37,284 participants. A subset of 2310 participants (6.2%) had at least one of 26 self-reported neurological conditions (e.g., stroke, epilepsy, multiple sclerosis, meningitis, Parkinson’s; see full details below), and these participants were excluded in a supplementary analysis (below). UK Biobank received ethical approval from the North West Multi-centre Research Ethics Committee (REC reference 11/NW/0382). The current study was conducted under approved UK Biobank application number 10279.

STRADL is a population-based study, developed from the Generation Scotland: Scottish Family Health Study (GS:SFHS)^6^. A subset of these participants (*n* = 1188) were invited to take part in an additional study (STRADL), which was primarily designed to study major depressive disorder, although participants were not selected based on depression^2^. STRADL participants underwent brain MRI and data collection included socio-economic and lifestyle variables, physical measures, mental health questionnaires, laboratory samples, and cognitive assessment. Participants were generally healthy, although approximately 30% had a lifetime history of a mood disorder. NHS records were previously obtained as part of the GS:SFHS (05/S1401/89) and subsequent procedures were conducted following an independent, but linked, NHS Tayside Committee on Research Ethics application (14/SS/0039).

The LBC1936 is a longitudinal study of community-dwelling older adults, most of whom took part in the Scottish Mental Survey of 1947 at ~11 years old, and who volunteered to participate in this cohort study at ~70 years old^3,4,7^. Data have been gathered on the influences on cognitive ageing from age 11 into the 8th and 9th decades of life. Participants underwent a thorough assessment including medical interviews, physical fitness tests, extensive cognitive assessments, completion of psycho-social questionnaires, and blood samples for DNA extraction and biomarker analyses. LBC1936 data were drawn from all available participants from the first brain imaging wave (~73 years of age). Participants were generally healthy, with no self-reported symptoms of dementia, while 76 (12.6%) met criteria for neuroradiologically identified stroke. The LBC1936 study was given ethical approval by the Multi-Centre Research Ethics Committee for Scotland, (MREC/01/0/56), the Lothian Research Ethics Committee (LREC/2003/2/29) and the Scotland A Research Ethics Committee (07/MRE00/58).

#### MRI acquisition

Details of the UKB MRI protocol and processing are freely available^5,8^. All imaging data used in this study were acquired using one of four Siemens Skyra 3T scanners across four UK imaging centres (Cheadle, Newcastle upon Tyne, Reading, and Bristol). 3D T1-weighted volumes were acquired using a magnetization-prepared rapid gradient-echo sequence at 1 × 1 × 1 mm resolution with 208 × 256 × 256 field of view. The dMRI data were acquired using a spin-echo echo-planar imaging sequence (50 *b* = 1000 s/mm2, 50 *b* = 2000 s/mm2 and 5 *b* ≈ 0 s/mm2) resulting in 100 distinct diffusion-encoding directions. The field of view was 104 × 104 mm with imaging matrix 52 × 52 and 72 slices with slice thickness of 2 mm resulting in 2 × 2 × 2 mm voxels.

Details of the STRADL protocol have been described previously^2^. Acquisition took place at two sites with the MRI acquisition parameters matched as closely as practical: Aberdeen (*n* = 582, 49%) and Dundee (*n* = 606, 51%). 3D T1-weighted fast gradient echo volumes with magnetisation preparation were acquired. The scanner used in Aberdeen was a 3T Philips Achieva TX-series MRI system (Philips Healthcare, Best, Netherlands) with a 32-channel phased-array head coil and a back facing mirror (gradients with maximum amplitude 80 mT/m and maximum slew rate 100 T/m/s). For T1-weighted images, 160 sagittal slices were acquired with a field of view of 240 mm and a matrix size of 240 x 240 pixels, giving a resolution of 1 × 1 × 1 mm. For the dMRI protocol, echo-planar (EP) diffusion-weighted whole-brain volumes (*b* = 1200 s mm^− 2^) were acquired in 64 non-collinear directions, along with eight T2-weighted volumes (*b* = 0 s mm^− 2^). Sixty contiguous 2.3 mm thick axial slices were acquired with a matrix size of 96 × 94 resulting in 2.3 mm^3^ isotropic voxels. The scanner used in Dundee was a Siemens 3T Prisma-FIT (Siemens, Erlangen, Germany) with 20 channel head and neck phased array coil and a back facing mirror (Syngo E11, gradient with max amplitude 80 mT/m and maximum slew rate 200 T/m/s). For T1-weighted images 208 sagittal slices were acquired with a field of view of 256 mm and matrix size 256 × 256 pixels giving a resolution of 1 × 1 × 1 mm. For the dMRI protocol, echo-planar (EP) diffusion-weighted whole-brain volumes (*b* = 1200 s mm^− 2^) were acquired in 64 non-collinear directions, along with eight T2-weighted volumes (*b* = 0 s mm^− 2^). Sixty contiguous 2.3 mm thick axial slices were acquired with a matrix size of 96 × 94 resulting in 2.3 mm^3^ isotropic voxels.

Details of the LBC1936 MRI protocol have been described previously^9^. All participants underwent brain MRI on the same 1.5T GE Signa Horizon HDx clinical scanner (General Electric, Milwaukee, WI, USA) with a manufacturer supplied 8-channel phased-array head coil. High resolution 3D T1-weighted inversion-recovery prepared, fast spoiled gradient-echo volumes were acquired in the coronal plane with 180 contiguous 1.3 mm thick slices resulting in voxel dimensions of 1 × 1 × 1.3 mm. For the dMRI protocol, single-shot spin-echo echo-planar (EP) diffusion-weighted whole-brain volumes (*b* = 1000 s mm^− 2^) were acquired in 64 non-collinear directions, along with seven T2-weighted volumes (*b* = 0 s mm^− 2^). Seventy-two contiguous axial 2 mm thick slices were acquired resulting in 2 mm^3^ isotropic voxels.

#### T1-weighted processing

Each T1-weighted image was segmented into 85 distinct neuroanatomical regions-of-interest (ROIs) using the volumetric segmentation and cortical reconstruction of the FreeSurfer image analysis suite (<http://surfer.nmr.mgh.harvard.edu>). The Desikan-Killiany atlas was used to delineate 34 cortical structures per hemisphere^10,11^. Subcortical segmentation was applied to obtain the brain stem and eight grey matter structures per hemisphere: accumbens area, amygdala, caudate, hippocampus, pallidum, putamen, thalamus and ventral diencapahlon^12,13^. Grey and white tissue matter masks were also obtained. Cortical processing was conducted locally using FreeSurfer v5.1.0 for LBC1936, and v5.3.0 for STRADL. UKB processing was conducted by the UKB imaging team using FreeSurfer v6.0.

#### FreeSurfer quality control procedure

The quality control (QC) approach differed by cohort. The LBC1936 Wave 2 data were collected between 2007-2010 and were processed, edited and underwent QC using FreeSurfer v5.1.0. STRADL data was processed using FreeSurfer v5.3.0. Manual QC of FreeSurfer data for STRADL and LBC1936 involved visual inspection of skull stripping, tissue segmentation and regional boundaries followed by manual editing to rectify segmentation and parcellation errors; participants with uncorrectable errors were excluded. UKB segmentations underwent automated quality checking and exclusion, which was conducted by the UKB imaging team^14^.

#### Diffusion MRI processing and tractography

All dMRI data were processed with the FSL toolkit (FMRIB, Oxford University: <http://www.fmrib.ox.ac.uk>)^15^. However, processing differed between cohorts due to limitations of the acquisition (e.g., absence of reverse phase encodings) and the FSL version available at the time of processing. All LBC1936 (FSL v4.1.9) and STRADL (FSL v5.0.9) data were processed locally as follows. Data underwent brain extraction^16^ performed on the T2-weighted EP volumes acquired along with the dMRI data. The brain mask was applied to all volumes after correcting for systematic eddy-current induced imaging distortions and head motion using affine registration to the first T2-weighted EP volume of each participant with ‘eddy_correct’^17^. UKB processing was performed by the UKB imaging team (FSL v6.0) using an UKB-specific pipeline that exploits the reverse phase encoding and multi-shell acquisition of the UKB dMRI data^8^. Briefly, UKB processing involved field map generation and subsequent correction for eddy currents, systematic distortions and head motion. For all dMRI volumes, diffusion tensors were fitted at each voxel (FSL dtifit). Water diffusion measures were estimated for: mean diffusivity (MD), which measure magnitudes of molecular water diffusion; and fractional anisotropy (FA), which measures the degree of anisotropic diffusion per voxel^18^. All diffusion measures and tractography were computed in diffusion space.

Whole-brain tractography was performed using an established probabilistic algorithm (BEDPOSTX/ProbtrackX)^19,20^. Probability density functions, which describe the uncertainty in the principal directions of diffusion, were computed with a multi-fibre model (two-fibres for LBC1936 and STRADL; three-fibres for UKB) per voxel^20,21^. Streamlines were then constructed by sampling from these distributions during tracking using 100 Markov Chain Monte Carlo iterations with a fixed step size of 0.5 mm between successive points. Tracking was initiated from all white matter voxels using stopping criteria as described previously^22^.

#### Network construction

Construction of whole-brain networks was harmonised as much as practical across cohorts using an automated connectivity mapping pipeline. For all participants with compatible T1 and dMRI data, white matter structural networks were constructed using the 85 neuroanatomical regions described above and the set of streamlines generated by tractography resulting in 85 × 85 connectivity matrices^22^. Networks were constructed by identifying connections between all ROI pairs. The endpoint of a streamline was recorded as the first ROI encountered (if any) when tracking from the seed location.

For each participant, three network types were computed reflecting different aspects of connectivity. Streamline count (SC) was computed by recording the total streamline count (uncorrected) between each pair of ROIs. FA and MD networks were computed by computing the mean value of each measure in all voxels along the interconnecting streamlines between each pair of ROIs. In addition, a variant of streamline count termed streamline density (SD) was computed by dividing counts by the volumes of the grey matter ROIs involved^23^.

As network density, i.e., the number of network connections identified, is known to depend on MRI field strength, acquisition parameters and dMRI processing, we chose to apply a cross-cohort network thresholding procedure to match network density across cohorts. First, consistency-thresholding^24^ was independently applied to each cohort to remove putatively spurious connections at a threshold level retaining the top 30% most consistent connections – this density was previously determined from a large single-scanner study in a portion of the UKB data^22^. Second, the three cohort masks (containing non-identical sets of connections) were combined and only connections (edges) found to be present in all three cohorts were retained. The resulting cross-cohort mask was then applied to each network across all participants.

For each network, three global graph-theoretic metrics were computed using weighted metrics from the Brain Connectivity Toolbox^25^: mean edge weight (mean of all edge weights per participant), global network efficiency (a measure of integration) and network clustering coefficient (a measure of segregation). Global efficiency measures how easily information can flow across an entire network, reflecting the network's overall ability to connect different nodes efficiently. Greater global efficiency signifies more effective ‘wiring’ across the entire brain. The mean clustering coefficient measures on average how many local clusters exist around a node in terms of its direct neighbours, indicating how tightly interconnected neighbouring nodes are within a network. A higher nodal coefficient indicates that nearby network nodes are densely interconnected, potentially reflecting specialised functions within those clusters. Although global graph-theory metrics are known to be highly correlated^26^, we chose to assess these three commonly used metrics. At the node level, only weighted local efficiency was assessed, quantifying how efficiently information is exchanged within the immediate neighbourhood of each node based on the strength of connections between neighbouring nodes^25^.

#### Connectome quality control procedure

An initial QC of whole-brain tractography was conducted on a randomly selected subset of STRADL participants (*n* = 50). This involved visual inspection of successful interregional streamlines alongside slices from the FA volume using TrackVis visualisation software (<https://trackvis.org>). This pilot QC indicated that low streamline counts could serve as a heuristic for identifying major tractography errors or other systematic errors during processing. Based on these findings, QC was performed on a subset of 80 participants, leading to the exclusion of 19 due to aberrant streamlines and sparse cortical connectivity. The final STRADL dataset included 938 connectome participants.

For UKB, we performed a similar manual QC of connectome outputs in a subset of 388 out of 37,436 participants (available imaging sample at time of processing) that were identified as outliers (± 3 SDs) based on the mean edge weights for SC, FA or MD using unthresholded networks. The QC for these outliers involved visual inspection of the successful interregional streamlines alongside slices from the FA volume. Exclusions were made for major tractography failures or problems in the FA volume for which 152 out of 388 (39.2%) outliers were excluded. 50 of these exclusions were due to tracking failures caused by incomplete FreeSurfer segmentations. Other reasons for failure were: signal dropout, warping or distortions in the dMRI (FA) volume; sparse or aberrant connectivity to cortical areas; and tracking failure across the corpus callosum. We observed that a low total number of interregional streamlines (i.e., 3 SDs below the mean SC) was often indicative of tractography error or other systematics errors during processing.

For LBC1936, we performed a manual QC of connectome outputs in a subset of 64 out of 615 participants (wave 2) that were identified as outliers (± 2 SDs) based on the mean edge weights for SC, FA or MD using unthresholded networks. We selected a more stringent outlier criteria than for UKB due to the more elderly sample. As above, the QC for these outliers involved visual inspection of the successful interregional streamlines alongside slices from the FA volume. Exclusions were made for major tractography failures or problems in the FA volume for which 12 out of 614 (18.8%) outliers were excluded. Four fails were due to problems in dMRI volumes (motion artefacts or poor eddy current correction), while eight fails had aberrant or sparse tractography in cortical areas.

After removing processing failures and quality checking, the STRADL dataset comprised 937 connectome participants (553 female, 384 male, mean age 59.5 years, 26.0–84.0 years), the UKB main sample had 18,642 participants (9,911 female, 8,731 male, mean age 63.7 years, 44.6–81.7 years), and LBC1936 had 603 participants (282 female, 321 male, mean age 72.7 years, 71.0–74.2 years). In addition, a further 18,642 UKB participants (9,946 female, 8,696 male, mean age 64.0, 44.8-82.7 years) were processed as a held-out replication sample.

#### Cognitive tests

Cognitive test data were collected as described in detail elsewhere: UKB^27^, STRADL^2^, LBC1936^28^. Although the test batteries varied across cohorts, tests were selected to cover multiple cognitive domains in each, including crystallised and fluid cognitive ability domains. This approach allowed us to estimate a latent factor of *g* that is known to be highly consistent across samples despite having been extracted from different tests^29,30^. The tests used in the present study are detailed below and summarised in Supplementary Tables 2-4.

The ten cognitive tests used for UKB^1,27^ were: reaction time (time taken to respond in snap-type computer game; UKB data field 20023), number span (maximum length of number string recalled; field 4282), verbal-numerical reasoning (number of 13 verbal and numerical logic questions correct; field 20016), trail making B (time taken to complete an alphanumeric path-making test; field 6350), matrix pattern (number of matrix pattern puzzles solved; field 6373)^31^, tower task (number of tower task puzzles solved; field 21003), digit-symbol substitution (number of digit-symbol pairs matched; field 23324)^32^, pairs matching (number of incorrect matches in a 6-pair matching game where zero indicates the participant made no mistakes; field 399), prospective memory (test of instruction recall after a delay; field 20018), and paired associates (number of novel word pairs correctly recalled after a delay; field 20197). We coded prospective memory as a binary variable with one indicating success at the first recall attempt. We removed trail making B scores for those who did not complete the test. Reaction time, trail making B, and pairs matching variables were log transformed and then multiplied by -1 so that a higher score indicated better performance.

The five tests used for STRADL^2^ were: matrix reasoning (number of puzzles correct;^31^, verbal fluency (number of words recalled beginning with C, F and L in 3x1 minute)^33^, Mill Hill vocabulary (number of word meanings explained correctly)^34^, digit symbol substitution (number of digit-symbol pairs matched)^35^, and logical memory (story recall score - total score from immediate and delayed tests)^35^.

The thirteen tests used for LBC1936^7^ were: matrix reasoning (number of puzzles correct)^35^, block design (number of puzzles correct in two minutes)^35^, spatial span (number of block sequences correct)^36^, National Adult Reading Test (number of words pronounced correctly)^37^, Wechsler test of adult reading (number of words pronounced correctly)^38^, verbal fluency (number of words recalled beginning with C, F and L in 3x1 minute)^39^, verbal paired associates (number of novel word pairs matched in recall - total from immediate and delayed tests)^36^, logical memory (number of story details recalled out of a total possible of 25 from immediate and delayed tests)^36^, digit span backwards (maximum number of a string of numbers recalled in reverse)^35^, symbol search (number of symbols correctly detected), digit-symbol substitution (number of digit-symbol pairs matched)^32^, inspection time (number of correct responses)^40^, and four-choice reaction time(s) (time taken to press the indicated button out of 4 buttons)^41^. Reaction times were multiplied by -1 so that a higher score indicated better performance.

For each cohort, a latent factor of *g* was estimated from these multi-domain cognitive tests using confirmatory factor analysis (CFA) in a structural equation modelling (SEM) framework. For STRADL, a residual covariance was set between Mill Hill vocabulary and digit symbol substitution tests. For LBC, within-domain residual covariances were added for four cognitive domains (visuospatial, crystallised ability, verbal memory, and processing speed) in keeping with previous models^42^. We used a full information maximum likelihood estimator to handle participants with missing test data. Model fits were assessed using the following fit indices: Comparative Fit Index (CFI), Tucker Lewis Index (TLI), Root Mean Square Error of Approximation (RMSEA), and the Root Mean Square Residual (SRMR).

### Supplementary results

#### Estimation of *g* from cognitive tests

For each cohort, we estimated a latent factor of *g* from multi-domain cognitive tests using CFA. Participants without any cognitive test data were excluded from subsequent analysis, resulting in the exclusion of 1628 participants from UKB. To handle missing test data, we employed a full information maximum likelihood estimator. The number of remaining participants with missing data for at least one cognitive test was: 5206 (30.6%) for UKB, 26 (4.3%) for LBC1936, and 8 (0.9%) for STRADL. All three CFA models resulted in an acceptable fit (all CFI > 0.94; TLI > 0.92; RMSEA < 0.06; SRMR < 0.05; Supplementary Table 1). Consistent with the extensive literature on the positive manifold in *g*^43,44^, and previous work using the present data^42^, cognitive tests showed a relatively even pattern of loadings across indicators (Supplementary Tables 2-4). For each cohort, all estimated paths to latent *g* were significant (*p* < 0.001).

#### Associations of *g* with age and education

To evaluate whether *g* estimates showed comparable associations with relevant covariates across cohorts, we examined correlations between *g* scores and variables known to relate to cognitive ability. These variables included age, age at leaving full-time education, duration of education, and highest educational qualification (Supplementary Table 5). Not all variables were available in every cohort. In addition, highest qualification was coded differently in UKB and LBC1936, and duration of education was measured in years in LBC1936 but on an ordinal scale in STRADL (Compulsory = 1; More than compulsory = 2; Post-secondary = 3). Despite the relatively narrow age range in LBC1936 (~3 years), *g* was negatively associated with age in all cohorts (*r* range: −0.36 to −0.10). Furthermore, despite scoring differences, correlations were comparable across cohorts for duration of education (*r* = 0.39 and 0.47) and highest qualification (*r* = 0.35 and 0.55).

#### Scanner-specific differences in network properties

Overall, network organisation was comparable across cohorts, although expected scanner-related differences were observed in network density and weight distributions (Supplementary Fig. 1). The mean edge weights for LBC1936 networks (all acquired at 1.5T MRI) were found to be lower than in the other two cohorts (both 3T MRI), particularly for FA and MD networks, although these cross-cohort differences were smaller for SC networks. Higher field strength acquisitions are known to identify more white matter connections than lower field strengths and differences in acquisition and pre-processing can also lead to apparent differences in the resulting FA and MD values^45^. Additionally, STRADL data were acquired on two different scanners (Aberdeen: Philips Achieva 3T; Dundee: Siemens 3T Prisma-FIT), but these acquisitions were well-matched, and the diffusion measures were broadly in the same range (mean edge weight range for FA: 0.35–0.48 for Aberdeen, and 0.33–0.46 for Dundee).

#### Effect of neurological exclusion on network-*g* associations

Our main analyses included all participants. However, we found that applying exclusion for neurological conditions led to some attenuation in *g* associations. In UKB, participants were excluded for any of the following 26 self-reported conditions (UKB codes 20001 and 20002):

brain cancer, meningeal cancer, stroke, transient ischaemic attack, subdural haematoma, subarachnoid haemorrhage, neurological trauma, infection of nervous system, brain abscess, encephalitis, meningitis, Guillain-Barré syndrome, other chronic neurological problems, motor neurone disease, multiple sclerosis, Parkinson’s disease, epilepsy, head injury, other demyelinating diseases, brain aneurysm, cerebral palsy, brain haemorrhage, spina bifida, ischaemic stroke, meningioma (benign). After excluding 1172 participants (6.3%) with neurological conditions in UKB, global network-*g* associations were modestly attenuated (0.8–7.3%). Information on neurological conditions differed between LBC1936 and UKB. In LBC1936, excluding 76 participants (12.6%) with neuroradiologically identified stroke weakened the network-*g* associations (attenuation: 2.1–28.2%), with two exceptions: the clustering coefficient with SC increased by 10.5%, and mean edge weight with MD remained null. Across both cohorts, FA and MD measures showed greater attenuation (UKB: 4.8–7.3%; LBC1936: 12.9–28.2%) than SC (UKB: 0.8–1.1%; LBC1936: 2.1–5.5%), though attenuations were broadly similar across network metrics. As neurological exclusion criteria were unavailable for STRADL at the time of analysis and the exclusion criteria differed across the other two cohorts, we conducted all main analyses using the full samples, which also preserved statistical power. Exclusions were applied only for these global associations for comparison.

#### Influence of brain volume on streamline counts

Our main analyses were based on streamline counts that were uncorrected for grey or white matter volumes. However, there is evidence that uncorrected streamline counts are influenced by head size and more interregional streamlines are identified in large brains than in small brains, given that voxel dimensions are constant across participants. We observed that the mean edge weight for SC networks was strongly correlated with total brain volume (*r* = 0.80 in UKB), but this was not the case for the diffusion tensor-based FA or MD networks (*r* < 0.1). However, it is unclear how best to apply volume correction of streamline counts to avoid overcorrection for volume-driven effects on these streamline weightings^22,46^. Consequently, we investigated node-*g* associations for a variant of streamline count, termed, streamline density (SD), which corrects SC by dividing each edge count by the combined volume of the pair of grey matter ROIs involved. Notably, the node-*g* associations were lower for SD (β range: -0.14 to 0.19; Supplementary Table 8) than for SC (β range: -0.10 to 0.27). However, the meta-analysis of node strength for SD networks found that 28/85 (32.9%) nodes retained significant node-*g* associations compared to 72/85 (84.7%) for SC. The cross-cohort correlations based on the node βs for SD were moderate (*r* range: 0.34 to 0.48).

#### Analysis of significant edge-*g* associations

Significant intra-hemisphere connections were considerably more numerous than inter-hemisphere connections with this ratio exceeding 4:1 across all network weightings (Supplementary Table 15). However, across all lobes, the significant intra-lobe connections were fewer than the significant connections to other lobes (Supplementary Table 16). These inter-lobe connections were predominantly ipsilateral rather than contralateral. This suggests that associations of *g* at the lobular level were facilitated more by ipsilateral inter-lobe connectivity than within-lobe connectivity.

The significant network edges obtained from the meta-analyses revealed patterns of strong (|β| > 0.1) intra- and inter-hemispheric connectivity to all cerebral lobes (SC, FA and MD networks). Strong associations for inter-hemispheric connections were observed, particularly between frontal regions (SC, FA and MD), and between bilateral parietal regions (SC). The inter-hemispheric associations were more widespread in MD networks than SC or FA. For all networks, strong associations were found for the precuneus and insula, which are considered integral to network communication. Strong associations were also found for subcortical connectivity, particularly for the thalamus, caudate, putamen and hippocampus, which mainly involved ipsilateral connectivity for SC networks, but some contralateral cortical connectivity for FA and MD networks. We observed little difference between the β distributions of significant intra- and inter-hemisphere connections (Supplementary Fig. 5A). We avoided conducting statistical tests of differences in these βs, because it would be invalid to test a quantity that has already been selected based on significance. Therefore, the following are treated as descriptive. MD networks appeared to show marginally stronger associations for inter- rather than intra-hemisphere connections. For lobe connectivity, we found little difference between β distributions for significant intra- and inter-lobe edges (Supplementary Fig. 5B). However, for FA and MD networks, the connections involving frontal and parietal nodes tended to have stronger associations compared to those involving subcortical, temporal or occipital nodes (Supplementary Fig. 5B and Supplementary Table 16).

### Tables

**Supplementary Table 1**. Model fit measures for the latent g model as estimated by confirmatory factor analysis. A single latent factor of g was estimated for each cohort: UKB (n=17,014), STRADL (n=937), and LBC1936 (n=603). The measures are: χ2, degrees of freedom (df), Comparative Fit Index (CFI), Tucker Lewis Index (TLI), Root Mean Square Error of Approximation (RMSEA), and the Root Mean Square Residual (SRMR).

| **Study** | ***X*^2^** | ***df*** | **CFI** | **TLI** | **RMSEA** | **SRMR** |
| --- | --- | --- | --- | --- | --- | --- |
| UKB | 1318.668 | 35 | 0.943 | 0.926 | 0.046 | 0.031 |
| STRADL | 12.566 | 4 | 0.986 | 0.965 | 0.048 | 0.022 |
| LBC1936 | 155.516 | 50 | 0.967 | 0.949 | 0.059 | 0.042 |

**Supplementary Table 2**. UKB cognitive tests used in latent g model with factor loadings (standardised estimates) for individual tests. All estimated paths to latent g were significant (p < 0.001).

| **Test** | ***n*** | **Standardised estimate** | ***SE*** |
| --- | --- | --- | --- |
| Reaction time | 16905 | 0.352 | 0.008 |
| Number span | 12800 | 0.462 | 0.008 |
| Verbal-numerical reasoning | 16684 | 0.621 | 0.007 |
| Trail making B | 12084 | 0.733 | 0.006 |
| Matrix pattern | 12388 | 0.630 | 0.007 |
| Tower task | 12276 | 0.564 | 0.007 |
| Digit-symbol substitution | 12394 | 0.612 | 0.007 |
| Pairs matching | 17014 | 0.279 | 0.009 |
| Prospective memory | 17007 | 0.364 | 0.008 |
| Paired associates | 12530 | 0.451 | 0.008 |

**Supplementary Table 3**. STRADL cognitive tests used in latent g model with factor loadings (standardised estimates) for individual tests. All estimated paths to latent g were significant (p < 0.001).

| **Test** | ***n*** | **Standardised estimate** | ***SE*** |
| --- | --- | --- | --- |
| Matrix reasoning | 937 | 0.545 | 0.031 |
| Verbal fluency | 937 | 0.483 | 0.031 |
| Mill Hill vocabulary | 937 | 0.721 | 0.035 |
| Digit symbol substitution | 929 | 0.573 | 0.039 |
| Logical memory | 937 | 0.447 | 0.031 |

**Supplementary Table 4**. LBC1936 cognitive tests used in latent g model with factor loadings (standardised estimates) for individual tests. All estimated paths to latent g were significant (p < 0.001).

| **Test** | ***n*** | **Standardised estimate** | ***SE*** |
| --- | --- | --- | --- |
| Matrix reasoning | 602 | 0.572 | 0.034 |
| Block design | 601 | 0.595 | 0.034 |
| Spatial span | 602 | 0.399 | 0.042 |
| National Adult Reading Test | 602 | 0.666 | 0.031 |
| Wechsler test of adult reading | 602 | 0.681 | 0.030 |
| Phonemic verbal fluency | 602 | 0.489 | 0.038 |
| Verbal paired associates | 592 | 0.543 | 0.037 |
| Logical memory | 602 | 0.560 | 0.036 |
| Digit span backwards | 603 | 0.573 | 0.035 |
| Symbol search | 602 | 0.591 | 0.035 |
| Digit-symbol substitution | 602 | 0.644 | 0.031 |
| Inspection time | 594 | 0.378 | 0.042 |
| Four-choice reaction time (s) | 603 | 0.403 | 0.041 |

**Supplementary Table 5**. Correlations (Pearson’s r) between g scores and demographic variables in each cohort: age, age at leaving full-time education, duration of education, and highest qualification obtained. Note the narrow age range in the LBC1936 cohort (~3 years). Dash indicates that information was not available. All correlations were significant (p < 0.01, uncorrected).

| **Study** | **Age** | **Age left full-time education** | **Duration of education** | **Highest qualification** |
| --- | --- | --- | --- | --- |
| UKB | -0.365 | 0.169 | - | 0.350 |
| STRADL | -0.102 | - | 0.391 | - |
| LBC1936 | -0.154 | 0.465 | 0.465 | 0.547 |

**Supplementary Table 6**. Associations (standardised estimates) between g and global network metrics in all cohorts, including the meta-analysis, across weightings (SC, FA, and MD), and across network metrics (mean edge weight, global efficiency, mean clustering coefficient). Standard errors (SE) and p-values were estimated using Structural Equation Modelling (SEM), and q-values were adjusted by False Discovery Rate (FDR).

| **Study** | **Weighting** | **Metric** | **β** | ***SE*** | ***p*** | ***q*** |
| --- | --- | --- | --- | --- | --- | --- |
| UKB | SC | Mean edge weight | 0.159 | 0.009 | 0.000 | 0.000 |
| UKB | SC | Efficiency | 0.152 | 0.009 | 0.000 | 0.000 |
| UKB | SC | Clustering | 0.166 | 0.009 | 0.000 | 0.000 |
| UKB | FA | Mean edge weight | 0.063 | 0.008 | 0.000 | 0.000 |
| UKB | FA | Efficiency | 0.058 | 0.008 | 0.000 | 0.000 |
| UKB | FA | Clustering | 0.066 | 0.008 | 0.000 | 0.000 |
| UKB | MD | Mean edge weight | -0.061 | 0.009 | 0.000 | 0.000 |
| UKB | MD | Efficiency | -0.075 | 0.009 | 0.000 | 0.000 |
| UKB | MD | Clustering | -0.061 | 0.009 | 0.000 | 0.000 |
| STRADL | SC | Mean edge weight | 0.277 | 0.036 | 0.000 | 0.000 |
| STRADL | SC | Efficiency | 0.267 | 0.037 | 0.000 | 0.000 |
| STRADL | SC | Clustering | 0.280 | 0.035 | 0.000 | 0.000 |
| STRADL | FA | Mean edge weight | 0.293 | 0.048 | 0.000 | 0.000 |
| STRADL | FA | Efficiency | 0.300 | 0.052 | 0.000 | 0.000 |
| STRADL | FA | Clustering | 0.272 | 0.049 | 0.000 | 0.000 |
| STRADL | MD | Mean edge weight | -0.159 | 0.060 | 0.008 | 0.008 |
| STRADL | MD | Efficiency | -0.216 | 0.056 | 0.000 | 0.000 |
| STRADL | MD | Clustering | -0.214 | 0.058 | 0.000 | 0.000 |
| LBC1936 | SC | Mean edge weight | 0.179 | 0.044 | 0.000 | 0.000 |
| LBC1936 | SC | Efficiency | 0.157 | 0.044 | 0.000 | 0.002 |
| LBC1936 | SC | Clustering | 0.120 | 0.045 | 0.007 | 0.016 |
| LBC1936 | FA | Mean edge weight | 0.127 | 0.045 | 0.005 | 0.014 |
| LBC1936 | FA | Efficiency | 0.108 | 0.045 | 0.015 | 0.028 |
| LBC1936 | FA | Clustering | 0.100 | 0.045 | 0.026 | 0.039 |
| LBC1936 | MD | Mean edge weight | -0.006 | 0.046 | 0.902 | 0.902 |
| LBC1936 | MD | Efficiency | -0.084 | 0.046 | 0.066 | 0.080 |
| LBC1936 | MD | Clustering | -0.082 | 0.046 | 0.071 | 0.080 |
| Meta-analysis | SC | Mean edge weight | 0.201 | 0.037 | 0.000 | 0.000 |
| Meta-analysis | SC | Efficiency | 0.188 | 0.037 | 0.000 | 0.000 |
| Meta-analysis | SC | Clustering | 0.189 | 0.045 | 0.000 | 0.000 |
| Meta-analysis | FA | Mean edge weight | 0.155 | 0.068 | 0.023 | 0.027 |
| Meta-analysis | FA | Efficiency | 0.149 | 0.072 | 0.039 | 0.039 |
| Meta-analysis | FA | Clustering | 0.140 | 0.062 | 0.024 | 0.027 |
| Meta-analysis | MD | Mean edge weight | -0.064 | 0.026 | 0.014 | 0.022 |
| Meta-analysis | MD | Efficiency | -0.112 | 0.040 | 0.005 | 0.011 |
| Meta-analysis | MD | Clustering | -0.106 | 0.043 | 0.013 | 0.022 |

**Supplementary Table 7**. Standardised associations between g and local efficiency in all cohorts, including the meta-analysis, for 85 FreeSurfer regions in uncorrected SC networks. SE and p-values were estimated by SEM, and q-values were adjusted by FDR.

|  |  | **UKB** | | | | **STRADL** | | | | **LBC1936** | | | | **Meta-analysis** | | | |
| --- | --- | --- | --- | --- | --- | --- | --- | --- | --- | --- | --- | --- | --- | --- | --- | --- | --- |
| **Hemisphere** | **Node** | **β** | ***SE*** | ***p*** | ***q*** | **β** | ***SE*** | ***p*** | ***q*** | **β** | ***SE*** | ***p*** | ***q*** | **β** | ***SE*** | ***p*** | ***q*** |
| Left | Thalamus | 0.172 | 0.009 | 0.000 | 0.000 | 0.216 | 0.037 | 0.000 | 0.000 | 0.158 | 0.044 | 0.000 | 0.005 | 0.174 | 0.008 | 0.000 | 0.000 |
| Left | Caudate | 0.056 | 0.008 | 0.000 | 0.000 | 0.181 | 0.037 | 0.000 | 0.000 | 0.007 | 0.045 | 0.879 | 0.937 | 0.081 | 0.049 | 0.097 | 0.106 |
| Left | Putamen | 0.139 | 0.009 | 0.000 | 0.000 | 0.217 | 0.037 | 0.000 | 0.000 | 0.084 | 0.045 | 0.060 | 0.147 | 0.148 | 0.033 | 0.000 | 0.000 |
| Left | Pallidum | 0.097 | 0.009 | 0.000 | 0.000 | 0.075 | 0.037 | 0.041 | 0.048 | 0.101 | 0.045 | 0.026 | 0.084 | 0.096 | 0.008 | 0.000 | 0.000 |
| Left | Hippocampus | 0.091 | 0.009 | 0.000 | 0.000 | 0.105 | 0.037 | 0.005 | 0.007 | 0.112 | 0.045 | 0.012 | 0.059 | 0.092 | 0.008 | 0.000 | 0.000 |
| Left | Amygdala | 0.061 | 0.008 | 0.000 | 0.000 | 0.050 | 0.037 | 0.172 | 0.179 | 0.072 | 0.045 | 0.111 | 0.219 | 0.061 | 0.008 | 0.000 | 0.000 |
| Left | Accumbens-area | 0.038 | 0.009 | 0.000 | 0.000 | 0.017 | 0.037 | 0.645 | 0.645 | 0.021 | 0.045 | 0.632 | 0.768 | 0.036 | 0.008 | 0.000 | 0.000 |
| Left | Ventral diencephalon | 0.079 | 0.008 | 0.000 | 0.000 | 0.058 | 0.041 | 0.155 | 0.165 | 0.077 | 0.045 | 0.086 | 0.191 | 0.078 | 0.008 | 0.000 | 0.000 |
| Left | Banks of superior temporal | 0.074 | 0.008 | 0.000 | 0.000 | 0.143 | 0.037 | 0.000 | 0.000 | 0.034 | 0.045 | 0.450 | 0.599 | 0.083 | 0.024 | 0.000 | 0.001 |
| Left | Caudal anterior cingulate | 0.063 | 0.008 | 0.000 | 0.000 | 0.143 | 0.037 | 0.000 | 0.000 | 0.017 | 0.045 | 0.706 | 0.822 | 0.075 | 0.031 | 0.016 | 0.020 |
| Left | Caudal middle frontal | 0.102 | 0.009 | 0.000 | 0.000 | 0.187 | 0.035 | 0.000 | 0.000 | 0.070 | 0.045 | 0.118 | 0.228 | 0.119 | 0.031 | 0.000 | 0.000 |
| Left | Cuneus | 0.043 | 0.008 | 0.000 | 0.000 | 0.119 | 0.036 | 0.001 | 0.001 | 0.023 | 0.045 | 0.610 | 0.768 | 0.059 | 0.025 | 0.018 | 0.022 |
| Left | Entorhinal | 0.087 | 0.008 | 0.000 | 0.000 | 0.137 | 0.037 | 0.000 | 0.000 | 0.087 | 0.045 | 0.052 | 0.137 | 0.090 | 0.009 | 0.000 | 0.000 |
| Left | Fusiform | 0.101 | 0.008 | 0.000 | 0.000 | 0.050 | 0.037 | 0.170 | 0.179 | 0.076 | 0.045 | 0.091 | 0.197 | 0.091 | 0.015 | 0.000 | 0.000 |
| Left | Inferior parietal | 0.142 | 0.008 | 0.000 | 0.000 | 0.167 | 0.036 | 0.000 | 0.000 | 0.081 | 0.045 | 0.071 | 0.166 | 0.141 | 0.008 | 0.000 | 0.000 |
| Left | Inferior temporal | 0.117 | 0.008 | 0.000 | 0.000 | 0.127 | 0.036 | 0.000 | 0.001 | 0.047 | 0.045 | 0.295 | 0.440 | 0.115 | 0.008 | 0.000 | 0.000 |
| Left | Isthmus cingulate | 0.060 | 0.009 | 0.000 | 0.000 | 0.177 | 0.037 | 0.000 | 0.000 | 0.062 | 0.046 | 0.174 | 0.321 | 0.097 | 0.038 | 0.011 | 0.014 |
| Left | Lateral occipital | 0.123 | 0.008 | 0.000 | 0.000 | 0.147 | 0.037 | 0.000 | 0.000 | 0.117 | 0.044 | 0.008 | 0.046 | 0.124 | 0.008 | 0.000 | 0.000 |
| Left | Lateral orbitofrontal | 0.082 | 0.009 | 0.000 | 0.000 | 0.205 | 0.038 | 0.000 | 0.000 | -0.051 | 0.045 | 0.255 | 0.428 | 0.080 | 0.071 | 0.260 | 0.273 |
| Left | Lingual | 0.057 | 0.008 | 0.000 | 0.000 | 0.118 | 0.039 | 0.002 | 0.003 | 0.058 | 0.045 | 0.203 | 0.360 | 0.065 | 0.015 | 0.000 | 0.000 |
| Left | Medial orbitofrontal | 0.058 | 0.008 | 0.000 | 0.000 | 0.118 | 0.037 | 0.001 | 0.002 | 0.014 | 0.046 | 0.754 | 0.855 | 0.063 | 0.018 | 0.000 | 0.001 |
| Left | Middle temporal | 0.113 | 0.009 | 0.000 | 0.000 | 0.188 | 0.038 | 0.000 | 0.000 | 0.094 | 0.045 | 0.037 | 0.112 | 0.127 | 0.023 | 0.000 | 0.000 |
| Left | Parahippocampal | 0.096 | 0.009 | 0.000 | 0.000 | 0.110 | 0.038 | 0.004 | 0.005 | 0.050 | 0.045 | 0.262 | 0.428 | 0.095 | 0.008 | 0.000 | 0.000 |
| Left | Paracentral | 0.081 | 0.008 | 0.000 | 0.000 | 0.164 | 0.035 | 0.000 | 0.000 | 0.109 | 0.044 | 0.014 | 0.064 | 0.110 | 0.027 | 0.000 | 0.000 |
| Left | Pars opercularis | 0.102 | 0.009 | 0.000 | 0.000 | 0.186 | 0.038 | 0.000 | 0.000 | 0.044 | 0.046 | 0.335 | 0.483 | 0.112 | 0.035 | 0.002 | 0.002 |
| Left | Pars orbitalis | 0.069 | 0.009 | 0.000 | 0.000 | 0.093 | 0.037 | 0.013 | 0.016 | 0.008 | 0.045 | 0.857 | 0.934 | 0.068 | 0.008 | 0.000 | 0.000 |
| Left | Pars triangularis | 0.099 | 0.009 | 0.000 | 0.000 | 0.159 | 0.037 | 0.000 | 0.000 | 0.049 | 0.045 | 0.280 | 0.440 | 0.104 | 0.021 | 0.000 | 0.000 |
| Left | Pericalcarine | 0.027 | 0.008 | 0.001 | 0.001 | 0.079 | 0.036 | 0.028 | 0.033 | -0.016 | 0.045 | 0.722 | 0.830 | 0.028 | 0.008 | 0.001 | 0.001 |
| Left | Postcentral | 0.121 | 0.009 | 0.000 | 0.000 | 0.181 | 0.036 | 0.000 | 0.000 | 0.125 | 0.046 | 0.007 | 0.040 | 0.133 | 0.017 | 0.000 | 0.000 |
| Left | Posterior cingulate | 0.064 | 0.008 | 0.000 | 0.000 | 0.120 | 0.037 | 0.001 | 0.002 | 0.079 | 0.045 | 0.075 | 0.173 | 0.075 | 0.016 | 0.000 | 0.000 |
| Left | Precentral | 0.127 | 0.009 | 0.000 | 0.000 | 0.274 | 0.035 | 0.000 | 0.000 | 0.131 | 0.045 | 0.003 | 0.029 | 0.176 | 0.048 | 0.000 | 0.000 |
| Left | Precuneus | 0.112 | 0.008 | 0.000 | 0.000 | 0.206 | 0.035 | 0.000 | 0.000 | 0.088 | 0.045 | 0.050 | 0.136 | 0.135 | 0.033 | 0.000 | 0.000 |
| Left | Rostral anterior cingulate | 0.059 | 0.008 | 0.000 | 0.000 | 0.137 | 0.037 | 0.000 | 0.000 | -0.007 | 0.045 | 0.882 | 0.937 | 0.065 | 0.036 | 0.068 | 0.076 |
| Left | Rostral middle frontal | 0.106 | 0.009 | 0.000 | 0.000 | 0.239 | 0.037 | 0.000 | 0.000 | 0.003 | 0.045 | 0.949 | 0.974 | 0.117 | 0.066 | 0.074 | 0.081 |
| Left | Superior frontal | 0.142 | 0.009 | 0.000 | 0.000 | 0.234 | 0.036 | 0.000 | 0.000 | 0.090 | 0.045 | 0.045 | 0.132 | 0.156 | 0.038 | 0.000 | 0.000 |
| Left | Superior parietal | 0.121 | 0.009 | 0.000 | 0.000 | 0.188 | 0.036 | 0.000 | 0.000 | 0.116 | 0.044 | 0.009 | 0.046 | 0.135 | 0.020 | 0.000 | 0.000 |
| Left | Superior temporal | 0.123 | 0.009 | 0.000 | 0.000 | 0.236 | 0.037 | 0.000 | 0.000 | 0.022 | 0.046 | 0.627 | 0.768 | 0.129 | 0.058 | 0.027 | 0.032 |
| Left | Supramarginal | 0.122 | 0.009 | 0.000 | 0.000 | 0.198 | 0.037 | 0.000 | 0.000 | 0.074 | 0.045 | 0.099 | 0.207 | 0.132 | 0.030 | 0.000 | 0.000 |
| Left | Frontal pole | 0.070 | 0.008 | 0.000 | 0.000 | 0.158 | 0.040 | 0.000 | 0.000 | -0.042 | 0.045 | 0.349 | 0.495 | 0.064 | 0.054 | 0.236 | 0.251 |
| Left | Temporal pole | 0.060 | 0.008 | 0.000 | 0.000 | 0.089 | 0.038 | 0.019 | 0.023 | -0.013 | 0.046 | 0.768 | 0.859 | 0.059 | 0.008 | 0.000 | 0.000 |
| Left | Transverse temporal | 0.081 | 0.008 | 0.000 | 0.000 | 0.136 | 0.036 | 0.000 | 0.000 | 0.001 | 0.045 | 0.986 | 0.986 | 0.078 | 0.032 | 0.015 | 0.019 |
| Left | Insula | 0.091 | 0.008 | 0.000 | 0.000 | 0.148 | 0.036 | 0.000 | 0.000 | -0.050 | 0.045 | 0.267 | 0.428 | 0.067 | 0.055 | 0.224 | 0.241 |
| Right | Thalamus | 0.187 | 0.009 | 0.000 | 0.000 | 0.254 | 0.037 | 0.000 | 0.000 | 0.168 | 0.044 | 0.000 | 0.003 | 0.198 | 0.020 | 0.000 | 0.000 |
| Right | Caudate | 0.045 | 0.009 | 0.000 | 0.000 | 0.102 | 0.039 | 0.008 | 0.010 | 0.018 | 0.046 | 0.694 | 0.819 | 0.046 | 0.009 | 0.000 | 0.000 |
| Right | Putamen | 0.154 | 0.009 | 0.000 | 0.000 | 0.243 | 0.036 | 0.000 | 0.000 | 0.160 | 0.044 | 0.000 | 0.005 | 0.180 | 0.028 | 0.000 | 0.000 |
| Right | Pallidum | 0.146 | 0.009 | 0.000 | 0.000 | 0.100 | 0.036 | 0.006 | 0.008 | 0.123 | 0.045 | 0.006 | 0.040 | 0.139 | 0.013 | 0.000 | 0.000 |
| Right | Hippocampus | 0.081 | 0.009 | 0.000 | 0.000 | 0.136 | 0.036 | 0.000 | 0.000 | 0.173 | 0.044 | 0.000 | 0.002 | 0.118 | 0.029 | 0.000 | 0.000 |
| Right | Amygdala | 0.039 | 0.009 | 0.000 | 0.000 | 0.041 | 0.040 | 0.305 | 0.312 | -0.059 | 0.045 | 0.192 | 0.347 | 0.018 | 0.027 | 0.497 | 0.503 |
| Right | Accumbens-area | 0.064 | 0.009 | 0.000 | 0.000 | 0.064 | 0.036 | 0.079 | 0.085 | 0.034 | 0.045 | 0.451 | 0.599 | 0.063 | 0.008 | 0.000 | 0.000 |
| Right | Ventral diencephalon | 0.096 | 0.008 | 0.000 | 0.000 | 0.103 | 0.037 | 0.005 | 0.007 | 0.089 | 0.045 | 0.047 | 0.134 | 0.096 | 0.008 | 0.000 | 0.000 |
| Right | Banks of superior temporal | 0.047 | 0.009 | 0.000 | 0.000 | 0.074 | 0.037 | 0.047 | 0.053 | 0.100 | 0.045 | 0.024 | 0.084 | 0.053 | 0.012 | 0.000 | 0.000 |
| Right | Caudal anterior cingulate | 0.042 | 0.008 | 0.000 | 0.000 | 0.145 | 0.036 | 0.000 | 0.000 | 0.022 | 0.045 | 0.620 | 0.768 | 0.068 | 0.035 | 0.055 | 0.064 |
| Right | Caudal middle frontal | 0.084 | 0.009 | 0.000 | 0.000 | 0.206 | 0.034 | 0.000 | 0.000 | 0.140 | 0.044 | 0.002 | 0.015 | 0.137 | 0.038 | 0.000 | 0.000 |
| Right | Cuneus | 0.027 | 0.008 | 0.001 | 0.001 | 0.145 | 0.036 | 0.000 | 0.000 | -0.050 | 0.045 | 0.263 | 0.428 | 0.042 | 0.054 | 0.430 | 0.440 |
| Right | Entorhinal | 0.064 | 0.008 | 0.000 | 0.000 | 0.064 | 0.036 | 0.076 | 0.083 | 0.048 | 0.045 | 0.294 | 0.440 | 0.064 | 0.008 | 0.000 | 0.000 |
| Right | Fusiform | 0.082 | 0.008 | 0.000 | 0.000 | 0.122 | 0.038 | 0.001 | 0.002 | 0.099 | 0.045 | 0.026 | 0.084 | 0.085 | 0.008 | 0.000 | 0.000 |
| Right | Inferior parietal | 0.130 | 0.008 | 0.000 | 0.000 | 0.202 | 0.035 | 0.000 | 0.000 | 0.105 | 0.044 | 0.018 | 0.076 | 0.144 | 0.024 | 0.000 | 0.000 |
| Right | Inferior temporal | 0.107 | 0.008 | 0.000 | 0.000 | 0.142 | 0.036 | 0.000 | 0.000 | 0.086 | 0.045 | 0.053 | 0.137 | 0.108 | 0.008 | 0.000 | 0.000 |
| Right | Isthmus cingulate | 0.050 | 0.009 | 0.000 | 0.000 | 0.107 | 0.038 | 0.006 | 0.007 | 0.042 | 0.045 | 0.359 | 0.501 | 0.054 | 0.011 | 0.000 | 0.000 |
| Right | Lateral occipital | 0.099 | 0.008 | 0.000 | 0.000 | 0.182 | 0.037 | 0.000 | 0.000 | 0.035 | 0.045 | 0.436 | 0.598 | 0.107 | 0.037 | 0.004 | 0.005 |
| Right | Lateral orbitofrontal | 0.086 | 0.009 | 0.000 | 0.000 | 0.145 | 0.038 | 0.000 | 0.000 | -0.002 | 0.045 | 0.962 | 0.974 | 0.081 | 0.036 | 0.027 | 0.032 |
| Right | Lingual | 0.047 | 0.008 | 0.000 | 0.000 | 0.145 | 0.037 | 0.000 | 0.000 | 0.046 | 0.045 | 0.309 | 0.452 | 0.075 | 0.031 | 0.017 | 0.021 |
| Right | Medial orbitofrontal | 0.073 | 0.008 | 0.000 | 0.000 | 0.099 | 0.036 | 0.006 | 0.008 | 0.011 | 0.046 | 0.803 | 0.886 | 0.072 | 0.008 | 0.000 | 0.000 |
| Right | Middle temporal | 0.115 | 0.008 | 0.000 | 0.000 | 0.192 | 0.037 | 0.000 | 0.000 | 0.142 | 0.045 | 0.001 | 0.015 | 0.139 | 0.024 | 0.000 | 0.000 |
| Right | Parahippocampal | 0.085 | 0.008 | 0.000 | 0.000 | 0.128 | 0.036 | 0.000 | 0.001 | 0.099 | 0.045 | 0.026 | 0.084 | 0.088 | 0.008 | 0.000 | 0.000 |
| Right | Paracentral | 0.094 | 0.008 | 0.000 | 0.000 | 0.133 | 0.036 | 0.000 | 0.000 | 0.121 | 0.044 | 0.006 | 0.040 | 0.098 | 0.010 | 0.000 | 0.000 |
| Right | Pars opercularis | 0.107 | 0.009 | 0.000 | 0.000 | 0.157 | 0.037 | 0.000 | 0.000 | 0.071 | 0.045 | 0.110 | 0.219 | 0.108 | 0.008 | 0.000 | 0.000 |
| Right | Pars orbitalis | 0.120 | 0.009 | 0.000 | 0.000 | 0.131 | 0.038 | 0.000 | 0.001 | 0.019 | 0.045 | 0.674 | 0.807 | 0.100 | 0.029 | 0.000 | 0.001 |
| Right | Pars triangularis | 0.094 | 0.009 | 0.000 | 0.000 | 0.148 | 0.037 | 0.000 | 0.000 | 0.117 | 0.045 | 0.009 | 0.046 | 0.106 | 0.016 | 0.000 | 0.000 |
| Right | Pericalcarine | 0.018 | 0.008 | 0.033 | 0.033 | 0.067 | 0.036 | 0.063 | 0.070 | 0.047 | 0.045 | 0.294 | 0.440 | 0.029 | 0.016 | 0.068 | 0.076 |
| Right | Postcentral | 0.129 | 0.008 | 0.000 | 0.000 | 0.179 | 0.035 | 0.000 | 0.000 | 0.205 | 0.044 | 0.000 | 0.000 | 0.157 | 0.025 | 0.000 | 0.000 |
| Right | Posterior cingulate | 0.077 | 0.008 | 0.000 | 0.000 | 0.131 | 0.038 | 0.000 | 0.001 | 0.030 | 0.045 | 0.503 | 0.658 | 0.078 | 0.008 | 0.000 | 0.000 |
| Right | Precentral | 0.126 | 0.009 | 0.000 | 0.000 | 0.214 | 0.036 | 0.000 | 0.000 | 0.201 | 0.043 | 0.000 | 0.000 | 0.171 | 0.032 | 0.000 | 0.000 |
| Right | Precuneus | 0.123 | 0.008 | 0.000 | 0.000 | 0.186 | 0.035 | 0.000 | 0.000 | 0.100 | 0.044 | 0.025 | 0.084 | 0.132 | 0.018 | 0.000 | 0.000 |
| Right | Rostral anterior cingulate | 0.052 | 0.008 | 0.000 | 0.000 | 0.164 | 0.039 | 0.000 | 0.000 | -0.053 | 0.045 | 0.243 | 0.422 | 0.056 | 0.059 | 0.346 | 0.358 |
| Right | Rostral middle frontal | 0.120 | 0.009 | 0.000 | 0.000 | 0.241 | 0.039 | 0.000 | 0.000 | 0.063 | 0.045 | 0.159 | 0.300 | 0.142 | 0.049 | 0.004 | 0.005 |
| Right | Superior frontal | 0.136 | 0.009 | 0.000 | 0.000 | 0.256 | 0.035 | 0.000 | 0.000 | 0.153 | 0.044 | 0.000 | 0.006 | 0.178 | 0.038 | 0.000 | 0.000 |
| Right | Superior parietal | 0.131 | 0.009 | 0.000 | 0.000 | 0.210 | 0.035 | 0.000 | 0.000 | 0.127 | 0.044 | 0.004 | 0.032 | 0.151 | 0.025 | 0.000 | 0.000 |
| Right | Superior temporal | 0.129 | 0.009 | 0.000 | 0.000 | 0.152 | 0.035 | 0.000 | 0.000 | 0.101 | 0.046 | 0.027 | 0.084 | 0.129 | 0.008 | 0.000 | 0.000 |
| Right | Supramarginal | 0.095 | 0.009 | 0.000 | 0.000 | 0.108 | 0.036 | 0.003 | 0.004 | 0.027 | 0.046 | 0.554 | 0.714 | 0.093 | 0.008 | 0.000 | 0.000 |
| Right | Frontal pole | 0.060 | 0.008 | 0.000 | 0.000 | 0.156 | 0.038 | 0.000 | 0.000 | 0.002 | 0.045 | 0.961 | 0.974 | 0.074 | 0.040 | 0.068 | 0.076 |
| Right | Temporal pole | 0.046 | 0.008 | 0.000 | 0.000 | -0.026 | 0.036 | 0.466 | 0.471 | -0.099 | 0.045 | 0.026 | 0.084 | -0.018 | 0.042 | 0.674 | 0.674 |
| Right | Transverse temporal | 0.068 | 0.008 | 0.000 | 0.000 | 0.078 | 0.037 | 0.032 | 0.037 | -0.006 | 0.047 | 0.897 | 0.941 | 0.067 | 0.008 | 0.000 | 0.000 |
| Right | Insula | 0.088 | 0.009 | 0.000 | 0.000 | 0.215 | 0.036 | 0.000 | 0.000 | 0.084 | 0.045 | 0.061 | 0.147 | 0.127 | 0.042 | 0.003 | 0.004 |
|  | Brainstem | 0.044 | 0.009 | 0.000 | 0.000 | 0.069 | 0.037 | 0.060 | 0.067 | 0.074 | 0.045 | 0.100 | 0.207 | 0.046 | 0.008 | 0.000 | 0.000 |

**Supplementary Table 8**. Standardised associations between g and local efficiency in all cohorts, including the meta-analysis, for 85 FreeSurfer regions in volume corrected SD networks. SE and p-values were estimated by SEM, and q-values were adjusted by FDR.

|  |  | **UKB** | | | | **STRADL** | | | | **LBC1936** | | | | **Meta-analysis** | | | |
| --- | --- | --- | --- | --- | --- | --- | --- | --- | --- | --- | --- | --- | --- | --- | --- | --- | --- |
| **Hemisphere** | **Node** | **β** | ***SE*** | ***p*** | ***q*** | **β** | ***SE*** | ***p*** | ***q*** | **β** | ***SE*** | ***p*** | ***q*** | **β** | ***SE*** | ***p*** | ***q*** |
| Left | Thalamus | 0.069 | 0.009 | 0.000 | 0.000 | 0.085 | 0.037 | 0.021 | 0.047 | -0.000 | 0.045 | 0.998 | 0.998 | 0.068 | 0.008 | 0.000 | 0.000 |
| Left | Caudate | -0.008 | 0.008 | 0.347 | 0.404 | 0.060 | 0.037 | 0.101 | 0.156 | -0.105 | 0.045 | 0.019 | 0.317 | -0.014 | 0.043 | 0.746 | 0.828 |
| Left | Putamen | 0.021 | 0.008 | 0.015 | 0.025 | 0.089 | 0.036 | 0.014 | 0.037 | -0.048 | 0.045 | 0.288 | 0.847 | 0.024 | 0.033 | 0.469 | 0.594 |
| Left | Pallidum | 0.019 | 0.009 | 0.027 | 0.044 | 0.005 | 0.037 | 0.902 | 0.924 | -0.043 | 0.045 | 0.344 | 0.867 | 0.016 | 0.009 | 0.094 | 0.207 |
| Left | Hippocampus | 0.037 | 0.009 | 0.000 | 0.000 | 0.045 | 0.038 | 0.233 | 0.315 | 0.037 | 0.046 | 0.426 | 0.867 | 0.037 | 0.008 | 0.000 | 0.000 |
| Left | Amygdala | -0.002 | 0.008 | 0.796 | 0.836 | -0.023 | 0.036 | 0.529 | 0.583 | -0.008 | 0.045 | 0.858 | 0.976 | -0.003 | 0.008 | 0.677 | 0.777 |
| Left | Accumbens-area | -0.013 | 0.008 | 0.113 | 0.159 | -0.048 | 0.037 | 0.195 | 0.268 | -0.034 | 0.045 | 0.451 | 0.867 | -0.016 | 0.008 | 0.052 | 0.127 |
| Left | Ventral diencephalon | -0.007 | 0.008 | 0.415 | 0.473 | -0.032 | 0.042 | 0.445 | 0.504 | -0.061 | 0.045 | 0.174 | 0.773 | -0.012 | 0.011 | 0.293 | 0.429 |
| Left | Banks of superior temporal | 0.013 | 0.008 | 0.127 | 0.169 | 0.080 | 0.038 | 0.037 | 0.072 | -0.035 | 0.045 | 0.438 | 0.867 | 0.019 | 0.024 | 0.414 | 0.559 |
| Left | Caudal anterior cingulate | 0.002 | 0.008 | 0.824 | 0.839 | 0.081 | 0.037 | 0.028 | 0.062 | -0.015 | 0.045 | 0.739 | 0.976 | 0.019 | 0.025 | 0.457 | 0.594 |
| Left | Caudal middle frontal | 0.042 | 0.008 | 0.000 | 0.000 | 0.134 | 0.035 | 0.000 | 0.001 | 0.021 | 0.045 | 0.644 | 0.976 | 0.064 | 0.032 | 0.042 | 0.105 |
| Left | Cuneus | 0.002 | 0.008 | 0.829 | 0.839 | 0.076 | 0.036 | 0.037 | 0.072 | -0.038 | 0.046 | 0.408 | 0.867 | 0.014 | 0.027 | 0.612 | 0.713 |
| Left | Entorhinal | 0.047 | 0.008 | 0.000 | 0.000 | 0.092 | 0.036 | 0.011 | 0.033 | 0.003 | 0.045 | 0.941 | 0.976 | 0.048 | 0.008 | 0.000 | 0.000 |
| Left | Fusiform | 0.031 | 0.009 | 0.000 | 0.001 | -0.036 | 0.037 | 0.339 | 0.412 | -0.034 | 0.046 | 0.459 | 0.867 | 0.000 | 0.026 | 0.988 | 0.988 |
| Left | Inferior parietal | 0.081 | 0.008 | 0.000 | 0.000 | 0.086 | 0.036 | 0.018 | 0.043 | -0.004 | 0.045 | 0.936 | 0.976 | 0.072 | 0.017 | 0.000 | 0.000 |
| Left | Inferior temporal | 0.040 | 0.008 | 0.000 | 0.000 | 0.055 | 0.037 | 0.136 | 0.196 | -0.047 | 0.045 | 0.299 | 0.847 | 0.030 | 0.020 | 0.125 | 0.231 |
| Left | Isthmus cingulate | -0.016 | 0.009 | 0.072 | 0.111 | 0.097 | 0.039 | 0.014 | 0.037 | -0.036 | 0.045 | 0.425 | 0.867 | 0.012 | 0.038 | 0.747 | 0.828 |
| Left | Lateral occipital | 0.063 | 0.008 | 0.000 | 0.000 | 0.072 | 0.040 | 0.068 | 0.123 | 0.047 | 0.045 | 0.293 | 0.847 | 0.063 | 0.008 | 0.000 | 0.000 |
| Left | Lateral orbitofrontal | 0.006 | 0.008 | 0.480 | 0.530 | 0.104 | 0.039 | 0.008 | 0.027 | -0.098 | 0.045 | 0.030 | 0.317 | 0.006 | 0.054 | 0.917 | 0.939 |
| Left | Lingual | 0.011 | 0.008 | 0.204 | 0.258 | 0.071 | 0.041 | 0.084 | 0.147 | -0.035 | 0.046 | 0.448 | 0.867 | 0.012 | 0.008 | 0.155 | 0.268 |
| Left | Medial orbitofrontal | -0.000 | 0.008 | 0.983 | 0.983 | 0.041 | 0.037 | 0.269 | 0.357 | -0.017 | 0.047 | 0.719 | 0.976 | 0.001 | 0.008 | 0.872 | 0.938 |
| Left | Middle temporal | 0.042 | 0.008 | 0.000 | 0.000 | 0.104 | 0.042 | 0.012 | 0.034 | 0.004 | 0.045 | 0.928 | 0.976 | 0.043 | 0.008 | 0.000 | 0.000 |
| Left | Parahippocampal | 0.050 | 0.009 | 0.000 | 0.000 | 0.058 | 0.038 | 0.132 | 0.196 | -0.025 | 0.045 | 0.572 | 0.934 | 0.048 | 0.008 | 0.000 | 0.000 |
| Left | Paracentral | 0.011 | 0.008 | 0.209 | 0.258 | 0.087 | 0.036 | 0.017 | 0.042 | 0.004 | 0.045 | 0.936 | 0.976 | 0.028 | 0.024 | 0.234 | 0.349 |
| Left | Pars opercularis | 0.039 | 0.008 | 0.000 | 0.000 | 0.107 | 0.036 | 0.003 | 0.012 | -0.031 | 0.046 | 0.500 | 0.904 | 0.042 | 0.033 | 0.202 | 0.312 |
| Left | Pars orbitalis | 0.013 | 0.008 | 0.115 | 0.160 | 0.020 | 0.036 | 0.574 | 0.618 | -0.040 | 0.045 | 0.378 | 0.867 | 0.012 | 0.008 | 0.139 | 0.246 |
| Left | Pars triangularis | 0.034 | 0.008 | 0.000 | 0.000 | 0.091 | 0.036 | 0.011 | 0.033 | -0.010 | 0.045 | 0.830 | 0.976 | 0.039 | 0.017 | 0.019 | 0.057 |
| Left | Pericalcarine | -0.011 | 0.008 | 0.209 | 0.258 | 0.038 | 0.037 | 0.303 | 0.391 | -0.070 | 0.045 | 0.120 | 0.773 | -0.010 | 0.008 | 0.216 | 0.327 |
| Left | Postcentral | 0.047 | 0.008 | 0.000 | 0.000 | 0.105 | 0.036 | 0.004 | 0.013 | 0.012 | 0.045 | 0.792 | 0.976 | 0.052 | 0.015 | 0.001 | 0.003 |
| Left | Posterior cingulate | -0.010 | 0.008 | 0.248 | 0.301 | 0.052 | 0.037 | 0.154 | 0.219 | 0.012 | 0.045 | 0.784 | 0.976 | 0.006 | 0.019 | 0.750 | 0.828 |
| Left | Precentral | 0.047 | 0.008 | 0.000 | 0.000 | 0.186 | 0.035 | 0.000 | 0.000 | 0.017 | 0.045 | 0.699 | 0.976 | 0.083 | 0.050 | 0.098 | 0.207 |
| Left | Precuneus | 0.032 | 0.008 | 0.000 | 0.000 | 0.119 | 0.038 | 0.002 | 0.010 | -0.041 | 0.046 | 0.375 | 0.867 | 0.038 | 0.041 | 0.347 | 0.491 |
| Left | Rostral anterior cingulate | 0.004 | 0.008 | 0.588 | 0.633 | 0.062 | 0.037 | 0.094 | 0.150 | -0.016 | 0.046 | 0.724 | 0.976 | 0.009 | 0.013 | 0.470 | 0.594 |
| Left | Rostral middle frontal | 0.042 | 0.008 | 0.000 | 0.000 | 0.150 | 0.037 | 0.000 | 0.001 | -0.044 | 0.046 | 0.332 | 0.867 | 0.051 | 0.053 | 0.329 | 0.475 |
| Left | Superior frontal | 0.063 | 0.009 | 0.000 | 0.000 | 0.137 | 0.036 | 0.000 | 0.001 | -0.022 | 0.046 | 0.629 | 0.976 | 0.063 | 0.041 | 0.123 | 0.231 |
| Left | Superior parietal | 0.048 | 0.008 | 0.000 | 0.000 | 0.108 | 0.035 | 0.002 | 0.011 | 0.026 | 0.045 | 0.568 | 0.934 | 0.056 | 0.017 | 0.001 | 0.005 |
| Left | Superior temporal | 0.034 | 0.008 | 0.000 | 0.000 | 0.159 | 0.040 | 0.000 | 0.001 | -0.068 | 0.045 | 0.134 | 0.773 | 0.043 | 0.063 | 0.494 | 0.599 |
| Left | Supramarginal | 0.050 | 0.008 | 0.000 | 0.000 | 0.131 | 0.037 | 0.000 | 0.003 | -0.005 | 0.045 | 0.920 | 0.976 | 0.060 | 0.034 | 0.074 | 0.176 |
| Left | Frontal pole | 0.034 | 0.008 | 0.000 | 0.000 | 0.095 | 0.041 | 0.021 | 0.047 | -0.060 | 0.046 | 0.195 | 0.773 | 0.027 | 0.038 | 0.489 | 0.599 |
| Left | Temporal pole | 0.013 | 0.008 | 0.124 | 0.169 | 0.023 | 0.037 | 0.535 | 0.583 | -0.064 | 0.045 | 0.153 | 0.773 | 0.011 | 0.008 | 0.180 | 0.289 |
| Left | Transverse temporal | 0.030 | 0.008 | 0.000 | 0.001 | 0.068 | 0.037 | 0.063 | 0.117 | -0.044 | 0.045 | 0.331 | 0.867 | 0.030 | 0.008 | 0.000 | 0.001 |
| Left | Insula | 0.017 | 0.008 | 0.036 | 0.058 | 0.076 | 0.037 | 0.037 | 0.072 | -0.117 | 0.045 | 0.009 | 0.226 | -0.004 | 0.053 | 0.943 | 0.954 |
| Right | Thalamus | 0.108 | 0.008 | 0.000 | 0.000 | 0.162 | 0.037 | 0.000 | 0.000 | 0.054 | 0.045 | 0.227 | 0.803 | 0.110 | 0.014 | 0.000 | 0.000 |
| Right | Caudate | 0.015 | 0.009 | 0.084 | 0.126 | 0.039 | 0.039 | 0.314 | 0.393 | -0.052 | 0.047 | 0.267 | 0.841 | 0.014 | 0.008 | 0.102 | 0.207 |
| Right | Putamen | 0.040 | 0.009 | 0.000 | 0.000 | 0.106 | 0.037 | 0.004 | 0.015 | 0.016 | 0.046 | 0.727 | 0.976 | 0.050 | 0.019 | 0.010 | 0.033 |
| Right | Pallidum | 0.084 | 0.008 | 0.000 | 0.000 | 0.036 | 0.036 | 0.314 | 0.393 | -0.027 | 0.045 | 0.551 | 0.934 | 0.043 | 0.032 | 0.180 | 0.289 |
| Right | Hippocampus | 0.030 | 0.009 | 0.000 | 0.001 | 0.065 | 0.038 | 0.084 | 0.147 | 0.073 | 0.045 | 0.105 | 0.773 | 0.036 | 0.011 | 0.002 | 0.007 |
| Right | Amygdala | -0.008 | 0.008 | 0.320 | 0.378 | -0.036 | 0.038 | 0.345 | 0.413 | -0.128 | 0.044 | 0.004 | 0.160 | -0.048 | 0.034 | 0.164 | 0.279 |
| Right | Accumbens-area | 0.017 | 0.008 | 0.040 | 0.063 | 0.019 | 0.037 | 0.612 | 0.651 | -0.001 | 0.046 | 0.980 | 0.992 | 0.017 | 0.008 | 0.038 | 0.097 |
| Right | Ventral diencephalon | 0.007 | 0.009 | 0.417 | 0.473 | -0.004 | 0.038 | 0.917 | 0.928 | -0.065 | 0.045 | 0.146 | 0.773 | -0.002 | 0.015 | 0.916 | 0.939 |
| Right | Banks of superior temporal | -0.020 | 0.009 | 0.021 | 0.035 | -0.031 | 0.039 | 0.434 | 0.498 | 0.019 | 0.045 | 0.671 | 0.976 | -0.019 | 0.008 | 0.021 | 0.060 |
| Right | Caudal anterior cingulate | 0.003 | 0.008 | 0.715 | 0.760 | 0.080 | 0.038 | 0.035 | 0.072 | -0.007 | 0.046 | 0.886 | 0.976 | 0.020 | 0.024 | 0.407 | 0.559 |
| Right | Caudal middle frontal | 0.014 | 0.008 | 0.101 | 0.145 | 0.137 | 0.035 | 0.000 | 0.001 | 0.059 | 0.045 | 0.191 | 0.773 | 0.064 | 0.038 | 0.093 | 0.207 |
| Right | Cuneus | -0.006 | 0.008 | 0.492 | 0.537 | 0.120 | 0.037 | 0.001 | 0.008 | -0.098 | 0.045 | 0.028 | 0.317 | 0.007 | 0.060 | 0.914 | 0.939 |
| Right | Entorhinal | 0.034 | 0.008 | 0.000 | 0.000 | 0.001 | 0.036 | 0.984 | 0.984 | -0.020 | 0.045 | 0.655 | 0.976 | 0.024 | 0.015 | 0.112 | 0.216 |
| Right | Fusiform | 0.019 | 0.009 | 0.023 | 0.038 | 0.036 | 0.038 | 0.339 | 0.412 | 0.013 | 0.045 | 0.776 | 0.976 | 0.020 | 0.008 | 0.014 | 0.044 |
| Right | Inferior parietal | 0.059 | 0.008 | 0.000 | 0.000 | 0.110 | 0.036 | 0.002 | 0.011 | -0.007 | 0.045 | 0.869 | 0.976 | 0.059 | 0.023 | 0.009 | 0.031 |
| Right | Inferior temporal | 0.033 | 0.008 | 0.000 | 0.000 | 0.066 | 0.040 | 0.095 | 0.150 | -0.018 | 0.046 | 0.694 | 0.976 | 0.033 | 0.008 | 0.000 | 0.000 |
| Right | Isthmus cingulate | -0.015 | 0.009 | 0.088 | 0.130 | 0.044 | 0.041 | 0.287 | 0.375 | -0.051 | 0.045 | 0.254 | 0.830 | -0.014 | 0.008 | 0.102 | 0.207 |
| Right | Lateral occipital | 0.032 | 0.008 | 0.000 | 0.000 | 0.133 | 0.042 | 0.002 | 0.009 | -0.058 | 0.045 | 0.200 | 0.773 | 0.036 | 0.051 | 0.475 | 0.594 |
| Right | Lateral orbitofrontal | 0.011 | 0.008 | 0.196 | 0.253 | 0.050 | 0.039 | 0.194 | 0.268 | -0.062 | 0.046 | 0.181 | 0.773 | 0.010 | 0.008 | 0.201 | 0.312 |
| Right | Lingual | 0.006 | 0.008 | 0.478 | 0.530 | 0.097 | 0.039 | 0.014 | 0.037 | -0.039 | 0.046 | 0.389 | 0.867 | 0.021 | 0.035 | 0.549 | 0.657 |
| Right | Medial orbitofrontal | 0.021 | 0.008 | 0.013 | 0.024 | 0.030 | 0.037 | 0.421 | 0.497 | 0.027 | 0.046 | 0.560 | 0.934 | 0.022 | 0.008 | 0.008 | 0.030 |
| Right | Middle temporal | 0.040 | 0.008 | 0.000 | 0.000 | 0.113 | 0.041 | 0.006 | 0.020 | 0.038 | 0.045 | 0.399 | 0.867 | 0.052 | 0.019 | 0.007 | 0.027 |
| Right | Parahippocampal | 0.039 | 0.008 | 0.000 | 0.000 | 0.055 | 0.037 | 0.134 | 0.196 | 0.004 | 0.045 | 0.921 | 0.976 | 0.039 | 0.008 | 0.000 | 0.000 |
| Right | Paracentral | 0.034 | 0.008 | 0.000 | 0.000 | 0.078 | 0.036 | 0.032 | 0.068 | 0.061 | 0.046 | 0.182 | 0.773 | 0.041 | 0.012 | 0.001 | 0.004 |
| Right | Pars opercularis | 0.054 | 0.008 | 0.000 | 0.000 | 0.074 | 0.037 | 0.044 | 0.084 | 0.004 | 0.045 | 0.936 | 0.976 | 0.053 | 0.008 | 0.000 | 0.000 |
| Right | Pars orbitalis | 0.061 | 0.008 | 0.000 | 0.000 | 0.061 | 0.037 | 0.093 | 0.150 | -0.028 | 0.045 | 0.540 | 0.934 | 0.047 | 0.021 | 0.026 | 0.070 |
| Right | Pars triangularis | 0.031 | 0.008 | 0.000 | 0.001 | 0.062 | 0.037 | 0.090 | 0.150 | 0.053 | 0.045 | 0.240 | 0.816 | 0.033 | 0.008 | 0.000 | 0.000 |
| Right | Pericalcarine | -0.014 | 0.008 | 0.084 | 0.126 | 0.029 | 0.037 | 0.431 | 0.498 | -0.005 | 0.045 | 0.917 | 0.976 | -0.012 | 0.008 | 0.133 | 0.241 |
| Right | Postcentral | 0.070 | 0.008 | 0.000 | 0.000 | 0.107 | 0.036 | 0.003 | 0.012 | 0.113 | 0.044 | 0.011 | 0.226 | 0.078 | 0.014 | 0.000 | 0.000 |
| Right | Posterior cingulate | 0.021 | 0.008 | 0.011 | 0.020 | 0.061 | 0.038 | 0.109 | 0.166 | -0.036 | 0.045 | 0.424 | 0.867 | 0.021 | 0.008 | 0.010 | 0.034 |
| Right | Precentral | 0.049 | 0.008 | 0.000 | 0.000 | 0.131 | 0.036 | 0.000 | 0.002 | 0.087 | 0.045 | 0.051 | 0.478 | 0.080 | 0.027 | 0.003 | 0.013 |
| Right | Precuneus | 0.056 | 0.008 | 0.000 | 0.000 | 0.117 | 0.038 | 0.002 | 0.010 | -0.006 | 0.046 | 0.902 | 0.976 | 0.059 | 0.026 | 0.023 | 0.064 |
| Right | Rostral anterior cingulate | 0.012 | 0.008 | 0.164 | 0.214 | 0.149 | 0.040 | 0.000 | 0.002 | -0.065 | 0.046 | 0.154 | 0.773 | 0.032 | 0.059 | 0.591 | 0.698 |
| Right | Rostral middle frontal | 0.056 | 0.008 | 0.000 | 0.000 | 0.146 | 0.041 | 0.000 | 0.002 | -0.011 | 0.047 | 0.821 | 0.976 | 0.065 | 0.040 | 0.101 | 0.207 |
| Right | Superior frontal | 0.057 | 0.009 | 0.000 | 0.000 | 0.158 | 0.036 | 0.000 | 0.000 | 0.061 | 0.045 | 0.177 | 0.773 | 0.088 | 0.033 | 0.007 | 0.027 |
| Right | Superior parietal | 0.062 | 0.008 | 0.000 | 0.000 | 0.133 | 0.035 | 0.000 | 0.001 | 0.012 | 0.045 | 0.795 | 0.976 | 0.071 | 0.029 | 0.014 | 0.043 |
| Right | Superior temporal | 0.059 | 0.008 | 0.000 | 0.000 | 0.062 | 0.037 | 0.095 | 0.150 | -0.019 | 0.045 | 0.669 | 0.976 | 0.053 | 0.013 | 0.000 | 0.000 |
| Right | Supramarginal | 0.028 | 0.008 | 0.001 | 0.001 | 0.017 | 0.037 | 0.652 | 0.676 | -0.101 | 0.045 | 0.023 | 0.317 | -0.010 | 0.038 | 0.788 | 0.859 |
| Right | Frontal pole | 0.026 | 0.008 | 0.002 | 0.004 | 0.091 | 0.039 | 0.019 | 0.044 | -0.017 | 0.046 | 0.717 | 0.976 | 0.032 | 0.020 | 0.106 | 0.209 |
| Right | Temporal pole | 0.013 | 0.008 | 0.125 | 0.169 | -0.091 | 0.036 | 0.011 | 0.033 | -0.137 | 0.044 | 0.002 | 0.157 | -0.064 | 0.046 | 0.167 | 0.279 |
| Right | Transverse temporal | 0.029 | 0.008 | 0.001 | 0.001 | 0.025 | 0.036 | 0.499 | 0.559 | -0.057 | 0.045 | 0.210 | 0.776 | 0.016 | 0.019 | 0.408 | 0.559 |
| Right | Insula | 0.009 | 0.008 | 0.277 | 0.332 | 0.108 | 0.037 | 0.003 | 0.012 | -0.032 | 0.045 | 0.474 | 0.877 | 0.029 | 0.038 | 0.450 | 0.594 |
|  | Brainstem | -0.002 | 0.008 | 0.818 | 0.839 | 0.016 | 0.036 | 0.651 | 0.676 | 0.002 | 0.045 | 0.967 | 0.990 | -0.001 | 0.008 | 0.910 | 0.939 |

**Supplementary Table 9**. Standardised associations between g and local efficiency in all cohorts, including the meta-analysis, for 85 FreeSurfer regions in FA networks. SE and p-values were estimated by SEM, and q-values were adjusted by FDR.

|  |  | **UKB** | | | | **STRADL** | | | | **LBC1936** | | | | **Meta-analysis** | | | |
| --- | --- | --- | --- | --- | --- | --- | --- | --- | --- | --- | --- | --- | --- | --- | --- | --- | --- |
| **Hemisphere** | **Node** | **β** | ***SE*** | ***p*** | ***q*** | **β** | ***SE*** | ***p*** | ***q*** | **β** | ***SE*** | ***p*** | ***q*** | **β** | ***SE*** | ***p*** | ***q*** |
| Left | Thalamus | 0.039 | 0.008 | 0.000 | 0.000 | 0.269 | 0.055 | 0.000 | 0.000 | 0.083 | 0.045 | 0.066 | 0.137 | 0.123 | 0.068 | 0.072 | 0.078 |
| Left | Caudate | 0.048 | 0.009 | 0.000 | 0.000 | 0.237 | 0.050 | 0.000 | 0.000 | 0.097 | 0.045 | 0.030 | 0.090 | 0.119 | 0.055 | 0.031 | 0.045 |
| Left | Putamen | 0.057 | 0.008 | 0.000 | 0.000 | 0.297 | 0.054 | 0.000 | 0.000 | 0.099 | 0.045 | 0.027 | 0.090 | 0.144 | 0.072 | 0.046 | 0.055 |
| Left | Pallidum | 0.044 | 0.008 | 0.000 | 0.000 | 0.206 | 0.054 | 0.000 | 0.000 | 0.067 | 0.045 | 0.137 | 0.207 | 0.095 | 0.047 | 0.042 | 0.053 |
| Left | Hippocampus | 0.057 | 0.008 | 0.000 | 0.000 | 0.187 | 0.044 | 0.000 | 0.000 | 0.074 | 0.045 | 0.104 | 0.181 | 0.099 | 0.039 | 0.012 | 0.029 |
| Left | Amygdala | 0.037 | 0.008 | 0.000 | 0.000 | 0.124 | 0.044 | 0.005 | 0.006 | 0.080 | 0.046 | 0.078 | 0.151 | 0.066 | 0.027 | 0.015 | 0.031 |
| Left | Accumbens-area | 0.039 | 0.008 | 0.000 | 0.000 | 0.109 | 0.043 | 0.011 | 0.013 | 0.001 | 0.045 | 0.987 | 0.987 | 0.043 | 0.014 | 0.003 | 0.010 |
| Left | Ventral diencephalon | 0.061 | 0.008 | 0.000 | 0.000 | 0.249 | 0.059 | 0.000 | 0.000 | 0.070 | 0.045 | 0.120 | 0.200 | 0.116 | 0.056 | 0.039 | 0.051 |
| Left | Banks of superior temporal | 0.061 | 0.008 | 0.000 | 0.000 | 0.216 | 0.049 | 0.000 | 0.000 | 0.083 | 0.045 | 0.066 | 0.137 | 0.112 | 0.046 | 0.015 | 0.031 |
| Left | Caudal anterior cingulate | 0.043 | 0.008 | 0.000 | 0.000 | 0.090 | 0.041 | 0.027 | 0.030 | -0.007 | 0.045 | 0.885 | 0.906 | 0.043 | 0.008 | 0.000 | 0.000 |
| Left | Caudal middle frontal | 0.059 | 0.009 | 0.000 | 0.000 | 0.219 | 0.046 | 0.000 | 0.000 | 0.068 | 0.045 | 0.128 | 0.204 | 0.110 | 0.050 | 0.028 | 0.043 |
| Left | Cuneus | 0.058 | 0.008 | 0.000 | 0.000 | 0.157 | 0.038 | 0.000 | 0.000 | 0.042 | 0.046 | 0.366 | 0.414 | 0.082 | 0.033 | 0.013 | 0.029 |
| Left | Entorhinal | 0.048 | 0.008 | 0.000 | 0.000 | 0.120 | 0.039 | 0.002 | 0.002 | 0.113 | 0.044 | 0.011 | 0.062 | 0.081 | 0.027 | 0.003 | 0.010 |
| Left | Fusiform | 0.079 | 0.008 | 0.000 | 0.000 | 0.212 | 0.046 | 0.000 | 0.000 | 0.086 | 0.046 | 0.059 | 0.128 | 0.119 | 0.041 | 0.003 | 0.011 |
| Left | Inferior parietal | 0.065 | 0.009 | 0.000 | 0.000 | 0.221 | 0.050 | 0.000 | 0.000 | 0.118 | 0.045 | 0.009 | 0.056 | 0.125 | 0.045 | 0.006 | 0.017 |
| Left | Inferior temporal | 0.044 | 0.008 | 0.000 | 0.000 | 0.216 | 0.050 | 0.000 | 0.000 | 0.074 | 0.045 | 0.103 | 0.181 | 0.103 | 0.051 | 0.043 | 0.053 |
| Left | Isthmus cingulate | 0.038 | 0.008 | 0.000 | 0.000 | 0.247 | 0.050 | 0.000 | 0.000 | 0.087 | 0.045 | 0.053 | 0.128 | 0.117 | 0.062 | 0.059 | 0.066 |
| Left | Lateral occipital | 0.071 | 0.009 | 0.000 | 0.000 | 0.223 | 0.050 | 0.000 | 0.000 | 0.137 | 0.045 | 0.003 | 0.056 | 0.133 | 0.045 | 0.003 | 0.010 |
| Left | Lateral orbitofrontal | 0.067 | 0.009 | 0.000 | 0.000 | 0.132 | 0.041 | 0.001 | 0.002 | 0.074 | 0.045 | 0.097 | 0.175 | 0.076 | 0.016 | 0.000 | 0.000 |
| Left | Lingual | 0.070 | 0.008 | 0.000 | 0.000 | 0.183 | 0.042 | 0.000 | 0.000 | 0.063 | 0.046 | 0.172 | 0.244 | 0.100 | 0.036 | 0.005 | 0.016 |
| Left | Medial orbitofrontal | 0.061 | 0.009 | 0.000 | 0.000 | 0.095 | 0.045 | 0.035 | 0.038 | 0.041 | 0.045 | 0.361 | 0.414 | 0.062 | 0.008 | 0.000 | 0.000 |
| Left | Middle temporal | 0.046 | 0.008 | 0.000 | 0.000 | 0.210 | 0.051 | 0.000 | 0.000 | 0.092 | 0.045 | 0.041 | 0.115 | 0.106 | 0.048 | 0.026 | 0.041 |
| Left | Parahippocampal | 0.071 | 0.008 | 0.000 | 0.000 | 0.210 | 0.040 | 0.000 | 0.000 | 0.054 | 0.045 | 0.232 | 0.295 | 0.109 | 0.047 | 0.021 | 0.036 |
| Left | Paracentral | 0.031 | 0.008 | 0.000 | 0.000 | 0.200 | 0.047 | 0.000 | 0.000 | 0.127 | 0.044 | 0.004 | 0.056 | 0.111 | 0.051 | 0.030 | 0.044 |
| Left | Pars opercularis | 0.056 | 0.009 | 0.000 | 0.000 | 0.198 | 0.048 | 0.000 | 0.000 | 0.081 | 0.045 | 0.071 | 0.144 | 0.103 | 0.042 | 0.014 | 0.031 |
| Left | Pars orbitalis | 0.065 | 0.009 | 0.000 | 0.000 | 0.187 | 0.042 | 0.000 | 0.000 | 0.076 | 0.045 | 0.089 | 0.164 | 0.103 | 0.037 | 0.006 | 0.017 |
| Left | Pars triangularis | 0.064 | 0.009 | 0.000 | 0.000 | 0.226 | 0.045 | 0.000 | 0.000 | 0.057 | 0.045 | 0.200 | 0.266 | 0.111 | 0.053 | 0.034 | 0.047 |
| Left | Pericalcarine | 0.068 | 0.008 | 0.000 | 0.000 | 0.175 | 0.041 | 0.000 | 0.000 | 0.087 | 0.045 | 0.056 | 0.128 | 0.102 | 0.033 | 0.002 | 0.008 |
| Left | Postcentral | 0.048 | 0.008 | 0.000 | 0.000 | 0.234 | 0.049 | 0.000 | 0.000 | 0.108 | 0.045 | 0.015 | 0.076 | 0.122 | 0.055 | 0.025 | 0.040 |
| Left | Posterior cingulate | 0.038 | 0.008 | 0.000 | 0.000 | 0.171 | 0.045 | 0.000 | 0.000 | 0.086 | 0.045 | 0.054 | 0.128 | 0.089 | 0.039 | 0.024 | 0.040 |
| Left | Precentral | 0.049 | 0.008 | 0.000 | 0.000 | 0.227 | 0.050 | 0.000 | 0.000 | 0.121 | 0.045 | 0.007 | 0.056 | 0.124 | 0.052 | 0.018 | 0.035 |
| Left | Precuneus | 0.043 | 0.008 | 0.000 | 0.000 | 0.223 | 0.051 | 0.000 | 0.000 | 0.104 | 0.045 | 0.020 | 0.081 | 0.115 | 0.053 | 0.029 | 0.044 |
| Left | Rostral anterior cingulate | 0.049 | 0.008 | 0.000 | 0.000 | 0.080 | 0.039 | 0.039 | 0.042 | -0.050 | 0.045 | 0.263 | 0.319 | 0.033 | 0.032 | 0.303 | 0.311 |
| Left | Rostral middle frontal | 0.052 | 0.009 | 0.000 | 0.000 | 0.200 | 0.044 | 0.000 | 0.000 | 0.105 | 0.044 | 0.018 | 0.081 | 0.111 | 0.044 | 0.012 | 0.029 |
| Left | Superior frontal | 0.063 | 0.009 | 0.000 | 0.000 | 0.228 | 0.047 | 0.000 | 0.000 | 0.097 | 0.045 | 0.029 | 0.090 | 0.122 | 0.049 | 0.013 | 0.029 |
| Left | Superior parietal | 0.051 | 0.008 | 0.000 | 0.000 | 0.234 | 0.051 | 0.000 | 0.000 | 0.118 | 0.045 | 0.009 | 0.056 | 0.126 | 0.053 | 0.019 | 0.035 |
| Left | Superior temporal | 0.049 | 0.008 | 0.000 | 0.000 | 0.243 | 0.052 | 0.000 | 0.000 | 0.056 | 0.045 | 0.216 | 0.278 | 0.110 | 0.061 | 0.071 | 0.077 |
| Left | Supramarginal | 0.058 | 0.008 | 0.000 | 0.000 | 0.222 | 0.050 | 0.000 | 0.000 | 0.051 | 0.045 | 0.257 | 0.317 | 0.104 | 0.052 | 0.047 | 0.055 |
| Left | Frontal pole | 0.043 | 0.009 | 0.000 | 0.000 | 0.157 | 0.041 | 0.000 | 0.000 | 0.059 | 0.045 | 0.190 | 0.260 | 0.079 | 0.035 | 0.022 | 0.037 |
| Left | Temporal pole | 0.030 | 0.008 | 0.000 | 0.000 | 0.068 | 0.039 | 0.079 | 0.084 | 0.069 | 0.045 | 0.127 | 0.204 | 0.036 | 0.012 | 0.002 | 0.010 |
| Left | Transverse temporal | 0.038 | 0.008 | 0.000 | 0.000 | 0.131 | 0.045 | 0.004 | 0.004 | -0.013 | 0.045 | 0.778 | 0.827 | 0.049 | 0.035 | 0.153 | 0.159 |
| Left | Insula | 0.055 | 0.008 | 0.000 | 0.000 | 0.178 | 0.045 | 0.000 | 0.000 | 0.035 | 0.045 | 0.445 | 0.491 | 0.084 | 0.040 | 0.037 | 0.050 |
| Right | Thalamus | 0.049 | 0.008 | 0.000 | 0.000 | 0.261 | 0.054 | 0.000 | 0.000 | 0.047 | 0.045 | 0.298 | 0.357 | 0.113 | 0.068 | 0.096 | 0.101 |
| Right | Caudate | 0.049 | 0.008 | 0.000 | 0.000 | 0.135 | 0.045 | 0.002 | 0.003 | 0.068 | 0.045 | 0.130 | 0.204 | 0.070 | 0.024 | 0.004 | 0.013 |
| Right | Putamen | 0.053 | 0.008 | 0.000 | 0.000 | 0.238 | 0.051 | 0.000 | 0.000 | 0.086 | 0.045 | 0.056 | 0.128 | 0.118 | 0.055 | 0.032 | 0.045 |
| Right | Pallidum | 0.057 | 0.008 | 0.000 | 0.000 | 0.143 | 0.050 | 0.004 | 0.005 | 0.024 | 0.045 | 0.598 | 0.652 | 0.059 | 0.008 | 0.000 | 0.000 |
| Right | Hippocampus | 0.051 | 0.008 | 0.000 | 0.000 | 0.132 | 0.042 | 0.002 | 0.002 | 0.080 | 0.045 | 0.075 | 0.149 | 0.075 | 0.025 | 0.003 | 0.010 |
| Right | Amygdala | 0.032 | 0.009 | 0.000 | 0.000 | 0.019 | 0.039 | 0.625 | 0.625 | -0.011 | 0.045 | 0.802 | 0.841 | 0.030 | 0.008 | 0.000 | 0.003 |
| Right | Accumbens-area | 0.025 | 0.008 | 0.002 | 0.003 | 0.047 | 0.038 | 0.225 | 0.230 | 0.016 | 0.045 | 0.721 | 0.776 | 0.026 | 0.008 | 0.001 | 0.007 |
| Right | Ventral diencephalon | 0.043 | 0.008 | 0.000 | 0.000 | 0.190 | 0.058 | 0.001 | 0.001 | 0.046 | 0.045 | 0.305 | 0.360 | 0.080 | 0.042 | 0.056 | 0.063 |
| Right | Banks of superior temporal | 0.040 | 0.008 | 0.000 | 0.000 | 0.202 | 0.049 | 0.000 | 0.000 | 0.062 | 0.045 | 0.168 | 0.242 | 0.094 | 0.049 | 0.054 | 0.063 |
| Right | Caudal anterior cingulate | 0.044 | 0.008 | 0.000 | 0.000 | 0.139 | 0.042 | 0.001 | 0.001 | -0.004 | 0.045 | 0.934 | 0.945 | 0.058 | 0.036 | 0.105 | 0.110 |
| Right | Caudal middle frontal | 0.078 | 0.009 | 0.000 | 0.000 | 0.186 | 0.041 | 0.000 | 0.000 | 0.099 | 0.045 | 0.028 | 0.090 | 0.113 | 0.033 | 0.001 | 0.004 |
| Right | Cuneus | 0.055 | 0.008 | 0.000 | 0.000 | 0.122 | 0.039 | 0.002 | 0.002 | 0.104 | 0.045 | 0.021 | 0.081 | 0.080 | 0.024 | 0.001 | 0.005 |
| Right | Entorhinal | 0.062 | 0.009 | 0.000 | 0.000 | 0.082 | 0.037 | 0.028 | 0.031 | 0.079 | 0.045 | 0.082 | 0.154 | 0.063 | 0.008 | 0.000 | 0.000 |
| Right | Fusiform | 0.073 | 0.008 | 0.000 | 0.000 | 0.156 | 0.041 | 0.000 | 0.000 | 0.102 | 0.045 | 0.024 | 0.090 | 0.098 | 0.025 | 0.000 | 0.001 |
| Right | Inferior parietal | 0.061 | 0.008 | 0.000 | 0.000 | 0.213 | 0.049 | 0.000 | 0.000 | 0.090 | 0.045 | 0.045 | 0.115 | 0.113 | 0.045 | 0.011 | 0.029 |
| Right | Inferior temporal | 0.048 | 0.008 | 0.000 | 0.000 | 0.213 | 0.048 | 0.000 | 0.000 | 0.067 | 0.045 | 0.136 | 0.207 | 0.102 | 0.050 | 0.041 | 0.052 |
| Right | Isthmus cingulate | 0.043 | 0.008 | 0.000 | 0.000 | 0.218 | 0.049 | 0.000 | 0.000 | 0.098 | 0.045 | 0.028 | 0.090 | 0.111 | 0.051 | 0.030 | 0.044 |
| Right | Lateral occipital | 0.071 | 0.008 | 0.000 | 0.000 | 0.182 | 0.048 | 0.000 | 0.000 | 0.120 | 0.045 | 0.007 | 0.056 | 0.111 | 0.033 | 0.001 | 0.005 |
| Right | Lateral orbitofrontal | 0.059 | 0.008 | 0.000 | 0.000 | 0.140 | 0.040 | 0.000 | 0.001 | 0.052 | 0.045 | 0.248 | 0.310 | 0.076 | 0.024 | 0.001 | 0.007 |
| Right | Lingual | 0.061 | 0.008 | 0.000 | 0.000 | 0.228 | 0.043 | 0.000 | 0.000 | 0.058 | 0.045 | 0.196 | 0.265 | 0.112 | 0.055 | 0.039 | 0.051 |
| Right | Medial orbitofrontal | 0.050 | 0.009 | 0.000 | 0.000 | 0.064 | 0.040 | 0.106 | 0.111 | -0.009 | 0.045 | 0.836 | 0.867 | 0.049 | 0.009 | 0.000 | 0.000 |
| Right | Middle temporal | 0.040 | 0.008 | 0.000 | 0.000 | 0.231 | 0.049 | 0.000 | 0.000 | 0.064 | 0.045 | 0.155 | 0.231 | 0.106 | 0.058 | 0.070 | 0.077 |
| Right | Parahippocampal | 0.068 | 0.008 | 0.000 | 0.000 | 0.173 | 0.041 | 0.000 | 0.000 | 0.125 | 0.045 | 0.005 | 0.056 | 0.112 | 0.034 | 0.001 | 0.005 |
| Right | Paracentral | 0.044 | 0.008 | 0.000 | 0.000 | 0.252 | 0.047 | 0.000 | 0.000 | 0.150 | 0.044 | 0.001 | 0.056 | 0.142 | 0.061 | 0.021 | 0.036 |
| Right | Pars opercularis | 0.073 | 0.009 | 0.000 | 0.000 | 0.176 | 0.043 | 0.000 | 0.000 | 0.091 | 0.045 | 0.043 | 0.115 | 0.103 | 0.030 | 0.001 | 0.005 |
| Right | Pars orbitalis | 0.079 | 0.009 | 0.000 | 0.000 | 0.196 | 0.041 | 0.000 | 0.000 | 0.120 | 0.044 | 0.007 | 0.056 | 0.124 | 0.036 | 0.001 | 0.004 |
| Right | Pars triangularis | 0.075 | 0.009 | 0.000 | 0.000 | 0.204 | 0.044 | 0.000 | 0.000 | 0.091 | 0.045 | 0.042 | 0.115 | 0.116 | 0.039 | 0.003 | 0.010 |
| Right | Pericalcarine | 0.073 | 0.008 | 0.000 | 0.000 | 0.145 | 0.043 | 0.001 | 0.001 | 0.120 | 0.045 | 0.007 | 0.056 | 0.098 | 0.024 | 0.000 | 0.001 |
| Right | Postcentral | 0.058 | 0.008 | 0.000 | 0.000 | 0.242 | 0.047 | 0.000 | 0.000 | 0.112 | 0.045 | 0.013 | 0.068 | 0.131 | 0.055 | 0.017 | 0.033 |
| Right | Posterior cingulate | 0.045 | 0.008 | 0.000 | 0.000 | 0.171 | 0.044 | 0.000 | 0.000 | 0.045 | 0.045 | 0.318 | 0.371 | 0.081 | 0.039 | 0.039 | 0.051 |
| Right | Precentral | 0.048 | 0.008 | 0.000 | 0.000 | 0.206 | 0.048 | 0.000 | 0.000 | 0.121 | 0.045 | 0.006 | 0.056 | 0.116 | 0.047 | 0.014 | 0.029 |
| Right | Precuneus | 0.049 | 0.008 | 0.000 | 0.000 | 0.230 | 0.050 | 0.000 | 0.000 | 0.118 | 0.045 | 0.008 | 0.056 | 0.124 | 0.053 | 0.019 | 0.035 |
| Right | Rostral anterior cingulate | 0.045 | 0.008 | 0.000 | 0.000 | 0.098 | 0.038 | 0.009 | 0.011 | -0.124 | 0.044 | 0.005 | 0.056 | 0.010 | 0.064 | 0.882 | 0.882 |
| Right | Rostral middle frontal | 0.059 | 0.009 | 0.000 | 0.000 | 0.191 | 0.042 | 0.000 | 0.000 | 0.103 | 0.044 | 0.020 | 0.081 | 0.110 | 0.040 | 0.006 | 0.016 |
| Right | Superior frontal | 0.063 | 0.009 | 0.000 | 0.000 | 0.215 | 0.046 | 0.000 | 0.000 | 0.107 | 0.045 | 0.017 | 0.080 | 0.120 | 0.045 | 0.008 | 0.020 |
| Right | Superior parietal | 0.059 | 0.008 | 0.000 | 0.000 | 0.263 | 0.052 | 0.000 | 0.000 | 0.118 | 0.045 | 0.009 | 0.056 | 0.138 | 0.060 | 0.020 | 0.036 |
| Right | Superior temporal | 0.040 | 0.008 | 0.000 | 0.000 | 0.197 | 0.048 | 0.000 | 0.000 | 0.070 | 0.045 | 0.120 | 0.200 | 0.094 | 0.046 | 0.043 | 0.053 |
| Right | Supramarginal | 0.052 | 0.008 | 0.000 | 0.000 | 0.228 | 0.048 | 0.000 | 0.000 | 0.056 | 0.045 | 0.210 | 0.275 | 0.107 | 0.055 | 0.054 | 0.063 |
| Right | Frontal pole | 0.041 | 0.009 | 0.000 | 0.000 | 0.105 | 0.040 | 0.009 | 0.011 | 0.061 | 0.045 | 0.176 | 0.246 | 0.054 | 0.018 | 0.003 | 0.010 |
| Right | Temporal pole | 0.020 | 0.008 | 0.016 | 0.016 | 0.020 | 0.038 | 0.606 | 0.613 | 0.063 | 0.045 | 0.162 | 0.237 | 0.022 | 0.008 | 0.007 | 0.019 |
| Right | Transverse temporal | 0.001 | 0.008 | 0.890 | 0.890 | 0.052 | 0.037 | 0.160 | 0.166 | -0.034 | 0.045 | 0.444 | 0.491 | 0.002 | 0.008 | 0.765 | 0.774 |
| Right | Insula | 0.034 | 0.008 | 0.000 | 0.000 | 0.182 | 0.048 | 0.000 | 0.000 | 0.090 | 0.045 | 0.045 | 0.115 | 0.093 | 0.044 | 0.034 | 0.047 |
|  | Brainstem | 0.037 | 0.008 | 0.000 | 0.000 | 0.149 | 0.047 | 0.001 | 0.002 | 0.085 | 0.045 | 0.058 | 0.128 | 0.078 | 0.034 | 0.020 | 0.036 |

**Supplementary Table 10**. Standardised associations between g and local efficiency in all cohorts, including the meta-analysis, for 85 FreeSurfer regions in MD networks. SE and p-values were estimated by SEM, and q-values were adjusted by FDR.

|  |  | **UKB** | | | | **STRADL** | | | | **LBC1936** | | | | **Meta-analysis** | | | |
| --- | --- | --- | --- | --- | --- | --- | --- | --- | --- | --- | --- | --- | --- | --- | --- | --- | --- |
| **Hemisphere** | **Node** | **β** | ***SE*** | ***p*** | ***q*** | **β** | ***SE*** | ***p*** | ***q*** | **β** | ***SE*** | ***p*** | ***q*** | **β** | ***SE*** | ***p*** | ***q*** |
| Left | Thalamus | -0.076 | 0.010 | 0.000 | 0.000 | -0.238 | 0.055 | 0.000 | 0.000 | -0.141 | 0.045 | 0.002 | 0.076 | -0.140 | 0.047 | 0.003 | 0.011 |
| Left | Caudate | -0.085 | 0.010 | 0.000 | 0.000 | -0.199 | 0.050 | 0.000 | 0.000 | -0.116 | 0.045 | 0.010 | 0.078 | -0.120 | 0.032 | 0.000 | 0.001 |
| Left | Putamen | -0.068 | 0.009 | 0.000 | 0.000 | -0.229 | 0.055 | 0.000 | 0.000 | -0.102 | 0.045 | 0.025 | 0.091 | -0.121 | 0.046 | 0.009 | 0.019 |
| Left | Pallidum | -0.061 | 0.009 | 0.000 | 0.000 | -0.224 | 0.058 | 0.000 | 0.000 | -0.060 | 0.046 | 0.194 | 0.235 | -0.104 | 0.049 | 0.032 | 0.043 |
| Left | Hippocampus | -0.040 | 0.009 | 0.000 | 0.000 | -0.119 | 0.046 | 0.009 | 0.011 | -0.046 | 0.046 | 0.319 | 0.348 | -0.053 | 0.020 | 0.009 | 0.019 |
| Left | Amygdala | -0.035 | 0.009 | 0.000 | 0.000 | -0.122 | 0.046 | 0.008 | 0.011 | -0.068 | 0.045 | 0.130 | 0.181 | -0.060 | 0.025 | 0.018 | 0.029 |
| Left | Accumbens-area | -0.021 | 0.008 | 0.013 | 0.014 | -0.080 | 0.040 | 0.048 | 0.052 | 0.002 | 0.045 | 0.971 | 0.971 | -0.023 | 0.008 | 0.006 | 0.017 |
| Left | Ventral diencephalon | -0.045 | 0.009 | 0.000 | 0.000 | -0.174 | 0.053 | 0.001 | 0.002 | -0.104 | 0.046 | 0.022 | 0.091 | -0.094 | 0.038 | 0.013 | 0.023 |
| Left | Banks of superior temporal | -0.048 | 0.009 | 0.000 | 0.000 | -0.199 | 0.055 | 0.000 | 0.001 | -0.066 | 0.046 | 0.149 | 0.197 | -0.092 | 0.043 | 0.032 | 0.043 |
| Left | Caudal anterior cingulate | -0.057 | 0.009 | 0.000 | 0.000 | -0.156 | 0.052 | 0.003 | 0.004 | 0.022 | 0.045 | 0.631 | 0.647 | -0.061 | 0.044 | 0.173 | 0.177 |
| Left | Caudal middle frontal | -0.076 | 0.009 | 0.000 | 0.000 | -0.211 | 0.050 | 0.000 | 0.000 | -0.077 | 0.045 | 0.088 | 0.145 | -0.113 | 0.041 | 0.006 | 0.017 |
| Left | Cuneus | -0.014 | 0.009 | 0.117 | 0.121 | -0.147 | 0.060 | 0.014 | 0.017 | -0.107 | 0.045 | 0.017 | 0.091 | -0.074 | 0.042 | 0.081 | 0.089 |
| Left | Entorhinal | -0.036 | 0.008 | 0.000 | 0.000 | -0.042 | 0.044 | 0.332 | 0.345 | -0.094 | 0.045 | 0.036 | 0.098 | -0.038 | 0.008 | 0.000 | 0.000 |
| Left | Fusiform | -0.047 | 0.009 | 0.000 | 0.000 | -0.194 | 0.054 | 0.000 | 0.001 | -0.074 | 0.046 | 0.103 | 0.154 | -0.093 | 0.042 | 0.026 | 0.036 |
| Left | Inferior parietal | -0.052 | 0.009 | 0.000 | 0.000 | -0.198 | 0.058 | 0.001 | 0.001 | -0.116 | 0.045 | 0.010 | 0.078 | -0.107 | 0.042 | 0.010 | 0.021 |
| Left | Inferior temporal | -0.048 | 0.009 | 0.000 | 0.000 | -0.195 | 0.055 | 0.000 | 0.001 | -0.100 | 0.046 | 0.028 | 0.091 | -0.101 | 0.042 | 0.016 | 0.026 |
| Left | Isthmus cingulate | -0.048 | 0.009 | 0.000 | 0.000 | -0.233 | 0.059 | 0.000 | 0.000 | -0.119 | 0.046 | 0.009 | 0.078 | -0.121 | 0.053 | 0.022 | 0.032 |
| Left | Lateral occipital | -0.031 | 0.009 | 0.001 | 0.001 | -0.191 | 0.059 | 0.001 | 0.002 | -0.125 | 0.045 | 0.006 | 0.076 | -0.102 | 0.048 | 0.034 | 0.043 |
| Left | Lateral orbitofrontal | -0.050 | 0.009 | 0.000 | 0.000 | -0.144 | 0.046 | 0.002 | 0.002 | -0.100 | 0.045 | 0.026 | 0.091 | -0.084 | 0.029 | 0.004 | 0.014 |
| Left | Lingual | -0.018 | 0.009 | 0.049 | 0.052 | -0.192 | 0.057 | 0.001 | 0.001 | -0.077 | 0.046 | 0.093 | 0.147 | -0.084 | 0.050 | 0.093 | 0.098 |
| Left | Medial orbitofrontal | -0.059 | 0.009 | 0.000 | 0.000 | -0.110 | 0.043 | 0.011 | 0.014 | -0.056 | 0.045 | 0.212 | 0.247 | -0.061 | 0.009 | 0.000 | 0.000 |
| Left | Middle temporal | -0.056 | 0.009 | 0.000 | 0.000 | -0.212 | 0.056 | 0.000 | 0.000 | -0.101 | 0.045 | 0.027 | 0.091 | -0.110 | 0.044 | 0.013 | 0.023 |
| Left | Parahippocampal | -0.006 | 0.009 | 0.478 | 0.483 | -0.187 | 0.050 | 0.000 | 0.000 | -0.042 | 0.046 | 0.364 | 0.392 | -0.071 | 0.054 | 0.189 | 0.191 |
| Left | Paracentral | -0.099 | 0.009 | 0.000 | 0.000 | -0.273 | 0.055 | 0.000 | 0.000 | -0.115 | 0.045 | 0.011 | 0.080 | -0.152 | 0.052 | 0.003 | 0.012 |
| Left | Pars opercularis | -0.071 | 0.009 | 0.000 | 0.000 | -0.223 | 0.052 | 0.000 | 0.000 | -0.087 | 0.045 | 0.054 | 0.111 | -0.118 | 0.045 | 0.009 | 0.019 |
| Left | Pars orbitalis | -0.069 | 0.009 | 0.000 | 0.000 | -0.172 | 0.047 | 0.000 | 0.001 | -0.085 | 0.045 | 0.059 | 0.117 | -0.097 | 0.029 | 0.001 | 0.005 |
| Left | Pars triangularis | -0.078 | 0.009 | 0.000 | 0.000 | -0.226 | 0.051 | 0.000 | 0.000 | -0.067 | 0.045 | 0.136 | 0.187 | -0.116 | 0.047 | 0.013 | 0.023 |
| Left | Pericalcarine | -0.019 | 0.009 | 0.034 | 0.036 | -0.147 | 0.055 | 0.008 | 0.010 | -0.060 | 0.045 | 0.184 | 0.230 | -0.060 | 0.036 | 0.093 | 0.098 |
| Left | Postcentral | -0.080 | 0.009 | 0.000 | 0.000 | -0.228 | 0.054 | 0.000 | 0.000 | -0.092 | 0.045 | 0.044 | 0.106 | -0.122 | 0.043 | 0.004 | 0.015 |
| Left | Posterior cingulate | -0.090 | 0.009 | 0.000 | 0.000 | -0.259 | 0.056 | 0.000 | 0.000 | -0.096 | 0.046 | 0.035 | 0.098 | -0.139 | 0.051 | 0.006 | 0.018 |
| Left | Precentral | -0.081 | 0.009 | 0.000 | 0.000 | -0.249 | 0.053 | 0.000 | 0.000 | -0.085 | 0.045 | 0.062 | 0.117 | -0.130 | 0.051 | 0.011 | 0.022 |
| Left | Precuneus | -0.058 | 0.009 | 0.000 | 0.000 | -0.248 | 0.059 | 0.000 | 0.000 | -0.112 | 0.045 | 0.014 | 0.090 | -0.128 | 0.054 | 0.018 | 0.029 |
| Left | Rostral anterior cingulate | -0.055 | 0.009 | 0.000 | 0.000 | -0.140 | 0.044 | 0.001 | 0.002 | -0.059 | 0.045 | 0.184 | 0.230 | -0.073 | 0.023 | 0.002 | 0.007 |
| Left | Rostral middle frontal | -0.072 | 0.009 | 0.000 | 0.000 | -0.181 | 0.048 | 0.000 | 0.000 | -0.102 | 0.045 | 0.022 | 0.091 | -0.106 | 0.032 | 0.001 | 0.005 |
| Left | Superior frontal | -0.094 | 0.009 | 0.000 | 0.000 | -0.223 | 0.051 | 0.000 | 0.000 | -0.097 | 0.045 | 0.030 | 0.091 | -0.128 | 0.038 | 0.001 | 0.004 |
| Left | Superior parietal | -0.060 | 0.009 | 0.000 | 0.000 | -0.230 | 0.057 | 0.000 | 0.000 | -0.123 | 0.045 | 0.007 | 0.076 | -0.125 | 0.048 | 0.010 | 0.021 |
| Left | Superior temporal | -0.048 | 0.009 | 0.000 | 0.000 | -0.232 | 0.057 | 0.000 | 0.000 | -0.090 | 0.046 | 0.049 | 0.107 | -0.112 | 0.053 | 0.034 | 0.043 |
| Left | Supramarginal | -0.059 | 0.009 | 0.000 | 0.000 | -0.221 | 0.054 | 0.000 | 0.000 | -0.071 | 0.046 | 0.119 | 0.169 | -0.107 | 0.048 | 0.026 | 0.036 |
| Left | Frontal pole | -0.044 | 0.009 | 0.000 | 0.000 | -0.134 | 0.046 | 0.003 | 0.005 | -0.086 | 0.046 | 0.060 | 0.117 | -0.074 | 0.028 | 0.007 | 0.019 |
| Left | Temporal pole | -0.042 | 0.009 | 0.000 | 0.000 | -0.083 | 0.047 | 0.076 | 0.080 | -0.082 | 0.045 | 0.068 | 0.126 | -0.046 | 0.011 | 0.000 | 0.000 |
| Left | Transverse temporal | -0.031 | 0.009 | 0.000 | 0.001 | -0.266 | 0.060 | 0.000 | 0.000 | -0.019 | 0.046 | 0.687 | 0.695 | -0.098 | 0.076 | 0.200 | 0.200 |
| Left | Insula | -0.061 | 0.009 | 0.000 | 0.000 | -0.199 | 0.055 | 0.000 | 0.001 | -0.075 | 0.045 | 0.097 | 0.150 | -0.099 | 0.039 | 0.011 | 0.021 |
| Right | Thalamus | -0.068 | 0.009 | 0.000 | 0.000 | -0.259 | 0.059 | 0.000 | 0.000 | -0.090 | 0.046 | 0.050 | 0.107 | -0.129 | 0.056 | 0.022 | 0.032 |
| Right | Caudate | -0.084 | 0.009 | 0.000 | 0.000 | -0.235 | 0.048 | 0.000 | 0.000 | -0.088 | 0.045 | 0.049 | 0.107 | -0.129 | 0.047 | 0.006 | 0.017 |
| Right | Putamen | -0.069 | 0.009 | 0.000 | 0.000 | -0.228 | 0.056 | 0.000 | 0.000 | -0.081 | 0.046 | 0.074 | 0.130 | -0.116 | 0.047 | 0.013 | 0.023 |
| Right | Pallidum | -0.069 | 0.009 | 0.000 | 0.000 | -0.205 | 0.058 | 0.000 | 0.001 | -0.031 | 0.046 | 0.506 | 0.525 | -0.092 | 0.045 | 0.039 | 0.048 |
| Right | Hippocampus | -0.028 | 0.009 | 0.001 | 0.001 | -0.122 | 0.050 | 0.015 | 0.018 | -0.049 | 0.046 | 0.290 | 0.325 | -0.049 | 0.025 | 0.048 | 0.056 |
| Right | Amygdala | -0.038 | 0.009 | 0.000 | 0.000 | -0.035 | 0.039 | 0.374 | 0.383 | -0.076 | 0.045 | 0.091 | 0.145 | -0.039 | 0.008 | 0.000 | 0.000 |
| Right | Accumbens-area | -0.039 | 0.009 | 0.000 | 0.000 | -0.076 | 0.039 | 0.048 | 0.052 | -0.064 | 0.045 | 0.153 | 0.197 | -0.042 | 0.008 | 0.000 | 0.000 |
| Right | Ventral diencephalon | -0.052 | 0.009 | 0.000 | 0.000 | -0.231 | 0.059 | 0.000 | 0.000 | -0.101 | 0.046 | 0.027 | 0.091 | -0.115 | 0.051 | 0.023 | 0.032 |
| Right | Banks of superior temporal | -0.043 | 0.009 | 0.000 | 0.000 | -0.234 | 0.062 | 0.000 | 0.000 | -0.059 | 0.046 | 0.202 | 0.238 | -0.100 | 0.056 | 0.073 | 0.082 |
| Right | Caudal anterior cingulate | -0.073 | 0.009 | 0.000 | 0.000 | -0.164 | 0.052 | 0.002 | 0.002 | -0.035 | 0.045 | 0.446 | 0.469 | -0.079 | 0.020 | 0.000 | 0.001 |
| Right | Caudal middle frontal | -0.082 | 0.009 | 0.000 | 0.000 | -0.179 | 0.046 | 0.000 | 0.000 | -0.056 | 0.045 | 0.220 | 0.252 | -0.099 | 0.030 | 0.001 | 0.005 |
| Right | Cuneus | -0.028 | 0.009 | 0.001 | 0.001 | -0.075 | 0.054 | 0.165 | 0.173 | -0.120 | 0.044 | 0.007 | 0.076 | -0.061 | 0.030 | 0.044 | 0.052 |
| Right | Entorhinal | -0.032 | 0.008 | 0.000 | 0.000 | -0.016 | 0.039 | 0.687 | 0.696 | -0.099 | 0.045 | 0.026 | 0.091 | -0.034 | 0.008 | 0.000 | 0.000 |
| Right | Fusiform | -0.029 | 0.009 | 0.001 | 0.001 | -0.119 | 0.052 | 0.023 | 0.027 | -0.095 | 0.046 | 0.039 | 0.103 | -0.064 | 0.030 | 0.035 | 0.043 |
| Right | Inferior parietal | -0.058 | 0.009 | 0.000 | 0.000 | -0.204 | 0.059 | 0.001 | 0.001 | -0.109 | 0.046 | 0.017 | 0.091 | -0.108 | 0.041 | 0.008 | 0.019 |
| Right | Inferior temporal | -0.038 | 0.009 | 0.000 | 0.000 | -0.160 | 0.056 | 0.004 | 0.006 | -0.092 | 0.046 | 0.045 | 0.106 | -0.080 | 0.035 | 0.024 | 0.033 |
| Right | Isthmus cingulate | -0.046 | 0.009 | 0.000 | 0.000 | -0.203 | 0.062 | 0.001 | 0.002 | -0.100 | 0.046 | 0.030 | 0.091 | -0.100 | 0.043 | 0.021 | 0.032 |
| Right | Lateral occipital | -0.026 | 0.009 | 0.003 | 0.004 | -0.138 | 0.061 | 0.025 | 0.028 | -0.124 | 0.045 | 0.006 | 0.076 | -0.080 | 0.040 | 0.044 | 0.052 |
| Right | Lateral orbitofrontal | -0.058 | 0.009 | 0.000 | 0.000 | -0.170 | 0.044 | 0.000 | 0.000 | -0.079 | 0.045 | 0.075 | 0.130 | -0.093 | 0.033 | 0.005 | 0.017 |
| Right | Lingual | -0.017 | 0.009 | 0.057 | 0.060 | -0.122 | 0.060 | 0.042 | 0.047 | -0.091 | 0.046 | 0.047 | 0.107 | -0.059 | 0.035 | 0.088 | 0.096 |
| Right | Medial orbitofrontal | -0.051 | 0.009 | 0.000 | 0.000 | -0.110 | 0.040 | 0.007 | 0.009 | -0.058 | 0.045 | 0.197 | 0.236 | -0.057 | 0.013 | 0.000 | 0.000 |
| Right | Middle temporal | -0.041 | 0.009 | 0.000 | 0.000 | -0.193 | 0.058 | 0.001 | 0.001 | -0.078 | 0.046 | 0.091 | 0.145 | -0.090 | 0.042 | 0.034 | 0.043 |
| Right | Parahippocampal | -0.004 | 0.008 | 0.657 | 0.657 | -0.153 | 0.059 | 0.009 | 0.012 | -0.077 | 0.047 | 0.101 | 0.154 | -0.063 | 0.044 | 0.150 | 0.155 |
| Right | Paracentral | -0.101 | 0.009 | 0.000 | 0.000 | -0.257 | 0.054 | 0.000 | 0.000 | -0.110 | 0.045 | 0.015 | 0.091 | -0.146 | 0.046 | 0.002 | 0.006 |
| Right | Pars opercularis | -0.082 | 0.009 | 0.000 | 0.000 | -0.219 | 0.048 | 0.000 | 0.000 | -0.071 | 0.045 | 0.119 | 0.169 | -0.117 | 0.044 | 0.007 | 0.019 |
| Right | Pars orbitalis | -0.066 | 0.009 | 0.000 | 0.000 | -0.183 | 0.046 | 0.000 | 0.000 | -0.058 | 0.045 | 0.194 | 0.235 | -0.095 | 0.036 | 0.009 | 0.019 |
| Right | Pars triangularis | -0.078 | 0.009 | 0.000 | 0.000 | -0.195 | 0.050 | 0.000 | 0.000 | -0.034 | 0.045 | 0.447 | 0.469 | -0.097 | 0.041 | 0.019 | 0.029 |
| Right | Pericalcarine | -0.013 | 0.009 | 0.120 | 0.123 | -0.121 | 0.058 | 0.037 | 0.042 | -0.129 | 0.045 | 0.004 | 0.076 | -0.075 | 0.042 | 0.077 | 0.086 |
| Right | Postcentral | -0.080 | 0.009 | 0.000 | 0.000 | -0.243 | 0.059 | 0.000 | 0.000 | -0.082 | 0.046 | 0.074 | 0.130 | -0.124 | 0.048 | 0.010 | 0.021 |
| Right | Posterior cingulate | -0.075 | 0.009 | 0.000 | 0.000 | -0.191 | 0.054 | 0.000 | 0.001 | -0.080 | 0.046 | 0.079 | 0.134 | -0.101 | 0.031 | 0.001 | 0.005 |
| Right | Precentral | -0.086 | 0.009 | 0.000 | 0.000 | -0.246 | 0.055 | 0.000 | 0.000 | -0.068 | 0.046 | 0.139 | 0.188 | -0.125 | 0.051 | 0.015 | 0.024 |
| Right | Precuneus | -0.063 | 0.009 | 0.000 | 0.000 | -0.257 | 0.060 | 0.000 | 0.000 | -0.129 | 0.045 | 0.004 | 0.076 | -0.137 | 0.055 | 0.012 | 0.023 |
| Right | Rostral anterior cingulate | -0.052 | 0.009 | 0.000 | 0.000 | -0.109 | 0.042 | 0.009 | 0.011 | -0.084 | 0.044 | 0.058 | 0.117 | -0.065 | 0.017 | 0.000 | 0.001 |
| Right | Rostral middle frontal | -0.082 | 0.009 | 0.000 | 0.000 | -0.204 | 0.045 | 0.000 | 0.000 | -0.100 | 0.045 | 0.024 | 0.091 | -0.120 | 0.036 | 0.001 | 0.005 |
| Right | Superior frontal | -0.097 | 0.009 | 0.000 | 0.000 | -0.221 | 0.048 | 0.000 | 0.000 | -0.091 | 0.045 | 0.044 | 0.106 | -0.128 | 0.038 | 0.001 | 0.004 |
| Right | Superior parietal | -0.065 | 0.009 | 0.000 | 0.000 | -0.249 | 0.060 | 0.000 | 0.000 | -0.134 | 0.045 | 0.003 | 0.076 | -0.137 | 0.052 | 0.009 | 0.019 |
| Right | Superior temporal | -0.039 | 0.009 | 0.000 | 0.000 | -0.214 | 0.059 | 0.000 | 0.001 | -0.104 | 0.046 | 0.023 | 0.091 | -0.106 | 0.050 | 0.033 | 0.043 |
| Right | Supramarginal | -0.057 | 0.009 | 0.000 | 0.000 | -0.230 | 0.059 | 0.000 | 0.000 | -0.052 | 0.046 | 0.257 | 0.292 | -0.103 | 0.053 | 0.050 | 0.058 |
| Right | Frontal pole | -0.041 | 0.009 | 0.000 | 0.000 | -0.137 | 0.042 | 0.001 | 0.002 | -0.092 | 0.045 | 0.042 | 0.106 | -0.078 | 0.031 | 0.011 | 0.021 |
| Right | Temporal pole | -0.037 | 0.009 | 0.000 | 0.000 | -0.002 | 0.039 | 0.955 | 0.955 | -0.096 | 0.045 | 0.031 | 0.091 | -0.038 | 0.008 | 0.000 | 0.000 |
| Right | Transverse temporal | -0.020 | 0.009 | 0.019 | 0.021 | -0.137 | 0.045 | 0.002 | 0.003 | -0.047 | 0.045 | 0.297 | 0.328 | -0.058 | 0.035 | 0.092 | 0.098 |
| Right | Insula | -0.055 | 0.009 | 0.000 | 0.000 | -0.177 | 0.056 | 0.001 | 0.002 | -0.073 | 0.046 | 0.110 | 0.161 | -0.085 | 0.033 | 0.009 | 0.019 |
|  | Brainstem | -0.042 | 0.008 | 0.000 | 0.000 | -0.091 | 0.046 | 0.049 | 0.053 | -0.065 | 0.045 | 0.151 | 0.197 | -0.044 | 0.008 | 0.000 | 0.000 |

**Supplementary Table 11**. Summary of standardised β coefficients for associations between g and nodal local efficiency across 85 network nodes, shown for each cohort and the meta-analysis for four network weightings (SC, SD, FA, and MD). Estimates were obtained using SEM, and n_node_ and n_nodeFDR_ indicate the number of significant nodes before and after FDR correction (p < 0.05).

| **Study** | **Weighting** | **Mean** | **Median** | **Min** | **Max** | ***n*_node_** | ***n_node_*_FDR_** |
| --- | --- | --- | --- | --- | --- | --- | --- |
| UKB | SC | 0.090 | 0.088 | 0.018 | 0.187 | 85 | 85 |
| UKB | SD | 0.028 | 0.030 | -0.020 | 0.108 | 54 | 52 |
| UKB | FA | 0.052 | 0.051 | 0.001 | 0.079 | 84 | 84 |
| UKB | MD | -0.054 | -0.055 | -0.101 | -0.004 | 80 | 79 |
| STRADL | SC | 0.146 | 0.145 | -0.026 | 0.274 | 75 | 74 |
| STRADL | SD | 0.071 | 0.076 | -0.091 | 0.186 | 45 | 38 |
| STRADL | FA | 0.179 | 0.196 | 0.019 | 0.297 | 79 | 79 |
| STRADL | MD | -0.179 | -0.194 | -0.273 | -0.002 | 79 | 76 |
| LBC1936 | SC | 0.062 | 0.070 | -0.099 | 0.205 | 31 | 17 |
| LBC1936 | SD | -0.019 | -0.018 | -0.137 | 0.113 | 8 | 0 |
| LBC1936 | FA | 0.072 | 0.080 | -0.124 | 0.150 | 33 | 0 |
| LBC1936 | MD | -0.083 | -0.085 | -0.141 | 0.022 | 39 | 0 |
| Meta-analysis | SC | 0.098 | 0.095 | -0.018 | 0.198 | 72 | 72 |
| Meta-analysis | SD | 0.029 | 0.030 | -0.064 | 0.110 | 34 | 28 |
| Meta-analysis | FA | 0.094 | 0.103 | 0.002 | 0.144 | 72 | 62 |
| Meta-analysis | MD | -0.093 | -0.098 | -0.152 | -0.023 | 73 | 70 |

**Supplementary Table 12**. Correlations (Pearson’s r) between paired node-*g* β coefficients across cerebral hemispheres in all cohorts, including the meta-analysis, and across network weightings (p < 0.001).

| **Study** | **Weighting** | ***r*** | ***p*** |
| --- | --- | --- | --- |
| UKB | SC | 0.880 | 0.000 |
| UKB | SD | 0.683 | 0.000 |
| UKB | FA | 0.694 | 0.000 |
| UKB | MD | 0.937 | 0.000 |
| STRADL | SC | 0.698 | 0.000 |
| STRADL | SD | 0.563 | 0.000 |
| STRADL | FA | 0.822 | 0.000 |
| STRADL | MD | 0.819 | 0.000 |
| LBC1936 | SC | 0.662 | 0.000 |
| LBC1936 | SD | 0.414 | 0.006 |
| LBC1936 | FA | 0.761 | 0.000 |
| LBC1936 | MD | 0.653 | 0.000 |
| Meta-analysis | SC | 0.839 | 0.000 |
| Meta-analysis | SD | 0.663 | 0.000 |
| Meta-analysis | FA | 0.865 | 0.000 |
| Meta-analysis | MD | 0.879 | 0.000 |

**Supplementary Table 13**. List of grey matter regions (Desikan-Killiany atlas) and abbreviations.

| **Region** | **Abbreviation** | **Lobe/Grouping** |
| --- | --- | --- |
| Caudal middle frontal | CaMF | Frontal |
| Frontal pole | FPo | Frontal |
| Lateral orbitofrontal | LOF | Frontal |
| Medial orbitofrontal | MedOr | Frontal |
| Paracentral | PaC | Frontal |
| Pars opercularis | ParOp | Frontal |
| Pars orbitalis | ParOr | Frontal |
| Pars triangularis | ParTr | Frontal |
| Precentral | PrC | Frontal |
| Rostral middle frontal | RosMF | Frontal |
| Superior frontal | SupF | Frontal |
| Caudal anterior cingulate | CaACg | Cingulate |
| Isthmus cingulate | IsCg | Cingulate |
| Posterior cingulate | PosCg | Cingulate |
| Rostral anterior cingulate | RosACg | Cingulate |
| Insula | Ins | - |
| Banks of superior temporal | bSTS | Temporal |
| Entorhinal | Ent | Temporal |
| Fusiform | Fus | Temporal |
| Inferior temporal | InfT | Temporal |
| Middle temporal | MT | Temporal |
| Parahippocampal | PaHip | Temporal |
| Superior temporal | SupT | Temporal |
| Temporal pole | TPo | Temporal |
| Transverse temporal | TrT | Temporal |
| Inferior parietal | InfP | Parietal |
| Postcentral | PosC | Parietal |
| Precuneus | PrCun | Parietal |
| Superior parietal | SupP | Parietal |
| Supramarginal | SuMar | Parietal |
| Cuneus | Cun | Occipital |
| Lateral occipital | LOc | Occipital |
| Lingual | Lin | Occipital |
| Pericalcarine | PerCa | Occipital |
| Nucleus accumbens | NAcc | Subcortical |
| Amygdala | Amg | Subcortical |
| Caudate nucleus | CaN | Subcortical |
| Hippocampus | Hip | Subcortical |
| Pallidum | Pal | Subcortical |
| Putamen | Put | Subcortical |
| Thalamus | Tha | Subcortical |
| Ventral diencephalon | VDC | Subcortical |
| Brainstem | BSt | - |

**Supplementary Table 14**. Summary of standardised β coefficients for associations between g and 818 network edges in all cohorts, including the meta-analysis, for three weightings (SC, FA, and MD). Estimates were obtained using SEM, and n_edge_ and n_edgeFDR_ indicate the number of significant edges before and after FDR correction (p < 0.05).

| **Study** | **Weighting** | **Mean** | **Median** | **Min** | **Max** | ***n*_edge_** | ***n_edge_*_FDR_** |
| --- | --- | --- | --- | --- | --- | --- | --- |
| UKB | SC | 0.029 | 0.028 | -0.050 | 0.100 | 572 | 554 |
| UKB | FA | 0.032 | 0.032 | -0.033 | 0.091 | 643 | 637 |
| UKB | MD | -0.038 | -0.036 | -0.104 | 0.024 | 619 | 609 |
| STRADL | SC | 0.043 | 0.044 | -0.091 | 0.191 | 243 | 109 |
| STRADL | FA | 0.086 | 0.087 | -0.068 | 0.240 | 471 | 409 |
| STRADL | MD | -0.078 | -0.076 | -0.278 | 0.124 | 389 | 320 |
| LBC1936 | SC | 0.018 | 0.018 | -0.152 | 0.229 | 109 | 16 |
| LBC1936 | FA | 0.039 | 0.039 | -0.146 | 0.194 | 148 | 4 |
| LBC1936 | MD | -0.021 | -0.023 | -0.219 | 0.162 | 119 | 7 |
| Meta-analysis | SC | 0.030 | 0.029 | -0.071 | 0.114 | 434 | 391 |
| Meta-analysis | FA | 0.046 | 0.046 | -0.034 | 0.134 | 503 | 433 |
| Meta-analysis | MD | -0.043 | -0.039 | -0.146 | 0.053 | 383 | 325 |

**Supplementary Table 15**. Summary of β coefficients by inter- and intra-hemisphere connectivity for significant edge-g associations (p < 0.05, FDR), derived from the separate meta-analyses of SC, FA, and MD networks. Modules include: whole-brain (all connections), inter-hemisphere (bilateral), intra-hemisphere (ipsilateral), and intra-hemisphere (left or right only) connections.

| **Study** | **Weighting** | **Connectivity** | ***n*_edge_** | **Mean** | **Median** | **Min** | **Max** |
| --- | --- | --- | --- | --- | --- | --- | --- |
| UKB | SC | Whole-brain | 554 | 0.040 | 0.039 | -0.050 | 0.100 |
| UKB | SC | Inter-hemisphere | 104 | 0.037 | 0.039 | -0.050 | 0.087 |
| UKB | SC | Intra-hemisphere | 447 | 0.041 | 0.040 | -0.049 | 0.100 |
| UKB | SC | Intra-hemisphere (left) | 226 | 0.040 | 0.037 | -0.049 | 0.100 |
| UKB | SC | Intra-hemisphere (right) | 221 | 0.043 | 0.042 | -0.049 | 0.097 |
| UKB | FA | Whole-brain | 637 | 0.040 | 0.038 | -0.033 | 0.091 |
| UKB | FA | Inter-hemisphere | 101 | 0.030 | 0.028 | -0.018 | 0.063 |
| UKB | FA | Intra-hemisphere | 532 | 0.042 | 0.042 | -0.033 | 0.091 |
| UKB | FA | Intra-hemisphere (left) | 271 | 0.040 | 0.040 | -0.027 | 0.084 |
| UKB | FA | Intra-hemisphere (right) | 261 | 0.043 | 0.043 | -0.033 | 0.091 |
| UKB | MD | Whole-brain | 609 | -0.049 | -0.047 | -0.104 | 0.024 |
| UKB | MD | Inter-hemisphere | 125 | -0.048 | -0.043 | -0.104 | 0.018 |
| UKB | MD | Intra-hemisphere | 480 | -0.050 | -0.048 | -0.104 | 0.024 |
| UKB | MD | Intra-hemisphere (left) | 251 | -0.048 | -0.047 | -0.095 | 0.023 |
| UKB | MD | Intra-hemisphere (right) | 229 | -0.052 | -0.049 | -0.104 | 0.024 |
| STRADL | SC | Whole-brain | 109 | 0.119 | 0.114 | 0.097 | 0.191 |
| STRADL | SC | Inter-hemisphere | 29 | 0.117 | 0.110 | 0.097 | 0.191 |
| STRADL | SC | Intra-hemisphere | 79 | 0.120 | 0.116 | 0.098 | 0.179 |
| STRADL | SC | Intra-hemisphere (left) | 37 | 0.120 | 0.116 | 0.098 | 0.168 |
| STRADL | SC | Intra-hemisphere (right) | 42 | 0.120 | 0.116 | 0.099 | 0.179 |
| STRADL | FA | Whole-brain | 409 | 0.128 | 0.121 | 0.082 | 0.240 |
| STRADL | FA | Inter-hemisphere | 106 | 0.127 | 0.123 | 0.085 | 0.217 |
| STRADL | FA | Intra-hemisphere | 301 | 0.128 | 0.121 | 0.082 | 0.240 |
| STRADL | FA | Intra-hemisphere (left) | 161 | 0.133 | 0.125 | 0.082 | 0.240 |
| STRADL | FA | Intra-hemisphere (right) | 140 | 0.122 | 0.116 | 0.083 | 0.208 |
| STRADL | MD | Whole-brain | 320 | -0.153 | -0.156 | -0.278 | 0.124 |
| STRADL | MD | Inter-hemisphere | 47 | -0.137 | -0.162 | -0.223 | 0.124 |
| STRADL | MD | Intra-hemisphere | 273 | -0.156 | -0.156 | -0.278 | 0.120 |
| STRADL | MD | Intra-hemisphere (left) | 145 | -0.158 | -0.154 | -0.243 | -0.095 |
| STRADL | MD | Intra-hemisphere (right) | 128 | -0.155 | -0.156 | -0.278 | 0.120 |
| LBC1936 | SC | Whole-brain | 16 | 0.090 | 0.157 | -0.152 | 0.229 |
| LBC1936 | SC | Inter-hemisphere | 1 | 0.152 | 0.152 | 0.152 | 0.152 |
| LBC1936 | SC | Intra-hemisphere | 14 | 0.081 | 0.157 | -0.152 | 0.229 |
| LBC1936 | SC | Intra-hemisphere (left) | 6 | 0.054 | 0.151 | -0.152 | 0.168 |
| LBC1936 | SC | Intra-hemisphere (right) | 8 | 0.101 | 0.166 | -0.152 | 0.229 |
| LBC1936 | FA | Whole-brain | 4 | 0.182 | 0.180 | 0.172 | 0.194 |
| LBC1936 | FA | Inter-hemisphere | 1 | 0.172 | 0.172 | 0.172 | 0.172 |
| LBC1936 | FA | Intra-hemisphere | 3 | 0.185 | 0.184 | 0.176 | 0.194 |
| LBC1936 | FA | Intra-hemisphere (left) | 1 | 0.194 | 0.194 | 0.194 | 0.194 |
| LBC1936 | FA | Intra-hemisphere (right) | 2 | 0.180 | 0.180 | 0.176 | 0.184 |
| LBC1936 | MD | Whole-brain | 7 | -0.127 | -0.164 | -0.219 | 0.162 |
| LBC1936 | MD | Inter-hemisphere | 4 | -0.180 | -0.172 | -0.219 | -0.159 |
| LBC1936 | MD | Intra-hemisphere | 3 | -0.056 | -0.162 | -0.166 | 0.162 |
| LBC1936 | MD | Intra-hemisphere (left) | 1 | -0.162 | -0.162 | -0.162 | -0.162 |
| LBC1936 | MD | Intra-hemisphere (right) | 2 | -0.002 | -0.002 | -0.166 | 0.162 |
| Meta-analysis | SC | Whole-brain | 391 | 0.046 | 0.044 | -0.071 | 0.109 |
| Meta-analysis | SC | Inter-hemisphere | 75 | 0.044 | 0.045 | -0.049 | 0.104 |
| Meta-analysis | SC | Intra-hemisphere | 315 | 0.046 | 0.044 | -0.071 | 0.109 |
| Meta-analysis | SC | Intra-hemisphere (left) | 158 | 0.044 | 0.040 | -0.050 | 0.103 |
| Meta-analysis | SC | Intra-hemisphere (right) | 157 | 0.049 | 0.049 | -0.071 | 0.109 |
| Meta-analysis | FA | Whole-brain | 433 | 0.053 | 0.051 | -0.033 | 0.134 |
| Meta-analysis | FA | Inter-hemisphere | 54 | 0.052 | 0.050 | 0.019 | 0.111 |
| Meta-analysis | FA | Intra-hemisphere | 378 | 0.053 | 0.051 | -0.033 | 0.134 |
| Meta-analysis | FA | Intra-hemisphere (left) | 180 | 0.053 | 0.051 | -0.026 | 0.134 |
| Meta-analysis | FA | Intra-hemisphere (right) | 198 | 0.053 | 0.053 | -0.033 | 0.117 |
| Meta-analysis | MD | Whole-brain | 325 | -0.062 | -0.060 | -0.146 | 0.021 |
| Meta-analysis | MD | Inter-hemisphere | 50 | -0.063 | -0.050 | -0.146 | -0.020 |
| Meta-analysis | MD | Intra-hemisphere | 272 | -0.063 | -0.061 | -0.139 | 0.021 |
| Meta-analysis | MD | Intra-hemisphere (left) | 141 | -0.060 | -0.059 | -0.121 | 0.021 |
| Meta-analysis | MD | Intra-hemisphere (right) | 131 | -0.065 | -0.063 | -0.139 | -0.019 |

**Supplementary Table 16**. Summary of β coefficients by intra- and inter-lobe connectivity for significant edge-g associations (p < 0.05, FDR), derived from the separate meta-analyses of SC, FA, and MD networks. Inter-lobe connectivity includes regions outside the designated lobe.

| **Study** | **Weighting** | **Lobe** | **Connectivity** | ***n*_edge_** | **Mean** | **Median** | **Min** | **Max** |
| --- | --- | --- | --- | --- | --- | --- | --- | --- |
| UKB | SC | Frontal | Intra | 57 | 0.038 | 0.037 | 0.020 | 0.067 |
| UKB | SC | Frontal | Inter | 160 | 0.041 | 0.037 | -0.034 | 0.097 |
| UKB | SC | Cingulate | Intra | 7 | 0.036 | 0.029 | 0.027 | 0.068 |
| UKB | SC | Cingulate | Inter | 40 | 0.019 | 0.026 | -0.050 | 0.059 |
| UKB | SC | Insula | Intra | 0 | - | - | - | - |
| UKB | SC | Insula | Inter | 34 | 0.031 | 0.034 | -0.035 | 0.086 |
| UKB | SC | Temporal | Intra | 27 | 0.042 | 0.042 | -0.022 | 0.087 |
| UKB | SC | Temporal | Inter | 155 | 0.042 | 0.042 | -0.031 | 0.089 |
| UKB | SC | Parietal | Intra | 28 | 0.048 | 0.043 | 0.018 | 0.087 |
| UKB | SC | Parietal | Inter | 178 | 0.044 | 0.043 | -0.032 | 0.093 |
| UKB | SC | Occipital | Intra | 0 | - | - | - | - |
| UKB | SC | Occipital | Inter | 44 | 0.041 | 0.043 | -0.024 | 0.100 |
| UKB | SC | Subcortical | Intra | 33 | 0.030 | 0.032 | -0.049 | 0.081 |
| UKB | SC | Subcortical | Inter | 193 | 0.043 | 0.042 | -0.050 | 0.100 |
| UKB | FA | Frontal | Intra | 72 | 0.042 | 0.041 | 0.018 | 0.069 |
| UKB | FA | Frontal | Inter | 185 | 0.043 | 0.042 | 0.017 | 0.091 |
| UKB | FA | Cingulate | Intra | 5 | 0.025 | 0.024 | 0.018 | 0.040 |
| UKB | FA | Cingulate | Inter | 53 | 0.032 | 0.030 | 0.018 | 0.048 |
| UKB | FA | Insula | Intra | 0 | - | - | - | - |
| UKB | FA | Insula | Inter | 42 | 0.029 | 0.031 | -0.033 | 0.064 |
| UKB | FA | Temporal | Intra | 29 | 0.033 | 0.033 | -0.020 | 0.059 |
| UKB | FA | Temporal | Inter | 178 | 0.037 | 0.036 | -0.018 | 0.074 |
| UKB | FA | Parietal | Intra | 29 | 0.044 | 0.043 | 0.017 | 0.081 |
| UKB | FA | Parietal | Inter | 201 | 0.042 | 0.041 | -0.018 | 0.091 |
| UKB | FA | Occipital | Intra | 12 | 0.046 | 0.043 | 0.028 | 0.062 |
| UKB | FA | Occipital | Inter | 71 | 0.049 | 0.048 | 0.017 | 0.084 |
| UKB | FA | Subcortical | Intra | 25 | 0.027 | 0.028 | -0.029 | 0.081 |
| UKB | FA | Subcortical | Inter | 200 | 0.039 | 0.037 | -0.033 | 0.084 |
| UKB | MD | Frontal | Intra | 73 | -0.059 | -0.062 | -0.096 | -0.018 |
| UKB | MD | Frontal | Inter | 202 | -0.064 | -0.064 | -0.104 | -0.017 |
| UKB | MD | Cingulate | Intra | 11 | -0.036 | -0.035 | -0.063 | -0.022 |
| UKB | MD | Cingulate | Inter | 59 | -0.046 | -0.043 | -0.085 | -0.018 |
| UKB | MD | Insula | Intra | 0 | - | - | - | - |
| UKB | MD | Insula | Inter | 48 | -0.041 | -0.038 | -0.087 | 0.018 |
| UKB | MD | Temporal | Intra | 20 | -0.025 | -0.027 | -0.037 | 0.018 |
| UKB | MD | Temporal | Inter | 144 | -0.036 | -0.035 | -0.073 | 0.024 |
| UKB | MD | Parietal | Intra | 30 | -0.047 | -0.049 | -0.078 | -0.019 |
| UKB | MD | Parietal | Inter | 200 | -0.050 | -0.050 | -0.098 | -0.018 |
| UKB | MD | Occipital | Intra | 1 | -0.018 | -0.018 | -0.018 | -0.018 |
| UKB | MD | Occipital | Inter | 32 | -0.022 | -0.022 | -0.037 | 0.024 |
| UKB | MD | Subcortical | Intra | 27 | -0.036 | -0.033 | -0.093 | 0.023 |
| UKB | MD | Subcortical | Inter | 209 | -0.053 | -0.050 | -0.104 | 0.018 |
| STRADL | SC | Frontal | Intra | 17 | 0.115 | 0.108 | 0.098 | 0.162 |
| STRADL | SC | Frontal | Inter | 35 | 0.122 | 0.119 | 0.099 | 0.166 |
| STRADL | SC | Cingulate | Intra | 3 | 0.156 | 0.155 | 0.122 | 0.191 |
| STRADL | SC | Cingulate | Inter | 10 | 0.126 | 0.116 | 0.100 | 0.179 |
| STRADL | SC | Insula | Intra | 0 | - | - | - | - |
| STRADL | SC | Insula | Inter | 10 | 0.113 | 0.112 | 0.099 | 0.128 |
| STRADL | SC | Temporal | Intra | 2 | 0.108 | 0.108 | 0.106 | 0.111 |
| STRADL | SC | Temporal | Inter | 20 | 0.114 | 0.106 | 0.097 | 0.157 |
| STRADL | SC | Parietal | Intra | 7 | 0.110 | 0.107 | 0.100 | 0.125 |
| STRADL | SC | Parietal | Inter | 45 | 0.121 | 0.116 | 0.097 | 0.168 |
| STRADL | SC | Occipital | Intra | 1 | 0.100 | 0.100 | 0.100 | 0.100 |
| STRADL | SC | Occipital | Inter | 5 | 0.115 | 0.112 | 0.106 | 0.123 |
| STRADL | SC | Subcortical | Intra | 3 | 0.141 | 0.127 | 0.116 | 0.178 |
| STRADL | SC | Subcortical | Inter | 27 | 0.117 | 0.110 | 0.098 | 0.179 |
| STRADL | FA | Frontal | Intra | 43 | 0.126 | 0.119 | 0.086 | 0.240 |
| STRADL | FA | Frontal | Inter | 118 | 0.129 | 0.125 | 0.082 | 0.203 |
| STRADL | FA | Cingulate | Intra | 1 | 0.094 | 0.094 | 0.094 | 0.094 |
| STRADL | FA | Cingulate | Inter | 37 | 0.123 | 0.119 | 0.082 | 0.203 |
| STRADL | FA | Insula | Intra | 0 | - | - | - | - |
| STRADL | FA | Insula | Inter | 18 | 0.114 | 0.117 | 0.088 | 0.158 |
| STRADL | FA | Temporal | Intra | 9 | 0.102 | 0.096 | 0.084 | 0.160 |
| STRADL | FA | Temporal | Inter | 129 | 0.129 | 0.124 | 0.083 | 0.190 |
| STRADL | FA | Parietal | Intra | 21 | 0.119 | 0.112 | 0.085 | 0.196 |
| STRADL | FA | Parietal | Inter | 154 | 0.130 | 0.124 | 0.083 | 0.225 |
| STRADL | FA | Occipital | Intra | 2 | 0.114 | 0.114 | 0.087 | 0.141 |
| STRADL | FA | Occipital | Inter | 43 | 0.134 | 0.114 | 0.084 | 0.236 |
| STRADL | FA | Subcortical | Intra | 9 | 0.119 | 0.112 | 0.100 | 0.154 |
| STRADL | FA | Subcortical | Inter | 149 | 0.133 | 0.124 | 0.083 | 0.236 |
| STRADL | MD | Frontal | Intra | 49 | -0.155 | -0.161 | -0.236 | -0.094 |
| STRADL | MD | Frontal | Inter | 97 | -0.159 | -0.162 | -0.229 | 0.091 |
| STRADL | MD | Cingulate | Intra | 3 | -0.178 | -0.172 | -0.202 | -0.160 |
| STRADL | MD | Cingulate | Inter | 37 | -0.159 | -0.157 | -0.216 | -0.101 |
| STRADL | MD | Insula | Intra | 0 | - | - | - | - |
| STRADL | MD | Insula | Inter | 21 | -0.150 | -0.147 | -0.238 | -0.102 |
| STRADL | MD | Temporal | Intra | 7 | -0.148 | -0.155 | -0.194 | -0.106 |
| STRADL | MD | Temporal | Inter | 64 | -0.143 | -0.147 | -0.243 | 0.120 |
| STRADL | MD | Parietal | Intra | 17 | -0.124 | -0.149 | -0.220 | 0.124 |
| STRADL | MD | Parietal | Inter | 104 | -0.161 | -0.168 | -0.278 | 0.110 |
| STRADL | MD | Occipital | Intra | 1 | -0.110 | -0.110 | -0.110 | -0.110 |
| STRADL | MD | Occipital | Inter | 17 | -0.116 | -0.143 | -0.229 | 0.120 |
| STRADL | MD | Subcortical | Intra | 11 | -0.128 | -0.131 | -0.189 | -0.097 |
| STRADL | MD | Subcortical | Inter | 124 | -0.163 | -0.168 | -0.278 | 0.110 |
| LBC1936 | SC | Frontal | Intra | 1 | 0.165 | 0.165 | 0.165 | 0.165 |
| LBC1936 | SC | Frontal | Inter | 3 | 0.062 | 0.168 | -0.150 | 0.168 |
| LBC1936 | SC | Cingulate | Intra | 1 | 0.158 | 0.158 | 0.158 | 0.158 |
| LBC1936 | SC | Cingulate | Inter | 2 | -0.151 | -0.151 | -0.152 | -0.150 |
| LBC1936 | SC | Insula | Intra | 0 | - | - | - | - |
| LBC1936 | SC | Insula | Inter | 1 | -0.150 | -0.150 | -0.150 | -0.150 |
| LBC1936 | SC | Temporal | Intra | 0 | - | - | - | - |
| LBC1936 | SC | Temporal | Inter | 1 | 0.156 | 0.156 | 0.156 | 0.156 |
| LBC1936 | SC | Parietal | Intra | 2 | 0.160 | 0.160 | 0.146 | 0.174 |
| LBC1936 | SC | Parietal | Inter | 2 | 0.160 | 0.160 | 0.152 | 0.168 |
| LBC1936 | SC | Occipital | Intra | 0 | - | - | - | - |
| LBC1936 | SC | Occipital | Inter | 1 | 0.152 | 0.152 | 0.152 | 0.152 |
| LBC1936 | SC | Subcortical | Intra | 5 | 0.121 | 0.158 | -0.152 | 0.229 |
| LBC1936 | SC | Subcortical | Inter | 4 | 0.005 | 0.003 | -0.152 | 0.168 |
| LBC1936 | FA | Frontal | Intra | 1 | 0.172 | 0.172 | 0.172 | 0.172 |
| LBC1936 | FA | Frontal | Inter | 2 | 0.189 | 0.189 | 0.184 | 0.194 |
| LBC1936 | FA | Cingulate | Intra | 0 | - | - | - | - |
| LBC1936 | FA | Cingulate | Inter | 0 | - | - | - | - |
| LBC1936 | FA | Insula | Intra | 0 | - | - | - | - |
| LBC1936 | FA | Insula | Inter | 0 | - | - | - | - |
| LBC1936 | FA | Temporal | Intra | 1 | 0.176 | 0.176 | 0.176 | 0.176 |
| LBC1936 | FA | Temporal | Inter | 0 | - | - | - | - |
| LBC1936 | FA | Parietal | Intra | 0 | - | - | - | - |
| LBC1936 | FA | Parietal | Inter | 0 | - | - | - | - |
| LBC1936 | FA | Occipital | Intra | 0 | - | - | - | - |
| LBC1936 | FA | Occipital | Inter | 0 | - | - | - | - |
| LBC1936 | FA | Subcortical | Intra | 0 | - | - | - | - |
| LBC1936 | FA | Subcortical | Inter | 2 | 0.189 | 0.189 | 0.184 | 0.194 |
| LBC1936 | MD | Frontal | Intra | 1 | -0.164 | -0.164 | -0.164 | -0.164 |
| LBC1936 | MD | Frontal | Inter | 3 | -0.079 | -0.180 | -0.219 | 0.162 |
| LBC1936 | MD | Cingulate | Intra | 0 | - | - | - | - |
| LBC1936 | MD | Cingulate | Inter | 1 | -0.219 | -0.219 | -0.219 | -0.219 |
| LBC1936 | MD | Insula | Intra | 0 | - | - | - | - |
| LBC1936 | MD | Insula | Inter | 0 | - | - | - | - |
| LBC1936 | MD | Temporal | Intra | 0 | - | - | - | - |
| LBC1936 | MD | Temporal | Inter | 0 | - | - | - | - |
| LBC1936 | MD | Parietal | Intra | 1 | -0.166 | -0.166 | -0.166 | -0.166 |
| LBC1936 | MD | Parietal | Inter | 1 | -0.162 | -0.162 | -0.162 | -0.162 |
| LBC1936 | MD | Occipital | Intra | 0 | - | - | - | - |
| LBC1936 | MD | Occipital | Inter | 1 | -0.162 | -0.162 | -0.162 | -0.162 |
| LBC1936 | MD | Subcortical | Intra | 1 | -0.159 | -0.159 | -0.159 | -0.159 |
| LBC1936 | MD | Subcortical | Inter | 2 | -0.009 | -0.009 | -0.180 | 0.162 |
| Meta-analysis | SC | Frontal | Intra | 38 | 0.044 | 0.043 | 0.019 | 0.104 |
| Meta-analysis | SC | Frontal | Inter | 112 | 0.048 | 0.043 | 0.019 | 0.097 |
| Meta-analysis | SC | Cingulate | Intra | 4 | 0.056 | 0.046 | 0.029 | 0.103 |
| Meta-analysis | SC | Cingulate | Inter | 27 | 0.024 | 0.027 | -0.071 | 0.102 |
| Meta-analysis | SC | Insula | Intra | 0 | - | - | - | - |
| Meta-analysis | SC | Insula | Inter | 14 | 0.035 | 0.036 | -0.029 | 0.086 |
| Meta-analysis | SC | Temporal | Intra | 19 | 0.047 | 0.046 | -0.027 | 0.088 |
| Meta-analysis | SC | Temporal | Inter | 113 | 0.045 | 0.045 | -0.029 | 0.095 |
| Meta-analysis | SC | Parietal | Intra | 21 | 0.054 | 0.051 | 0.022 | 0.099 |
| Meta-analysis | SC | Parietal | Inter | 130 | 0.051 | 0.049 | -0.031 | 0.102 |
| Meta-analysis | SC | Occipital | Intra | 0 | - | - | - | - |
| Meta-analysis | SC | Occipital | Inter | 36 | 0.047 | 0.044 | 0.019 | 0.097 |
| Meta-analysis | SC | Subcortical | Intra | 21 | 0.034 | 0.029 | -0.050 | 0.109 |
| Meta-analysis | SC | Subcortical | Inter | 144 | 0.047 | 0.049 | -0.071 | 0.097 |
| Meta-analysis | FA | Frontal | Intra | 56 | 0.055 | 0.056 | 0.020 | 0.111 |
| Meta-analysis | FA | Frontal | Inter | 126 | 0.060 | 0.060 | 0.020 | 0.117 |
| Meta-analysis | FA | Cingulate | Intra | 4 | 0.028 | 0.027 | 0.019 | 0.038 |
| Meta-analysis | FA | Cingulate | Inter | 27 | 0.041 | 0.039 | 0.020 | 0.113 |
| Meta-analysis | FA | Insula | Intra | 0 | - | - | - | - |
| Meta-analysis | FA | Insula | Inter | 28 | 0.036 | 0.037 | -0.033 | 0.077 |
| Meta-analysis | FA | Temporal | Intra | 24 | 0.041 | 0.039 | -0.019 | 0.086 |
| Meta-analysis | FA | Temporal | Inter | 105 | 0.050 | 0.049 | 0.018 | 0.101 |
| Meta-analysis | FA | Parietal | Intra | 23 | 0.055 | 0.054 | 0.022 | 0.087 |
| Meta-analysis | FA | Parietal | Inter | 137 | 0.058 | 0.058 | 0.019 | 0.107 |
| Meta-analysis | FA | Occipital | Intra | 9 | 0.049 | 0.044 | 0.029 | 0.082 |
| Meta-analysis | FA | Occipital | Inter | 52 | 0.060 | 0.054 | 0.019 | 0.134 |
| Meta-analysis | FA | Subcortical | Intra | 13 | 0.029 | 0.029 | -0.027 | 0.069 |
| Meta-analysis | FA | Subcortical | Inter | 133 | 0.054 | 0.053 | -0.033 | 0.134 |
| Meta-analysis | MD | Frontal | Intra | 41 | -0.069 | -0.071 | -0.135 | -0.020 |
| Meta-analysis | MD | Frontal | Inter | 122 | -0.074 | -0.069 | -0.146 | -0.030 |
| Meta-analysis | MD | Cingulate | Intra | 5 | -0.039 | -0.035 | -0.062 | -0.021 |
| Meta-analysis | MD | Cingulate | Inter | 40 | -0.063 | -0.054 | -0.139 | -0.024 |
| Meta-analysis | MD | Insula | Intra | 0 | - | - | - | - |
| Meta-analysis | MD | Insula | Inter | 27 | -0.052 | -0.047 | -0.105 | -0.021 |
| Meta-analysis | MD | Temporal | Intra | 9 | -0.027 | -0.027 | -0.038 | -0.019 |
| Meta-analysis | MD | Temporal | Inter | 62 | -0.043 | -0.039 | -0.076 | -0.020 |
| Meta-analysis | MD | Parietal | Intra | 18 | -0.067 | -0.068 | -0.110 | -0.032 |
| Meta-analysis | MD | Parietal | Inter | 97 | -0.065 | -0.064 | -0.139 | -0.020 |
| Meta-analysis | MD | Occipital | Intra | 1 | -0.020 | -0.020 | -0.020 | -0.020 |
| Meta-analysis | MD | Occipital | Inter | 8 | -0.032 | -0.026 | -0.077 | -0.020 |
| Meta-analysis | MD | Subcortical | Intra | 16 | -0.043 | -0.034 | -0.112 | 0.021 |
| Meta-analysis | MD | Subcortical | Inter | 114 | -0.070 | -0.072 | -0.146 | -0.020 |

**Supplementary Table 17**. Correlations (Pearson’s r) between the meta-analytic node-*g* β coefficients (derived from SC, FA, and MD networks) and cortical morphometry-*g* β coefficients (volume, surface area, and thickness) across 68 cortical regions from the Desikan-Killiany atlas. Bold indicates significance (*p* < 0.05, uncorrected)

|  | **Volume** | | **Surface area** | | **Thickness** | |
| --- | --- | --- | --- | --- | --- | --- |
| **Weighting** | ***r*** | ***p*** | ***r*** | ***p*** | ***r*** | ***p*** |
| SC | **0.653** | **0.000** | **0.556** | **0.000** | **0.331** | **0.006** |
| FA | **0.428** | **0.000** | **0.346** | **0.004** | **0.267** | **0.028** |
| MD | **-0.406** | **0.001** | **-0.386** | **0.001** | -0.115 | 0.350 |

**Supplementary Table 18**. Moderation estimates for age (standardised estimates) in relation to *g* and global network metrics for the two cohorts with broad age ranges (UKB and STRADL), across weightings (SC, FA, and MD), and across network metrics (mean edge weight, global efficiency, mean clustering coefficient). SE and p-values were estimated by SEM, and q-values were adjusted by FDR.

| **Study** | **Weighting** | **Metric** | **β** | ***SE*** | ***p*** | ***q*** |
| --- | --- | --- | --- | --- | --- | --- |
| UKB | SC | Mean edge weight | 0.000 | 0.007 | 0.961 | 0.961 |
| UKB | SC | Efficiency | 0.011 | 0.007 | 0.105 | 0.135 |
| UKB | SC | Clustering | 0.006 | 0.007 | 0.364 | 0.409 |
| UKB | FA | Mean edge weight | 0.017 | 0.007 | 0.018 | 0.033 |
| UKB | FA | Efficiency | 0.013 | 0.007 | 0.065 | 0.097 |
| UKB | FA | Clustering | 0.017 | 0.007 | 0.017 | 0.033 |
| UKB | MD | Mean edge weight | -0.042 | 0.007 | 0.000 | 0.000 |
| UKB | MD | Efficiency | -0.047 | 0.007 | 0.000 | 0.000 |
| UKB | MD | Clustering | -0.041 | 0.007 | 0.000 | 0.000 |
| STRADL | SC | Mean edge weight | -0.005 | 0.033 | 0.869 | 0.874 |
| STRADL | SC | Efficiency | 0.005 | 0.033 | 0.874 | 0.874 |
| STRADL | SC | Clustering | 0.010 | 0.033 | 0.752 | 0.874 |
| STRADL | FA | Mean edge weight | 0.039 | 0.029 | 0.168 | 0.555 |
| STRADL | FA | Efficiency | 0.037 | 0.028 | 0.185 | 0.555 |
| STRADL | FA | Clustering | 0.039 | 0.028 | 0.160 | 0.555 |
| STRADL | MD | Mean edge weight | -0.024 | 0.030 | 0.418 | 0.665 |
| STRADL | MD | Efficiency | -0.025 | 0.030 | 0.408 | 0.665 |
| STRADL | MD | Clustering | -0.023 | 0.030 | 0.443 | 0.665 |

**Supplementary Table 19**. Moderation estimates for age (standardised estimates) in relation to g and local efficiency for the two cohorts with broad age ranges (UKB and STRADL), for 85 FreeSurfer regions in uncorrected SC networks. SE and p-values were estimated by SEM, and q-values were adjusted by FDR.

|  |  | **UKB** | | | | **STRADL** | | | |
| --- | --- | --- | --- | --- | --- | --- | --- | --- | --- |
| **Hemisphere** | **Node** | **β** | ***SE*** | ***p*** | ***q*** | **β** | ***SE*** | ***p*** | ***q*** |
| Left | Thalamus | 0.011 | 0.007 | 0.117 | 0.268 | -0.034 | 0.034 | 0.322 | 0.830 |
| Left | Caudate | -0.006 | 0.007 | 0.350 | 0.518 | 0.027 | 0.029 | 0.356 | 0.836 |
| Left | Putamen | 0.009 | 0.007 | 0.203 | 0.373 | 0.031 | 0.033 | 0.344 | 0.836 |
| Left | Pallidum | -0.021 | 0.007 | 0.004 | 0.018 | -0.045 | 0.034 | 0.182 | 0.768 |
| Left | Hippocampus | 0.035 | 0.007 | 0.000 | 0.000 | 0.035 | 0.033 | 0.291 | 0.820 |
| Left | Amygdala | 0.021 | 0.007 | 0.003 | 0.015 | 0.091 | 0.033 | 0.005 | 0.183 |
| Left | Accumbens-area | -0.010 | 0.007 | 0.167 | 0.331 | 0.025 | 0.031 | 0.416 | 0.852 |
| Left | Ventral diencephalon | 0.004 | 0.007 | 0.532 | 0.675 | 0.002 | 0.036 | 0.966 | 0.991 |
| Left | Banks of superior temporal | -0.003 | 0.007 | 0.683 | 0.792 | 0.039 | 0.031 | 0.203 | 0.768 |
| Left | Caudal anterior cingulate | 0.002 | 0.007 | 0.739 | 0.827 | -0.028 | 0.033 | 0.395 | 0.845 |
| Left | Caudal middle frontal | -0.021 | 0.007 | 0.003 | 0.015 | -0.029 | 0.031 | 0.364 | 0.836 |
| Left | Cuneus | -0.002 | 0.007 | 0.783 | 0.854 | 0.063 | 0.032 | 0.048 | 0.342 |
| Left | Entorhinal | 0.023 | 0.007 | 0.001 | 0.009 | 0.081 | 0.030 | 0.007 | 0.183 |
| Left | Fusiform | 0.025 | 0.007 | 0.000 | 0.004 | 0.076 | 0.031 | 0.013 | 0.183 |
| Left | Inferior parietal | 0.008 | 0.007 | 0.236 | 0.410 | -0.003 | 0.033 | 0.930 | 0.991 |
| Left | Inferior temporal | -0.001 | 0.007 | 0.876 | 0.919 | -0.004 | 0.034 | 0.897 | 0.991 |
| Left | Isthmus cingulate | -0.027 | 0.007 | 0.000 | 0.002 | -0.034 | 0.035 | 0.334 | 0.834 |
| Left | Lateral occipital | 0.024 | 0.007 | 0.001 | 0.006 | -0.003 | 0.034 | 0.939 | 0.991 |
| Left | Lateral orbitofrontal | 0.008 | 0.007 | 0.252 | 0.428 | 0.019 | 0.033 | 0.578 | 0.927 |
| Left | Lingual | 0.004 | 0.007 | 0.625 | 0.759 | 0.034 | 0.032 | 0.292 | 0.820 |
| Left | Medial orbitofrontal | 0.007 | 0.007 | 0.342 | 0.518 | 0.029 | 0.029 | 0.309 | 0.820 |
| Left | Middle temporal | 0.022 | 0.007 | 0.002 | 0.012 | -0.022 | 0.034 | 0.514 | 0.891 |
| Left | Parahippocampal | 0.037 | 0.007 | 0.000 | 0.000 | 0.076 | 0.031 | 0.012 | 0.183 |
| Left | Paracentral | 0.012 | 0.007 | 0.096 | 0.232 | -0.020 | 0.032 | 0.541 | 0.909 |
| Left | Pars opercularis | 0.008 | 0.007 | 0.275 | 0.448 | -0.026 | 0.033 | 0.434 | 0.858 |
| Left | Pars orbitalis | -0.001 | 0.007 | 0.906 | 0.927 | -0.021 | 0.031 | 0.491 | 0.870 |
| Left | Pars triangularis | 0.013 | 0.007 | 0.062 | 0.165 | -0.001 | 0.033 | 0.981 | 0.991 |
| Left | Pericalcarine | -0.012 | 0.007 | 0.094 | 0.232 | 0.007 | 0.031 | 0.814 | 0.991 |
| Left | Postcentral | -0.005 | 0.007 | 0.479 | 0.635 | 0.002 | 0.033 | 0.955 | 0.991 |
| Left | Posterior cingulate | 0.003 | 0.007 | 0.651 | 0.777 | -0.074 | 0.035 | 0.035 | 0.274 |
| Left | Precentral | -0.000 | 0.007 | 0.984 | 0.984 | -0.013 | 0.035 | 0.717 | 0.979 |
| Left | Precuneus | 0.014 | 0.007 | 0.054 | 0.153 | -0.015 | 0.032 | 0.639 | 0.964 |
| Left | Rostral anterior cingulate | -0.020 | 0.007 | 0.006 | 0.026 | -0.009 | 0.031 | 0.782 | 0.991 |
| Left | Rostral middle frontal | 0.008 | 0.007 | 0.235 | 0.410 | 0.019 | 0.032 | 0.553 | 0.909 |
| Left | Superior frontal | 0.005 | 0.007 | 0.522 | 0.673 | -0.050 | 0.034 | 0.144 | 0.767 |
| Left | Superior parietal | -0.006 | 0.007 | 0.422 | 0.579 | 0.007 | 0.035 | 0.851 | 0.991 |
| Left | Superior temporal | 0.002 | 0.007 | 0.799 | 0.860 | -0.030 | 0.035 | 0.397 | 0.845 |
| Left | Supramarginal | -0.009 | 0.007 | 0.206 | 0.373 | -0.053 | 0.034 | 0.120 | 0.730 |
| Left | Frontal pole | 0.006 | 0.007 | 0.365 | 0.521 | -0.005 | 0.034 | 0.886 | 0.991 |
| Left | Temporal pole | 0.002 | 0.007 | 0.736 | 0.827 | 0.024 | 0.033 | 0.471 | 0.859 |
| Left | Transverse temporal | 0.005 | 0.007 | 0.475 | 0.635 | -0.017 | 0.033 | 0.599 | 0.942 |
| Left | Insula | -0.013 | 0.007 | 0.068 | 0.176 | -0.073 | 0.033 | 0.028 | 0.274 |
| Right | Thalamus | 0.019 | 0.007 | 0.007 | 0.031 | 0.019 | 0.033 | 0.556 | 0.909 |
| Right | Caudate | 0.039 | 0.007 | 0.000 | 0.000 | 0.033 | 0.027 | 0.217 | 0.768 |
| Right | Putamen | -0.003 | 0.007 | 0.690 | 0.792 | -0.047 | 0.034 | 0.162 | 0.768 |
| Right | Pallidum | 0.007 | 0.007 | 0.339 | 0.518 | 0.002 | 0.033 | 0.945 | 0.991 |
| Right | Hippocampus | 0.028 | 0.007 | 0.000 | 0.001 | 0.035 | 0.034 | 0.307 | 0.820 |
| Right | Amygdala | -0.007 | 0.007 | 0.326 | 0.514 | -0.025 | 0.035 | 0.466 | 0.859 |
| Right | Accumbens-area | 0.014 | 0.007 | 0.053 | 0.153 | -0.035 | 0.029 | 0.234 | 0.797 |
| Right | Ventral diencephalon | 0.008 | 0.007 | 0.268 | 0.447 | -0.013 | 0.032 | 0.691 | 0.979 |
| Right | Banks of superior temporal | -0.018 | 0.007 | 0.010 | 0.039 | -0.070 | 0.032 | 0.028 | 0.274 |
| Right | Caudal anterior cingulate | 0.011 | 0.007 | 0.120 | 0.268 | -0.001 | 0.034 | 0.983 | 0.991 |
| Right | Caudal middle frontal | -0.019 | 0.007 | 0.006 | 0.027 | 0.027 | 0.033 | 0.421 | 0.852 |
| Right | Cuneus | -0.015 | 0.007 | 0.039 | 0.127 | -0.034 | 0.033 | 0.307 | 0.820 |
| Right | Entorhinal | 0.025 | 0.007 | 0.000 | 0.004 | 0.116 | 0.037 | 0.002 | 0.140 |
| Right | Fusiform | 0.008 | 0.007 | 0.279 | 0.448 | 0.089 | 0.035 | 0.012 | 0.183 |
| Right | Inferior parietal | 0.016 | 0.007 | 0.023 | 0.080 | 0.008 | 0.031 | 0.800 | 0.991 |
| Right | Inferior temporal | 0.006 | 0.007 | 0.375 | 0.523 | 0.001 | 0.033 | 0.978 | 0.991 |
| Right | Isthmus cingulate | -0.025 | 0.007 | 0.000 | 0.004 | -0.075 | 0.035 | 0.034 | 0.274 |
| Right | Lateral occipital | 0.010 | 0.007 | 0.146 | 0.303 | 0.070 | 0.032 | 0.030 | 0.274 |
| Right | Lateral orbitofrontal | 0.019 | 0.007 | 0.008 | 0.033 | 0.029 | 0.034 | 0.397 | 0.845 |
| Right | Lingual | -0.006 | 0.007 | 0.368 | 0.521 | 0.014 | 0.032 | 0.658 | 0.964 |
| Right | Medial orbitofrontal | 0.010 | 0.007 | 0.172 | 0.331 | -0.033 | 0.031 | 0.292 | 0.820 |
| Right | Middle temporal | 0.003 | 0.007 | 0.658 | 0.777 | 0.011 | 0.032 | 0.730 | 0.979 |
| Right | Parahippocampal | 0.029 | 0.007 | 0.000 | 0.001 | 0.041 | 0.032 | 0.208 | 0.768 |
| Right | Paracentral | 0.018 | 0.007 | 0.011 | 0.042 | -0.042 | 0.033 | 0.207 | 0.768 |
| Right | Pars opercularis | 0.017 | 0.007 | 0.018 | 0.062 | -0.009 | 0.035 | 0.805 | 0.991 |
| Right | Pars orbitalis | 0.021 | 0.007 | 0.003 | 0.016 | 0.001 | 0.032 | 0.966 | 0.991 |
| Right | Pars triangularis | -0.000 | 0.007 | 0.968 | 0.980 | -0.007 | 0.033 | 0.841 | 0.991 |
| Right | Pericalcarine | -0.001 | 0.007 | 0.893 | 0.925 | 0.015 | 0.035 | 0.675 | 0.973 |
| Right | Postcentral | -0.004 | 0.007 | 0.611 | 0.753 | 0.005 | 0.033 | 0.874 | 0.991 |
| Right | Posterior cingulate | 0.011 | 0.007 | 0.123 | 0.268 | -0.043 | 0.034 | 0.213 | 0.768 |
| Right | Precentral | 0.013 | 0.007 | 0.059 | 0.163 | -0.015 | 0.034 | 0.654 | 0.964 |
| Right | Precuneus | 0.021 | 0.007 | 0.003 | 0.015 | -0.000 | 0.033 | 0.991 | 0.991 |
| Right | Rostral anterior cingulate | -0.011 | 0.007 | 0.121 | 0.268 | -0.055 | 0.032 | 0.084 | 0.548 |
| Right | Rostral middle frontal | 0.014 | 0.007 | 0.047 | 0.143 | -0.025 | 0.033 | 0.453 | 0.859 |
| Right | Superior frontal | 0.002 | 0.007 | 0.749 | 0.827 | -0.038 | 0.033 | 0.250 | 0.816 |
| Right | Superior parietal | 0.010 | 0.007 | 0.141 | 0.301 | 0.049 | 0.033 | 0.144 | 0.767 |
| Right | Superior temporal | -0.001 | 0.007 | 0.856 | 0.909 | -0.003 | 0.032 | 0.923 | 0.991 |
| Right | Supramarginal | -0.004 | 0.007 | 0.579 | 0.724 | -0.011 | 0.033 | 0.737 | 0.979 |
| Right | Frontal pole | 0.010 | 0.007 | 0.160 | 0.325 | 0.009 | 0.028 | 0.764 | 0.991 |
| Right | Temporal pole | 0.010 | 0.007 | 0.175 | 0.331 | 0.013 | 0.036 | 0.712 | 0.979 |
| Right | Transverse temporal | -0.006 | 0.007 | 0.353 | 0.518 | 0.024 | 0.033 | 0.475 | 0.859 |
| Right | Insula | -0.014 | 0.007 | 0.047 | 0.143 | -0.044 | 0.032 | 0.164 | 0.768 |
|  | Brainstem | -0.005 | 0.007 | 0.485 | 0.635 | -0.016 | 0.033 | 0.632 | 0.964 |

**Supplementary Table 20**. Moderation estimates for age (standardised estimates) in relation to g and local efficiency for the two cohorts with broad age ranges (UKB and STRADL), for 85 FreeSurfer regions in FA networks. SE and p-values were estimated by SEM, and q-values were adjusted by FDR.

|  |  | **UKB** | | | | **STRADL** | | | |
| --- | --- | --- | --- | --- | --- | --- | --- | --- | --- |
| **Hemisphere** | **Node** | **β** | ***SE*** | ***p*** | ***q*** | **β** | ***SE*** | ***p*** | ***q*** |
| Left | Thalamus | -0.002 | 0.007 | 0.820 | 0.917 | 0.038 | 0.029 | 0.194 | 0.417 |
| Left | Caudate | 0.021 | 0.007 | 0.003 | 0.009 | 0.052 | 0.027 | 0.061 | 0.417 |
| Left | Putamen | 0.001 | 0.007 | 0.940 | 0.957 | 0.047 | 0.028 | 0.097 | 0.417 |
| Left | Pallidum | -0.013 | 0.007 | 0.063 | 0.122 | 0.055 | 0.030 | 0.065 | 0.417 |
| Left | Hippocampus | 0.007 | 0.007 | 0.299 | 0.423 | 0.028 | 0.028 | 0.327 | 0.455 |
| Left | Amygdala | 0.000 | 0.007 | 0.977 | 0.977 | 0.044 | 0.030 | 0.140 | 0.417 |
| Left | Accumbens-area | -0.010 | 0.007 | 0.169 | 0.256 | 0.043 | 0.028 | 0.133 | 0.417 |
| Left | Ventral diencephalon | -0.008 | 0.007 | 0.240 | 0.351 | 0.030 | 0.030 | 0.313 | 0.455 |
| Left | Banks of superior temporal | -0.005 | 0.007 | 0.499 | 0.642 | 0.045 | 0.030 | 0.125 | 0.417 |
| Left | Caudal anterior cingulate | 0.010 | 0.007 | 0.143 | 0.230 | 0.029 | 0.031 | 0.350 | 0.455 |
| Left | Caudal middle frontal | 0.010 | 0.007 | 0.162 | 0.251 | 0.031 | 0.030 | 0.300 | 0.455 |
| Left | Cuneus | -0.001 | 0.007 | 0.875 | 0.937 | 0.003 | 0.032 | 0.915 | 0.926 |
| Left | Entorhinal | 0.023 | 0.007 | 0.001 | 0.004 | 0.014 | 0.028 | 0.623 | 0.671 |
| Left | Fusiform | 0.012 | 0.007 | 0.088 | 0.166 | 0.044 | 0.027 | 0.105 | 0.417 |
| Left | Inferior parietal | 0.011 | 0.007 | 0.142 | 0.230 | 0.033 | 0.028 | 0.242 | 0.417 |
| Left | Inferior temporal | 0.001 | 0.007 | 0.903 | 0.939 | 0.038 | 0.029 | 0.186 | 0.417 |
| Left | Isthmus cingulate | -0.003 | 0.007 | 0.709 | 0.825 | 0.037 | 0.029 | 0.193 | 0.417 |
| Left | Lateral occipital | 0.018 | 0.007 | 0.010 | 0.024 | 0.027 | 0.028 | 0.334 | 0.455 |
| Left | Lateral orbitofrontal | 0.018 | 0.007 | 0.010 | 0.024 | 0.051 | 0.029 | 0.077 | 0.417 |
| Left | Lingual | 0.002 | 0.007 | 0.741 | 0.852 | 0.030 | 0.028 | 0.289 | 0.455 |
| Left | Medial orbitofrontal | 0.023 | 0.007 | 0.001 | 0.004 | 0.037 | 0.027 | 0.176 | 0.417 |
| Left | Middle temporal | 0.005 | 0.007 | 0.510 | 0.647 | 0.045 | 0.029 | 0.116 | 0.417 |
| Left | Parahippocampal | 0.006 | 0.007 | 0.428 | 0.587 | 0.027 | 0.031 | 0.371 | 0.455 |
| Left | Paracentral | 0.011 | 0.007 | 0.132 | 0.221 | 0.064 | 0.030 | 0.036 | 0.417 |
| Left | Pars opercularis | 0.014 | 0.007 | 0.047 | 0.096 | 0.038 | 0.029 | 0.190 | 0.417 |
| Left | Pars orbitalis | 0.039 | 0.007 | 0.000 | 0.000 | 0.049 | 0.027 | 0.072 | 0.417 |
| Left | Pars triangularis | 0.025 | 0.007 | 0.001 | 0.003 | 0.045 | 0.028 | 0.110 | 0.417 |
| Left | Pericalcarine | 0.003 | 0.007 | 0.703 | 0.825 | 0.036 | 0.031 | 0.244 | 0.417 |
| Left | Postcentral | 0.002 | 0.007 | 0.807 | 0.914 | 0.053 | 0.030 | 0.078 | 0.417 |
| Left | Posterior cingulate | 0.019 | 0.007 | 0.007 | 0.019 | 0.041 | 0.028 | 0.147 | 0.417 |
| Left | Precentral | 0.005 | 0.007 | 0.447 | 0.594 | 0.041 | 0.030 | 0.165 | 0.417 |
| Left | Precuneus | 0.003 | 0.007 | 0.650 | 0.778 | 0.036 | 0.029 | 0.210 | 0.417 |
| Left | Rostral anterior cingulate | -0.000 | 0.007 | 0.945 | 0.957 | 0.017 | 0.031 | 0.594 | 0.653 |
| Left | Rostral middle frontal | 0.030 | 0.007 | 0.000 | 0.000 | 0.035 | 0.028 | 0.217 | 0.417 |
| Left | Superior frontal | 0.029 | 0.007 | 0.000 | 0.000 | 0.041 | 0.028 | 0.145 | 0.417 |
| Left | Superior parietal | 0.005 | 0.007 | 0.519 | 0.649 | 0.037 | 0.029 | 0.206 | 0.417 |
| Left | Superior temporal | 0.001 | 0.007 | 0.864 | 0.937 | 0.046 | 0.029 | 0.111 | 0.417 |
| Left | Supramarginal | -0.003 | 0.007 | 0.634 | 0.778 | 0.037 | 0.029 | 0.204 | 0.417 |
| Left | Frontal pole | 0.030 | 0.007 | 0.000 | 0.000 | 0.044 | 0.029 | 0.127 | 0.417 |
| Left | Temporal pole | 0.005 | 0.007 | 0.444 | 0.594 | 0.037 | 0.030 | 0.213 | 0.417 |
| Left | Transverse temporal | -0.013 | 0.007 | 0.058 | 0.116 | 0.062 | 0.030 | 0.040 | 0.417 |
| Left | Insula | 0.001 | 0.007 | 0.906 | 0.939 | 0.041 | 0.029 | 0.160 | 0.417 |
| Right | Thalamus | 0.010 | 0.007 | 0.172 | 0.256 | 0.030 | 0.029 | 0.316 | 0.455 |
| Right | Caudate | 0.021 | 0.007 | 0.003 | 0.010 | 0.026 | 0.028 | 0.351 | 0.455 |
| Right | Putamen | 0.014 | 0.007 | 0.047 | 0.096 | 0.032 | 0.029 | 0.267 | 0.445 |
| Right | Pallidum | 0.011 | 0.007 | 0.109 | 0.194 | 0.069 | 0.031 | 0.025 | 0.417 |
| Right | Hippocampus | 0.021 | 0.007 | 0.003 | 0.010 | 0.013 | 0.029 | 0.654 | 0.686 |
| Right | Amygdala | 0.011 | 0.007 | 0.129 | 0.220 | 0.000 | 0.032 | 0.997 | 0.997 |
| Right | Accumbens-area | -0.008 | 0.007 | 0.262 | 0.377 | 0.033 | 0.033 | 0.313 | 0.455 |
| Right | Ventral diencephalon | 0.001 | 0.007 | 0.852 | 0.937 | 0.023 | 0.030 | 0.445 | 0.504 |
| Right | Banks of superior temporal | 0.005 | 0.007 | 0.497 | 0.642 | 0.035 | 0.029 | 0.235 | 0.417 |
| Right | Caudal anterior cingulate | 0.020 | 0.007 | 0.005 | 0.016 | 0.015 | 0.029 | 0.600 | 0.653 |
| Right | Caudal middle frontal | 0.030 | 0.007 | 0.000 | 0.000 | 0.026 | 0.032 | 0.419 | 0.488 |
| Right | Cuneus | -0.006 | 0.007 | 0.371 | 0.517 | -0.027 | 0.031 | 0.397 | 0.476 |
| Right | Entorhinal | 0.035 | 0.007 | 0.000 | 0.000 | 0.038 | 0.030 | 0.203 | 0.417 |
| Right | Fusiform | 0.025 | 0.007 | 0.000 | 0.002 | 0.056 | 0.029 | 0.053 | 0.417 |
| Right | Inferior parietal | 0.024 | 0.007 | 0.001 | 0.003 | 0.035 | 0.029 | 0.236 | 0.417 |
| Right | Inferior temporal | 0.019 | 0.007 | 0.008 | 0.021 | 0.040 | 0.029 | 0.162 | 0.417 |
| Right | Isthmus cingulate | 0.001 | 0.007 | 0.882 | 0.937 | 0.025 | 0.028 | 0.368 | 0.455 |
| Right | Lateral occipital | 0.026 | 0.007 | 0.000 | 0.002 | 0.039 | 0.028 | 0.169 | 0.417 |
| Right | Lateral orbitofrontal | 0.024 | 0.007 | 0.001 | 0.003 | 0.042 | 0.030 | 0.162 | 0.417 |
| Right | Lingual | 0.012 | 0.007 | 0.102 | 0.189 | 0.029 | 0.030 | 0.323 | 0.455 |
| Right | Medial orbitofrontal | 0.025 | 0.007 | 0.001 | 0.003 | 0.015 | 0.032 | 0.633 | 0.673 |
| Right | Middle temporal | 0.020 | 0.007 | 0.006 | 0.018 | 0.042 | 0.029 | 0.143 | 0.417 |
| Right | Parahippocampal | 0.019 | 0.007 | 0.008 | 0.021 | 0.024 | 0.029 | 0.418 | 0.488 |
| Right | Paracentral | 0.023 | 0.007 | 0.001 | 0.004 | 0.030 | 0.029 | 0.304 | 0.455 |
| Right | Pars opercularis | 0.030 | 0.007 | 0.000 | 0.000 | 0.028 | 0.030 | 0.364 | 0.455 |
| Right | Pars orbitalis | 0.036 | 0.007 | 0.000 | 0.000 | 0.043 | 0.029 | 0.139 | 0.417 |
| Right | Pars triangularis | 0.033 | 0.007 | 0.000 | 0.000 | 0.033 | 0.030 | 0.273 | 0.447 |
| Right | Pericalcarine | 0.015 | 0.007 | 0.041 | 0.089 | 0.021 | 0.030 | 0.467 | 0.522 |
| Right | Postcentral | 0.010 | 0.007 | 0.154 | 0.242 | 0.047 | 0.032 | 0.136 | 0.417 |
| Right | Posterior cingulate | 0.025 | 0.007 | 0.000 | 0.003 | 0.037 | 0.028 | 0.189 | 0.417 |
| Right | Precentral | 0.018 | 0.007 | 0.012 | 0.029 | 0.029 | 0.031 | 0.345 | 0.455 |
| Right | Precuneus | 0.017 | 0.007 | 0.015 | 0.034 | 0.034 | 0.028 | 0.230 | 0.417 |
| Right | Rostral anterior cingulate | 0.015 | 0.007 | 0.030 | 0.067 | 0.027 | 0.033 | 0.425 | 0.488 |
| Right | Rostral middle frontal | 0.031 | 0.007 | 0.000 | 0.000 | 0.026 | 0.030 | 0.375 | 0.455 |
| Right | Superior frontal | 0.031 | 0.007 | 0.000 | 0.000 | 0.034 | 0.029 | 0.245 | 0.417 |
| Right | Superior parietal | 0.011 | 0.007 | 0.110 | 0.194 | 0.029 | 0.029 | 0.312 | 0.455 |
| Right | Superior temporal | 0.011 | 0.007 | 0.119 | 0.207 | 0.043 | 0.030 | 0.150 | 0.417 |
| Right | Supramarginal | 0.021 | 0.007 | 0.003 | 0.010 | 0.027 | 0.030 | 0.364 | 0.455 |
| Right | Frontal pole | 0.032 | 0.007 | 0.000 | 0.000 | 0.040 | 0.030 | 0.181 | 0.417 |
| Right | Temporal pole | 0.026 | 0.007 | 0.000 | 0.002 | 0.037 | 0.031 | 0.222 | 0.417 |
| Right | Transverse temporal | -0.003 | 0.007 | 0.644 | 0.778 | -0.006 | 0.040 | 0.873 | 0.905 |
| Right | Insula | 0.014 | 0.007 | 0.048 | 0.096 | 0.042 | 0.029 | 0.151 | 0.417 |
|  | Brainstem | 0.022 | 0.007 | 0.002 | 0.007 | -0.004 | 0.033 | 0.908 | 0.926 |

**Supplementary Table 21**. Moderation estimates for age (standardised estimates) in relation to g and local efficiency for the two cohorts with broad age ranges (UKB and STRADL), for 85 FreeSurfer regions in MD networks. SE and p-values were estimated by SEM, and q-values were adjusted by FDR.

|  |  | **UKB** | | | | **STRADL** | | | |
| --- | --- | --- | --- | --- | --- | --- | --- | --- | --- |
| **Hemisphere** | **Node** | **β** | ***SE*** | ***p*** | ***q*** | **β** | ***SE*** | ***p*** | ***q*** |
| Left | Thalamus | -0.050 | 0.007 | 0.000 | 0.000 | -0.017 | 0.031 | 0.579 | 0.722 |
| Left | Caudate | -0.063 | 0.007 | 0.000 | 0.000 | -0.018 | 0.030 | 0.544 | 0.722 |
| Left | Putamen | -0.041 | 0.007 | 0.000 | 0.000 | -0.021 | 0.031 | 0.487 | 0.722 |
| Left | Pallidum | -0.037 | 0.007 | 0.000 | 0.000 | -0.019 | 0.031 | 0.547 | 0.722 |
| Left | Hippocampus | -0.015 | 0.007 | 0.038 | 0.045 | -0.044 | 0.031 | 0.157 | 0.722 |
| Left | Amygdala | -0.002 | 0.007 | 0.781 | 0.781 | -0.049 | 0.030 | 0.106 | 0.722 |
| Left | Accumbens-area | -0.003 | 0.007 | 0.657 | 0.665 | -0.079 | 0.031 | 0.011 | 0.455 |
| Left | Ventral diencephalon | -0.020 | 0.007 | 0.005 | 0.007 | -0.044 | 0.032 | 0.171 | 0.722 |
| Left | Banks of superior temporal | -0.024 | 0.007 | 0.001 | 0.001 | -0.016 | 0.031 | 0.607 | 0.737 |
| Left | Caudal anterior cingulate | -0.053 | 0.007 | 0.000 | 0.000 | -0.015 | 0.030 | 0.627 | 0.740 |
| Left | Caudal middle frontal | -0.047 | 0.007 | 0.000 | 0.000 | -0.008 | 0.032 | 0.813 | 0.856 |
| Left | Cuneus | -0.005 | 0.007 | 0.511 | 0.523 | -0.030 | 0.030 | 0.326 | 0.722 |
| Left | Entorhinal | 0.006 | 0.007 | 0.416 | 0.442 | -0.042 | 0.032 | 0.190 | 0.722 |
| Left | Fusiform | -0.006 | 0.007 | 0.383 | 0.412 | -0.040 | 0.031 | 0.187 | 0.722 |
| Left | Inferior parietal | -0.028 | 0.007 | 0.000 | 0.000 | -0.018 | 0.031 | 0.555 | 0.722 |
| Left | Inferior temporal | -0.026 | 0.007 | 0.000 | 0.000 | -0.026 | 0.031 | 0.412 | 0.722 |
| Left | Isthmus cingulate | -0.038 | 0.007 | 0.000 | 0.000 | -0.026 | 0.030 | 0.390 | 0.722 |
| Left | Lateral occipital | -0.022 | 0.007 | 0.002 | 0.003 | -0.034 | 0.030 | 0.259 | 0.722 |
| Left | Lateral orbitofrontal | -0.029 | 0.007 | 0.000 | 0.000 | -0.043 | 0.030 | 0.152 | 0.722 |
| Left | Lingual | 0.006 | 0.007 | 0.446 | 0.468 | -0.042 | 0.030 | 0.171 | 0.722 |
| Left | Medial orbitofrontal | -0.050 | 0.007 | 0.000 | 0.000 | -0.035 | 0.030 | 0.243 | 0.722 |
| Left | Middle temporal | -0.032 | 0.007 | 0.000 | 0.000 | -0.024 | 0.031 | 0.447 | 0.722 |
| Left | Parahippocampal | 0.017 | 0.007 | 0.018 | 0.022 | -0.030 | 0.032 | 0.346 | 0.722 |
| Left | Paracentral | -0.056 | 0.007 | 0.000 | 0.000 | -0.009 | 0.031 | 0.769 | 0.838 |
| Left | Pars opercularis | -0.049 | 0.007 | 0.000 | 0.000 | -0.001 | 0.031 | 0.968 | 0.968 |
| Left | Pars orbitalis | -0.053 | 0.007 | 0.000 | 0.000 | -0.028 | 0.030 | 0.352 | 0.722 |
| Left | Pars triangularis | -0.052 | 0.007 | 0.000 | 0.000 | -0.008 | 0.031 | 0.789 | 0.849 |
| Left | Pericalcarine | -0.010 | 0.007 | 0.178 | 0.196 | -0.040 | 0.031 | 0.197 | 0.722 |
| Left | Postcentral | -0.045 | 0.007 | 0.000 | 0.000 | -0.007 | 0.031 | 0.816 | 0.856 |
| Left | Posterior cingulate | -0.062 | 0.007 | 0.000 | 0.000 | -0.023 | 0.030 | 0.429 | 0.722 |
| Left | Precentral | -0.048 | 0.007 | 0.000 | 0.000 | -0.003 | 0.031 | 0.915 | 0.936 |
| Left | Precuneus | -0.035 | 0.007 | 0.000 | 0.000 | -0.020 | 0.031 | 0.508 | 0.722 |
| Left | Rostral anterior cingulate | -0.057 | 0.007 | 0.000 | 0.000 | -0.037 | 0.031 | 0.240 | 0.722 |
| Left | Rostral middle frontal | -0.064 | 0.007 | 0.000 | 0.000 | -0.012 | 0.030 | 0.700 | 0.794 |
| Left | Superior frontal | -0.062 | 0.007 | 0.000 | 0.000 | -0.015 | 0.030 | 0.626 | 0.740 |
| Left | Superior parietal | -0.034 | 0.007 | 0.000 | 0.000 | -0.019 | 0.031 | 0.550 | 0.722 |
| Left | Superior temporal | -0.033 | 0.007 | 0.000 | 0.000 | -0.024 | 0.031 | 0.435 | 0.722 |
| Left | Supramarginal | -0.032 | 0.007 | 0.000 | 0.000 | -0.013 | 0.031 | 0.665 | 0.763 |
| Left | Frontal pole | -0.062 | 0.007 | 0.000 | 0.000 | -0.017 | 0.030 | 0.572 | 0.722 |
| Left | Temporal pole | -0.006 | 0.007 | 0.378 | 0.412 | -0.061 | 0.030 | 0.045 | 0.722 |
| Left | Transverse temporal | -0.028 | 0.007 | 0.000 | 0.000 | -0.009 | 0.030 | 0.760 | 0.838 |
| Left | Insula | -0.031 | 0.007 | 0.000 | 0.000 | -0.022 | 0.030 | 0.453 | 0.722 |
| Right | Thalamus | -0.049 | 0.007 | 0.000 | 0.000 | -0.021 | 0.030 | 0.483 | 0.722 |
| Right | Caudate | -0.060 | 0.007 | 0.000 | 0.000 | -0.014 | 0.030 | 0.640 | 0.745 |
| Right | Putamen | -0.048 | 0.007 | 0.000 | 0.000 | -0.020 | 0.030 | 0.512 | 0.722 |
| Right | Pallidum | -0.046 | 0.007 | 0.000 | 0.000 | -0.030 | 0.031 | 0.333 | 0.722 |
| Right | Hippocampus | -0.022 | 0.007 | 0.003 | 0.004 | -0.039 | 0.030 | 0.197 | 0.722 |
| Right | Amygdala | -0.017 | 0.007 | 0.021 | 0.026 | -0.043 | 0.032 | 0.186 | 0.722 |
| Right | Accumbens-area | -0.026 | 0.007 | 0.000 | 0.000 | -0.085 | 0.031 | 0.006 | 0.455 |
| Right | Ventral diencephalon | -0.038 | 0.007 | 0.000 | 0.000 | -0.023 | 0.031 | 0.444 | 0.722 |
| Right | Banks of superior temporal | -0.032 | 0.007 | 0.000 | 0.000 | -0.020 | 0.030 | 0.503 | 0.722 |
| Right | Caudal anterior cingulate | -0.055 | 0.007 | 0.000 | 0.000 | -0.024 | 0.030 | 0.432 | 0.722 |
| Right | Caudal middle frontal | -0.057 | 0.007 | 0.000 | 0.000 | -0.010 | 0.032 | 0.749 | 0.838 |
| Right | Cuneus | -0.011 | 0.007 | 0.113 | 0.128 | -0.028 | 0.030 | 0.340 | 0.722 |
| Right | Entorhinal | -0.005 | 0.007 | 0.468 | 0.485 | -0.066 | 0.035 | 0.064 | 0.722 |
| Right | Fusiform | -0.016 | 0.007 | 0.030 | 0.036 | -0.044 | 0.030 | 0.141 | 0.722 |
| Right | Inferior parietal | -0.042 | 0.007 | 0.000 | 0.000 | -0.023 | 0.030 | 0.443 | 0.722 |
| Right | Inferior temporal | -0.029 | 0.007 | 0.000 | 0.000 | -0.037 | 0.030 | 0.228 | 0.722 |
| Right | Isthmus cingulate | -0.035 | 0.007 | 0.000 | 0.000 | -0.027 | 0.029 | 0.353 | 0.722 |
| Right | Lateral occipital | -0.029 | 0.007 | 0.000 | 0.000 | -0.033 | 0.030 | 0.278 | 0.722 |
| Right | Lateral orbitofrontal | -0.044 | 0.007 | 0.000 | 0.000 | -0.034 | 0.030 | 0.259 | 0.722 |
| Right | Lingual | -0.017 | 0.007 | 0.018 | 0.023 | -0.033 | 0.030 | 0.265 | 0.722 |
| Right | Medial orbitofrontal | -0.057 | 0.007 | 0.000 | 0.000 | -0.047 | 0.031 | 0.130 | 0.722 |
| Right | Middle temporal | -0.034 | 0.007 | 0.000 | 0.000 | -0.036 | 0.030 | 0.237 | 0.722 |
| Right | Parahippocampal | 0.010 | 0.007 | 0.167 | 0.187 | -0.033 | 0.030 | 0.279 | 0.722 |
| Right | Paracentral | -0.059 | 0.007 | 0.000 | 0.000 | -0.018 | 0.030 | 0.541 | 0.722 |
| Right | Pars opercularis | -0.055 | 0.007 | 0.000 | 0.000 | -0.018 | 0.031 | 0.549 | 0.722 |
| Right | Pars orbitalis | -0.054 | 0.007 | 0.000 | 0.000 | -0.029 | 0.030 | 0.331 | 0.722 |
| Right | Pars triangularis | -0.051 | 0.007 | 0.000 | 0.000 | -0.016 | 0.030 | 0.586 | 0.722 |
| Right | Pericalcarine | -0.014 | 0.007 | 0.047 | 0.054 | -0.030 | 0.031 | 0.330 | 0.722 |
| Right | Postcentral | -0.047 | 0.007 | 0.000 | 0.000 | -0.017 | 0.030 | 0.576 | 0.722 |
| Right | Posterior cingulate | -0.056 | 0.007 | 0.000 | 0.000 | -0.024 | 0.030 | 0.435 | 0.722 |
| Right | Precentral | -0.053 | 0.007 | 0.000 | 0.000 | -0.006 | 0.031 | 0.842 | 0.873 |
| Right | Precuneus | -0.041 | 0.007 | 0.000 | 0.000 | -0.020 | 0.031 | 0.509 | 0.722 |
| Right | Rostral anterior cingulate | -0.071 | 0.007 | 0.000 | 0.000 | -0.048 | 0.030 | 0.116 | 0.722 |
| Right | Rostral middle frontal | -0.065 | 0.007 | 0.000 | 0.000 | -0.017 | 0.030 | 0.570 | 0.722 |
| Right | Superior frontal | -0.065 | 0.007 | 0.000 | 0.000 | -0.019 | 0.030 | 0.520 | 0.722 |
| Right | Superior parietal | -0.040 | 0.007 | 0.000 | 0.000 | -0.019 | 0.031 | 0.526 | 0.722 |
| Right | Superior temporal | -0.036 | 0.007 | 0.000 | 0.000 | -0.031 | 0.030 | 0.311 | 0.722 |
| Right | Supramarginal | -0.037 | 0.007 | 0.000 | 0.000 | -0.018 | 0.030 | 0.540 | 0.722 |
| Right | Frontal pole | -0.060 | 0.007 | 0.000 | 0.000 | -0.026 | 0.030 | 0.382 | 0.722 |
| Right | Temporal pole | -0.015 | 0.007 | 0.045 | 0.052 | -0.080 | 0.034 | 0.019 | 0.526 |
| Right | Transverse temporal | -0.029 | 0.007 | 0.000 | 0.000 | -0.026 | 0.040 | 0.517 | 0.722 |
| Right | Insula | -0.044 | 0.007 | 0.000 | 0.000 | -0.035 | 0.030 | 0.240 | 0.722 |
|  | Brainstem | -0.027 | 0.007 | 0.000 | 0.000 | -0.003 | 0.035 | 0.925 | 0.936 |

**Supplementary Table 22**. Correlations (Pearson’s r) between predicted composite scores and actual g scores in the UKB held-out sample (n = 18,642), alongside those in the three samples contributing to the meta-analysis for comparison.

| **Sample** | **SC** | **FA** | **MD** |
| --- | --- | --- | --- |
| UKB held-out | 0.256 | 0.130 | 0.184 |
| UKB main | 0.262 | 0.140 | 0.192 |
| STRADL | 0.264 | 0.173 | 0.128 |
| LBC1936 | 0.187 | 0.151 | 0.084 |

### Figures

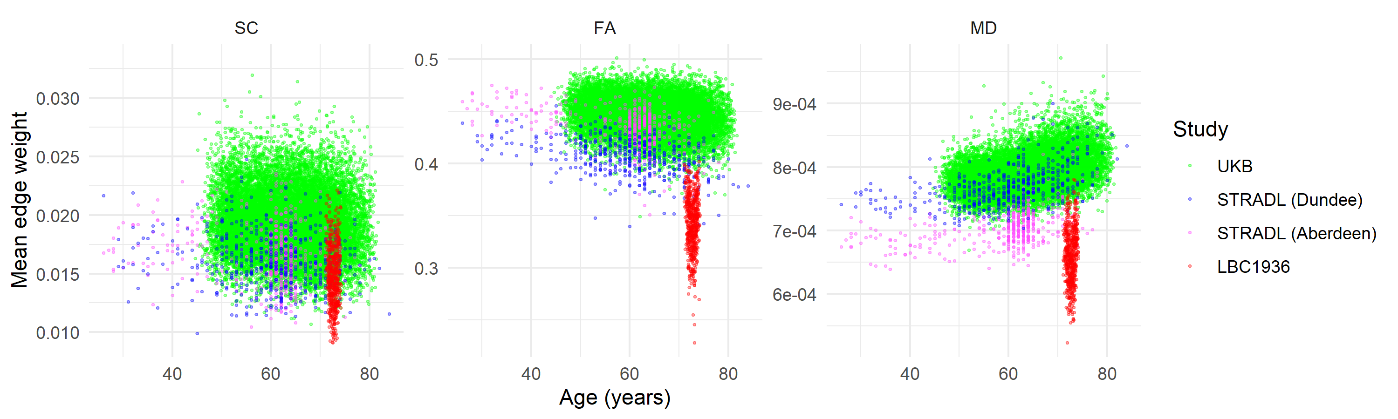

**Supplementary Fig. 1**. Scatter plots for three network weightings showing mean edge weight against age across the three cohorts, with the STRADL dataset separated into Aberdeen and Dundee scan sites. Network weightings are uncorrected streamline count (SC), fractional anisotropy (FA) and mean diffusivity (MD).

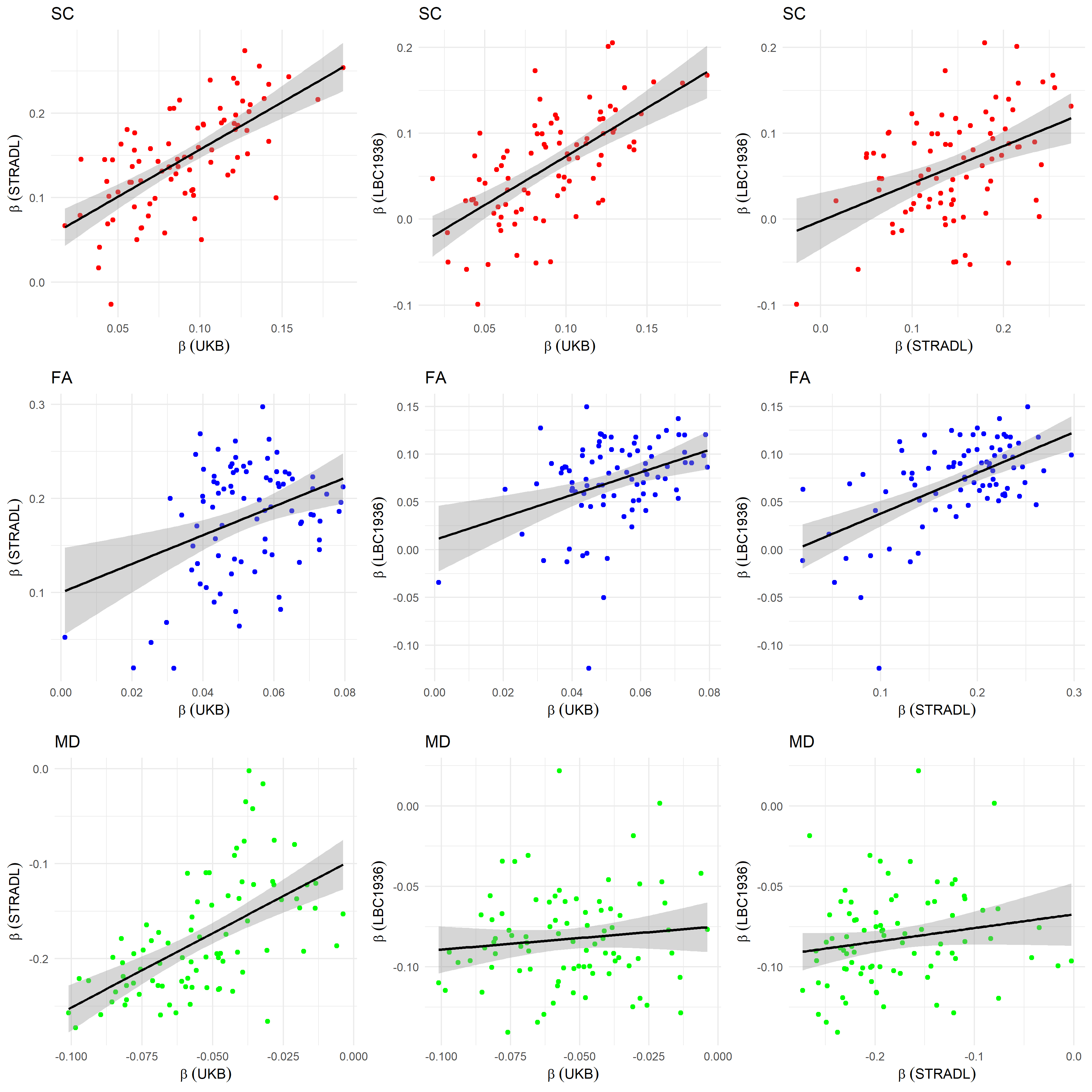

**Supplementary Fig. 2**. Scatter plots of the relationships between nodal β coefficients for the associations between *g* and nodal local efficiency, for each of the three possible pairings of cohorts across weighting schemes (SC, FA, and MD). Black line indicates linear fit.

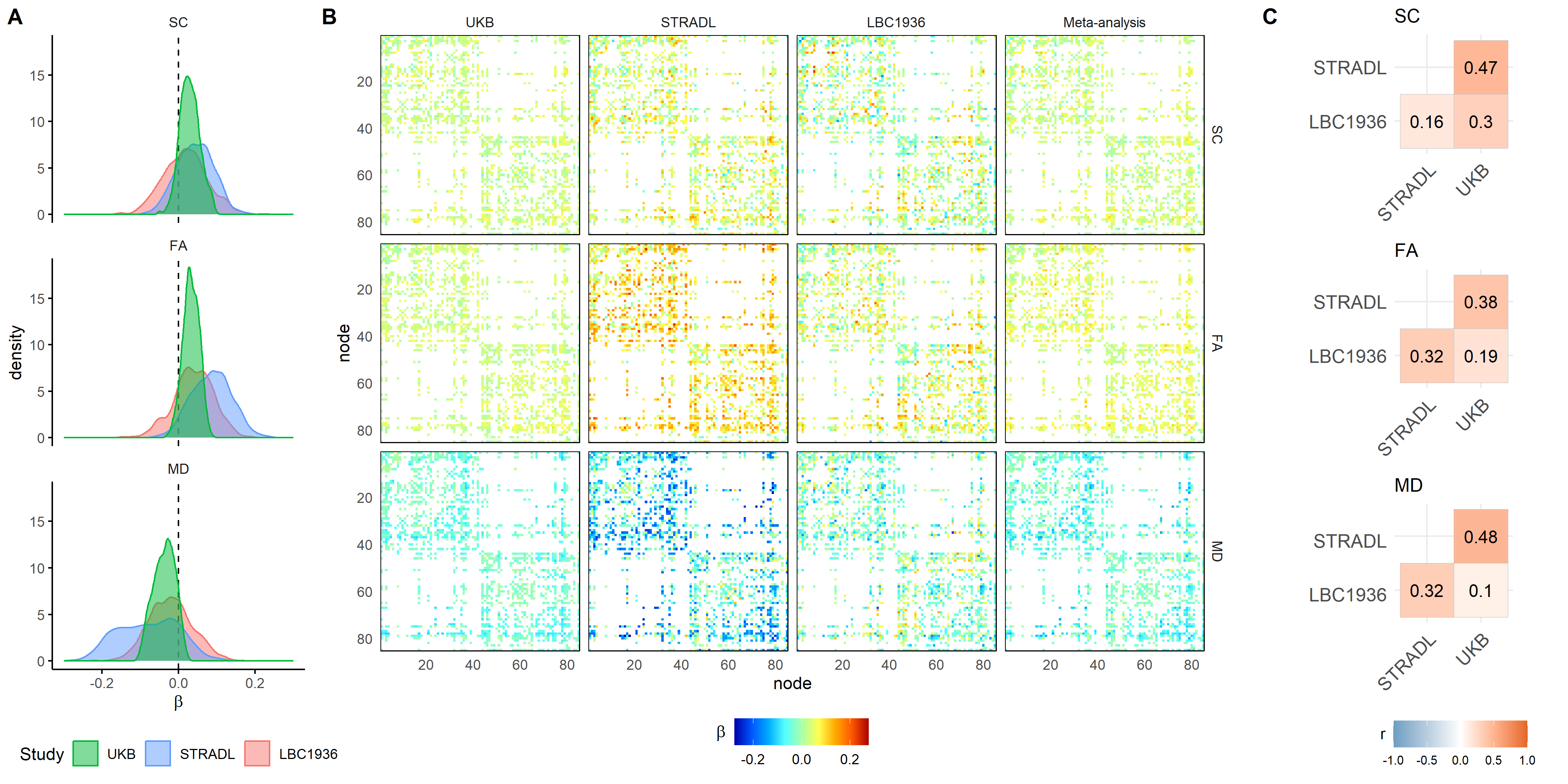

**Supplementary Fig. 3**. A) Histograms of all regression coefficients (standardised βs) for g ~ edge, corrected for age, sex and scanner. B) Connectivity matrices of the estimated βs for each cohort and the meta-analysis, where white indicates an edge was not present in the cross-short network mask. C) Correlations of the above β coefficients between cohorts.

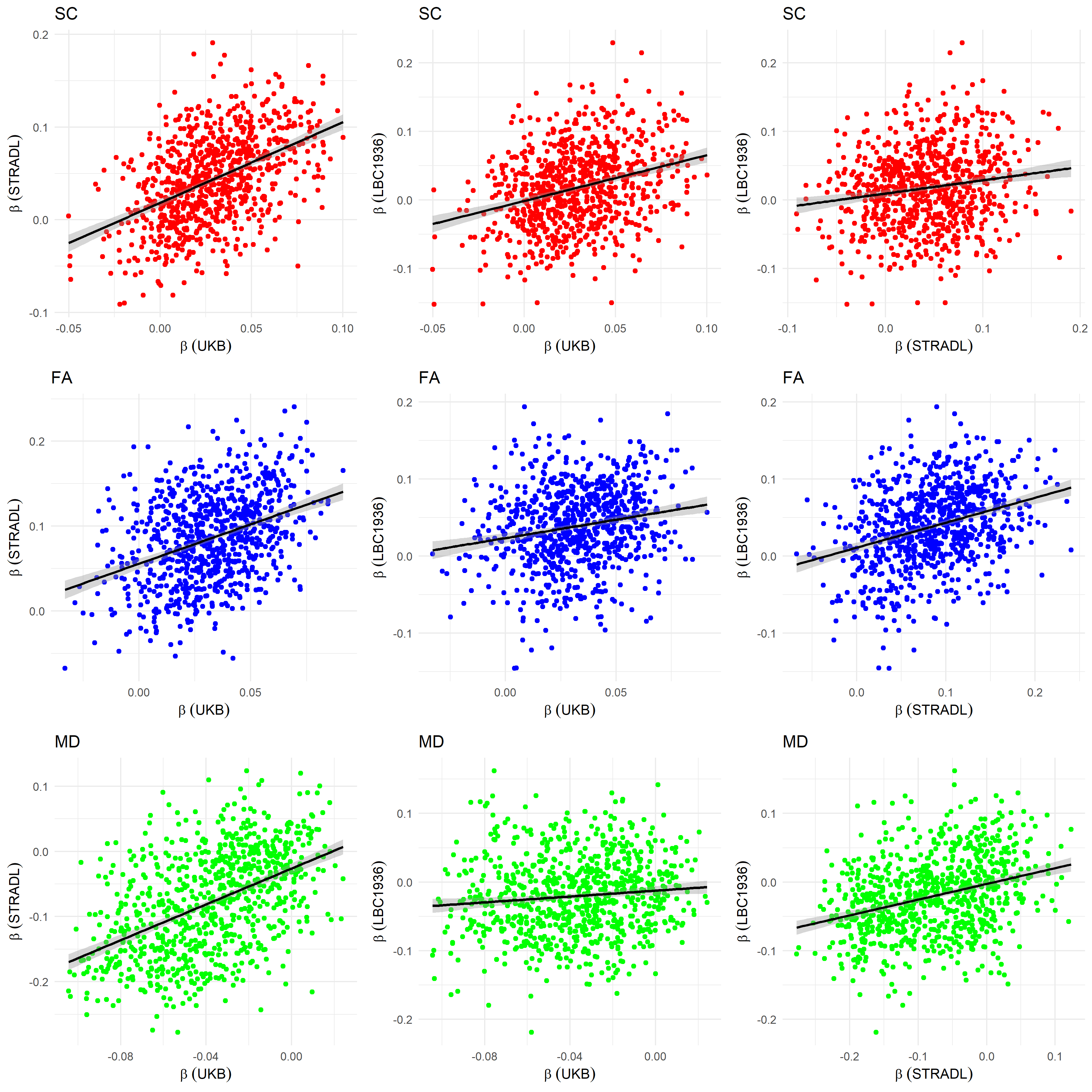

**Supplementary Fig. 4**. Scatter plots of the relationship between edge-g β coefficients for each of the three possible pairings of cohorts across weighting schemes (SC, FA, and MD). Black line indicates linear fit.

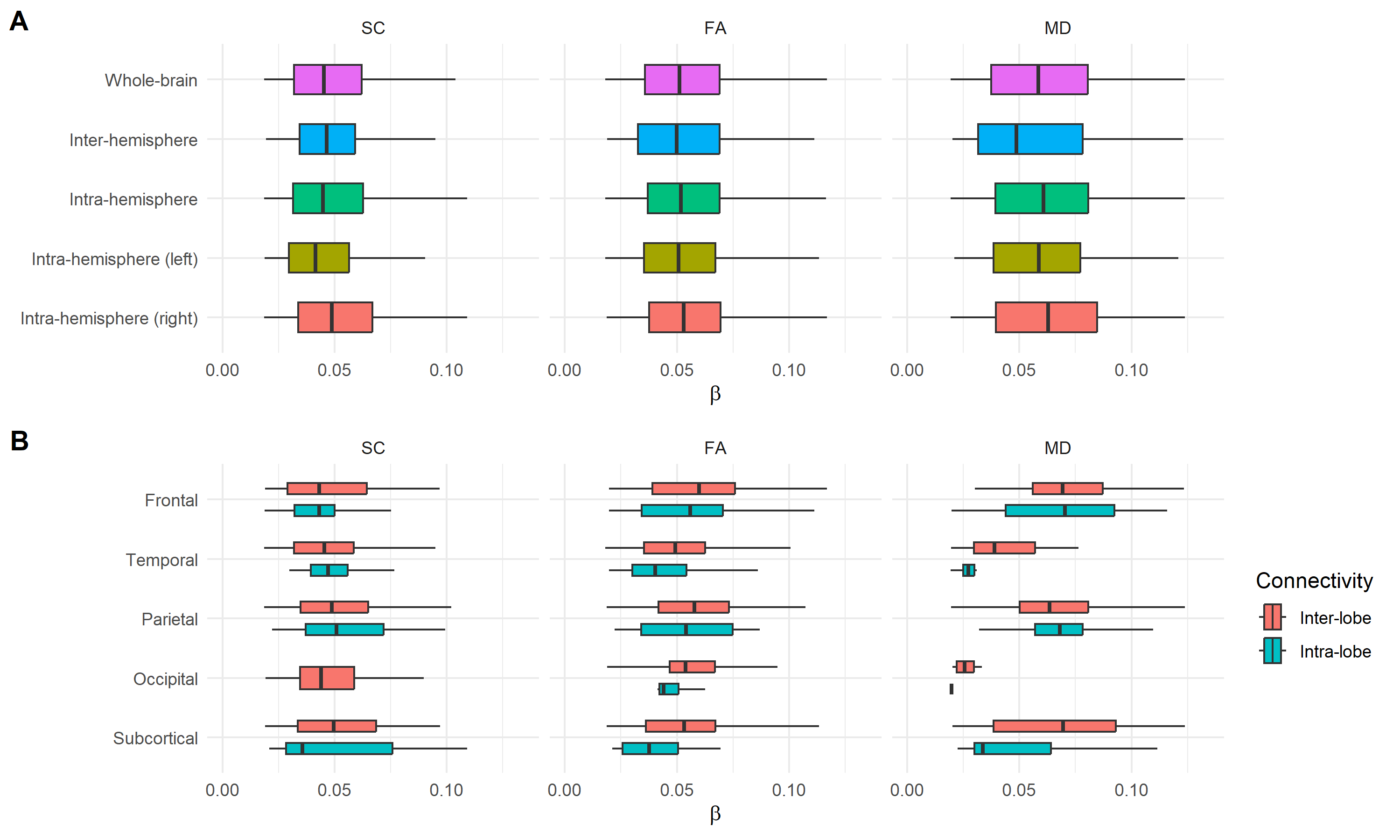

**Supplementary Fig. 5**. Boxplots of intra- and inter-module connectivity for significant edge-g associations (p < 0.05, FDR), derived from separate meta-analyses of SC, FA, and MD networks. Modules include: A) Whole-brain (all connections), inter-hemisphere (bilateral), intra-hemisphere (ipsilateral), and intra-hemisphere (left or right only) connections; B) Intra-lobe and inter-lobe (connections to any brain region located outside the designated lobe) for five brain lobes/regions. Insula and cingulate regions were excluded due to an insufficient number of edges. MD βs were sign flipped for plotting.

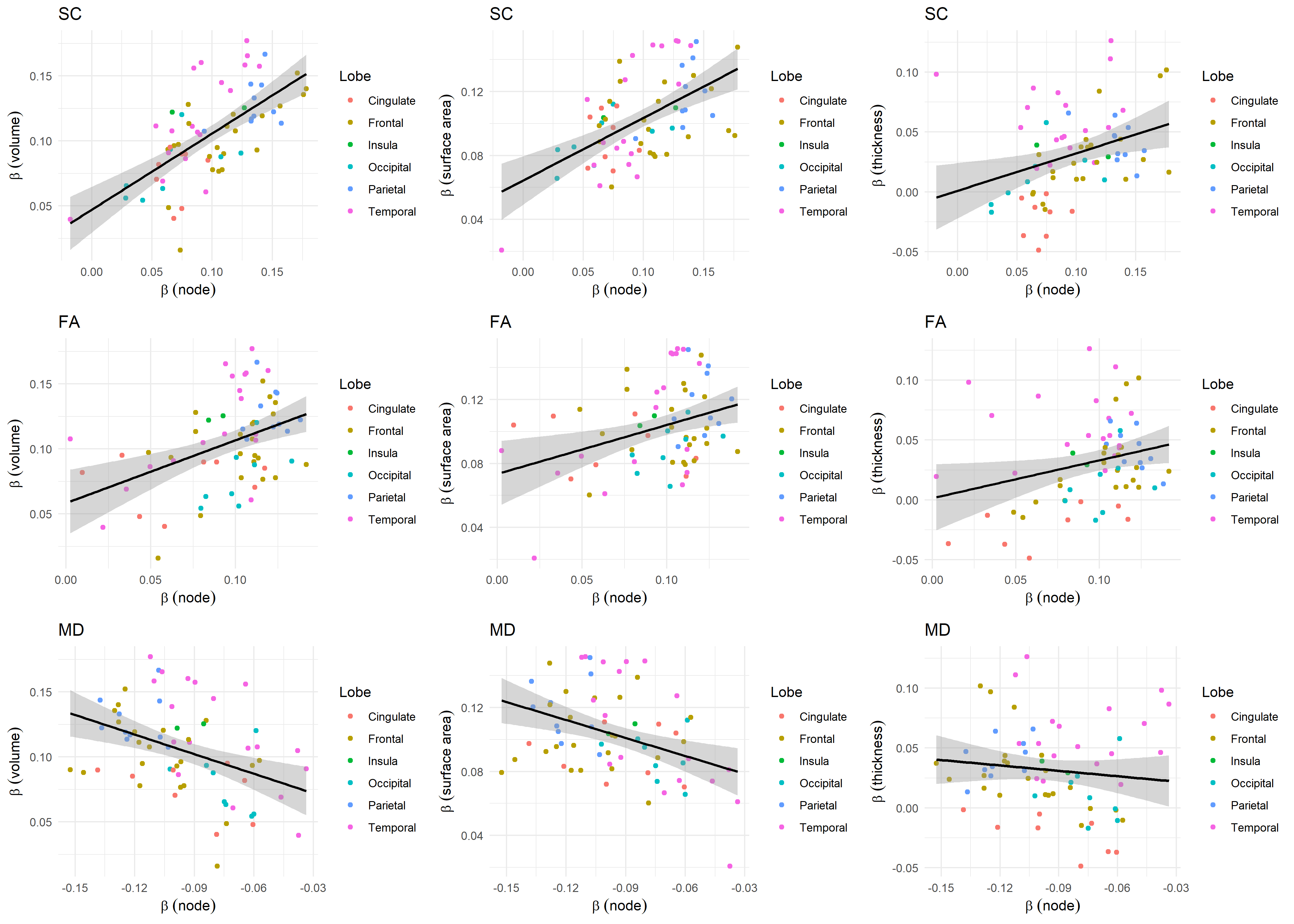

**Supplementary Fig. 6**: Scatter plots of the relationship between meta-analytic node-g β coefficients (derived from SC, FA, and MD networks) and the corresponding cortical morphometry-g β coefficients (volume, surface area, and thickness) across 68 cortical regions from the Desikan–Killiany atlas. Points are coloured by lobe, and black lines show linear fits.

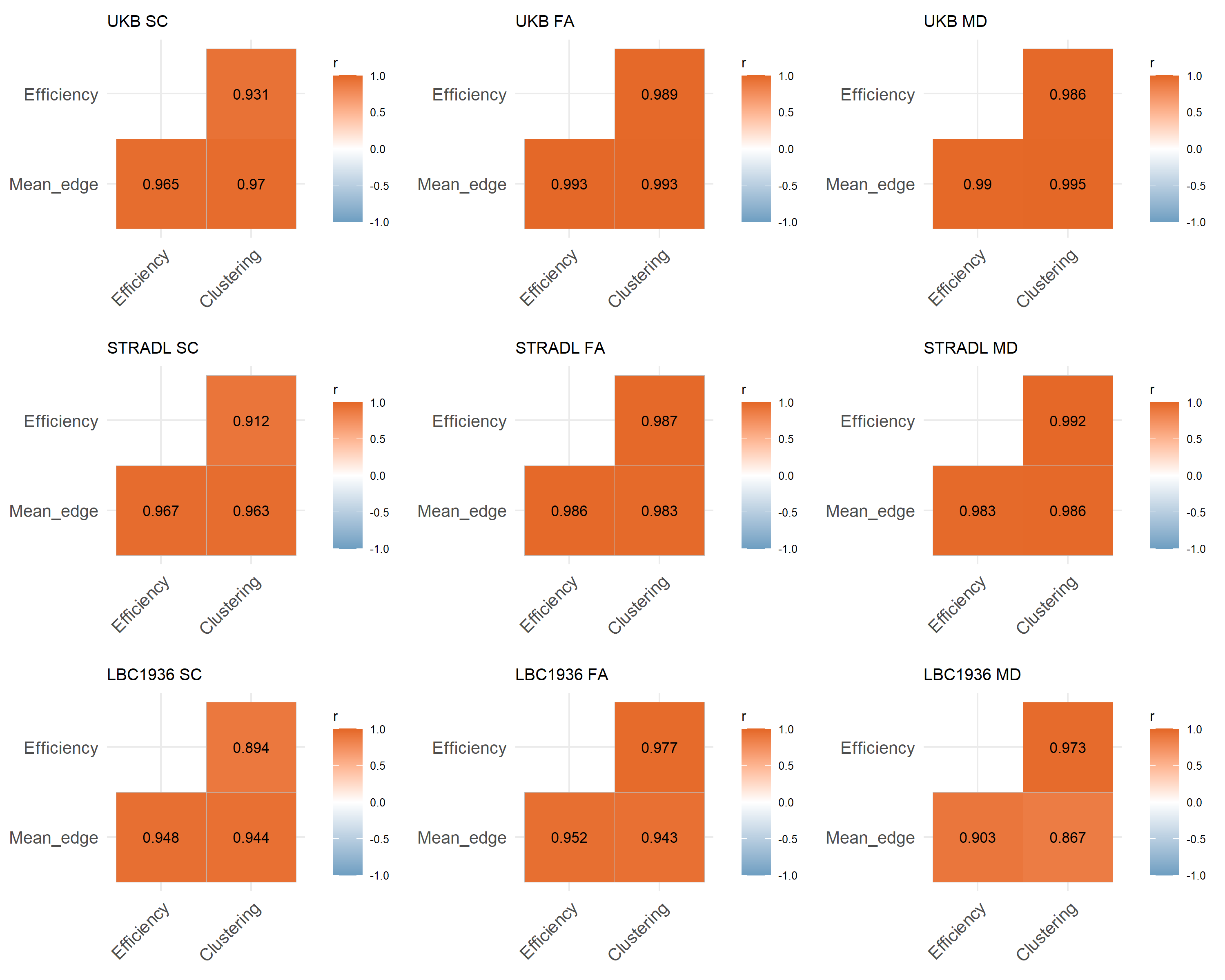

**Supplementary Fig. 7**. Correlations (Pearson’s *r*) among three network metrics (mean edge weight, global efficiency and network clustering coefficient) measured for each cohort and weighting scheme (SC, FA and MD). All correlations were significant (*p* < 0.001, uncorrected).
